# Stereoselective Covalent Targeting of BTK(C481S) and Kinases with β-Lactone Electrophiles

**DOI:** 10.64898/2026.07.03.736436

**Authors:** Celine D. Wang, Polina E. Barzova, Julian Robles, Ethan S. Toriki, Francisco J. Garcia, Jeffrey M. McKenna, Markus Schirle, Ziyang Zhang

## Abstract

The cysteine to serine mutation at residue 481 of Bruton’s tyrosine kinase (BTK) is the most common mechanism of clinical resistance against ibrutinib for the treatment of mantle cell lymphoma and chronic lymphocytic leukemia. We report small molecule ligands containing chiral β-lactone electrophiles to address this challenge. The asymmetric warhead enabled stereoselective covalent modification of wild-type and ibrutinib-resistant mutant BTK(C481S) through distinct sites of reactivity. Building on these findings, we developed kinase-directed β-lactone probes and demonstrated that individual enantiomers preferentially engage distinct subsets of the kinome. These studies establish β-lactones as stereochemically encodable covalent warheads whose stereochemistry can serve as a selectivity filter in covalent drug discovery.

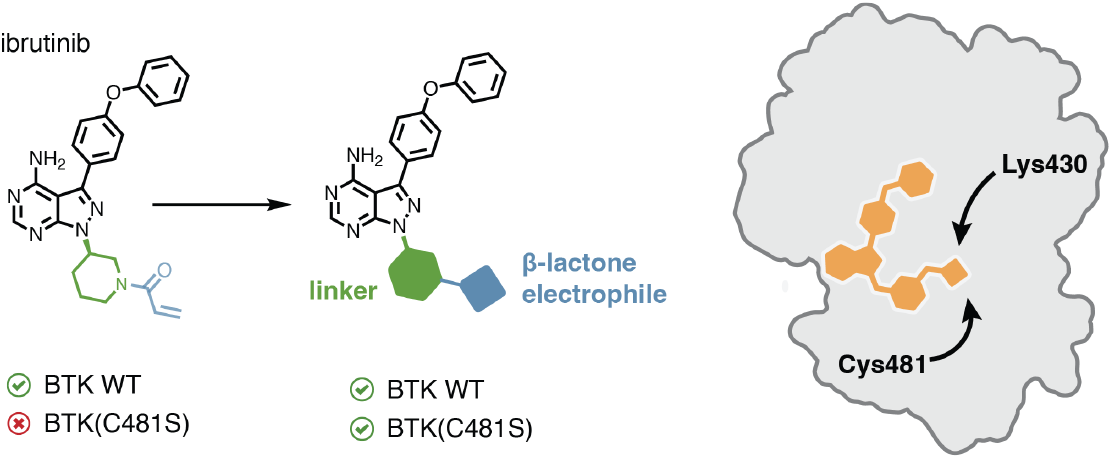

## Introduction

Targeted covalent ligands serve as a powerful therapeutic approach for the specific inhibition of oncogenic proteins. These molecules combine chemically encoded selectivity with reactivity-driven potency and increased residence time.^1,2^ Despite these advantages, the majority of currently used covalent drugs heavily rely on the presence of a strongly nucleophilic cysteine residue. Missense mutations of these cysteines often result in drastic reduction in the therapeutic efficacy of these covalent inhibitors and drive clinical resistance.

For example, ibrutinib, a first-in-class irreversible inhibitor targeting Bruton’s tyrosine kinase (BTK) achieves potent and selective BTK inhibition through a covalent bond between its acrylamide and a non-conserved cysteine.^3–5^ Clinical resistance takes the form of a C481S mutation in over 90% of affected patients.^6–8^ Such a mutation removes the mechanistically critical cysteine and renders BTK drastically less sensitive to ibrutinib inhibition, resulting in relapse of cancer in patients who were initially responsive.^6,9^ Most importantly, this mechanism of resistance has also been observed for second generation BTK inhibitors (e.g. acalabrutinib, zanubrutinib).^10,11^ The cysteine-to-serine mutation is not unique to BTK – clinical resistance against osimertinib, an inhibitor of epidermal growth factor receptor (EGFR), is frequently driven by the C797S mutation.^12,13^

Existing strategies to target these resistant mutants have focused on non-covalent approaches, often requiring redesigning the chemical scaffold^14–18^ or developing alternative modalities such as degraders.^19^ Since the acquired serines can still act as weak nucleophiles, we envisioned that a tailored electrophile would provide a complementary approach to overcoming resistance that preserves the favorable pharmacological features of covalent inhibition.

One class of such privileged electrophiles are β-lactones, which are highly strained (∼22.7 kcal/mol)^20^ electrophiles and have previously been exploited by both natural products and synthetic compounds to target catalytic and noncatalytic serines, threonines, or glutamic/aspartic acids.^21–28^ In addition, the two sp3 carbons in the 4-membered ring enable the construction of a chiral electrophilic environment which shapes the trajectory for the nucleophilic attack, a feature that has not been extensively studied.

Herein, we report the identification of chiral β-lactone electrophiles that covalently ligate both BTK(WT) and BTK(C481S). Careful exploration of the electrophile design revealed the crucial influence of the absolute configuration of the warhead and led to stereoselective ligands. High-resolution X-ray crystal structures uncovered two distinct modes of reactivity of the same compound against the wild-type and the mutant protein. This finding informed the design of β-lactone probes to profile stereoselective target engagement across the kinome.

## Results and Discussion

We hypothesized that replacing the cysteine-reactive acrylamide on ibrutinib with a β-lactone may confer reactivity against drug-resistant BTK(C481S). To test this hypothesis, we synthesized a pair of β-lactone-containing ibrutinib analogs, **1a** and **1b**, depicted in Figure 1A. These compounds were incubated together with recombinant BTK at ambient temperature, and samples were subjected to intact protein mass spectrometry (MS). Over a 24-hour treatment, we observed rapid and full modification of wild-type BTK by both compounds. We were also encouraged to detect 10% modification on BTK(C481S) by **1b**. At the same time, ibrutinib showed no modification, indicating the possibility of covalent engagement of BTK(C481S) by β-lactone ligands. Furthermore, diastereoisomer **1a**, which employs the enantiomer of the warhead, showed no modification of mutant BTK in this time frame (Figure 1B). This stereoselective reactivity motivated further investigation of the β -lactone probes.

**Figure 1.**
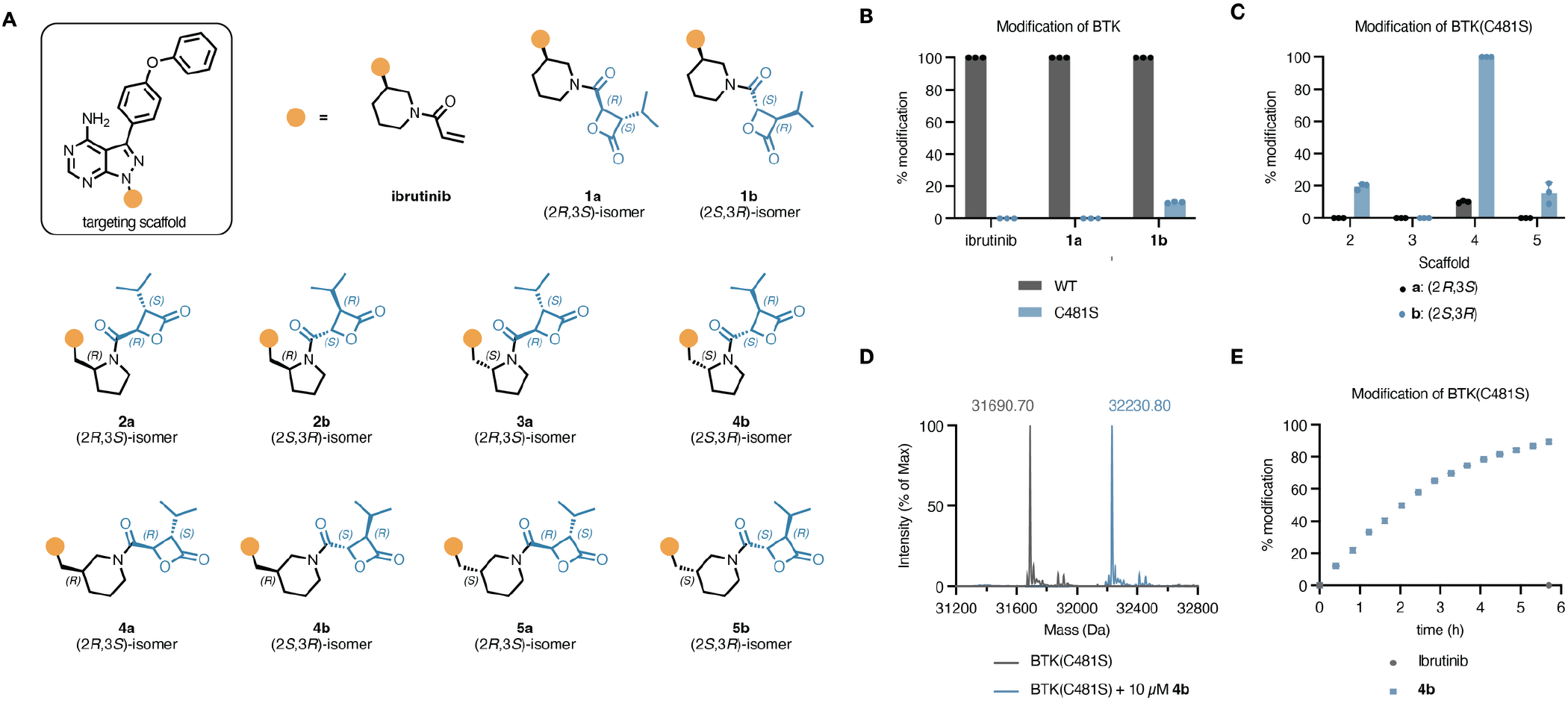
(A) Chemical structures of ibrutinib and β-lactone-containing analogs. (B) Intact MS analysis of covalent modification with BTK or BTK(C481S) kinase domain (KD) with ibrutinib analogs **1a,b.** Compounds (10 µM) were incubated with recombinant BTK(WT) or BTK(C481S) KD (1 µM) for 24 h, followed by MS analysis. (C) Intact MS analysis of covalent modification of BTK(C481S) KD by ibrutinib analogs **2-5**. Incubation reduced to 12 h (D) Intact MS analysis of BTK(C481S) kinase domain overlaid with MS analysis of BTK(C481S)•**4b** adduct. (E) Time-course intact MS analysis of modification of BTK(C481S) KD by **4b**.

Previous studies have shown that the reactivity of β-lactones is highly sensitive to the positioning of the electrophile.^24,25,29^ We therefore sought to optimize the molecules for better target engagement. To this end, a series of compounds (**2**–**5**) with varying length and geometry of the linker between pyrazolopyrimidine scaffold and β-lactone electrophile was designed and synthesized (Figure 1A). Next, the obtained ligands were assessed for their ability to covalently modify BTK(C481S) (Figure 1C).

Substitution of the piperidine ring linker on **1** with a 2-methylenylpyrrolidine in **2** led to a slight increase in covalent modification to 20% for compound **2b** containing a (2*S*,3*R*) configured β-lactone. By contrast, no modification was observed for the (2*R*,3*S*)-configured diastereomer **2a** or compounds **3a,b** in which the alpha stereocenter of the pyrrolidine is inverted (Figure 1C). We observed greatly improved labeling with compounds **4a,b** which contain a 2-methylenylpiperidine linker – one methylene unit longer than that in **1a,b**. Compound **4b** fully modified BTK(C481S) over 24 h, with 50% labeling observed after 2 h (Figure 1D,E). The diastereomer **4a** was less effective, resulting in 10% modification over 24 h. As observed for compounds **3a,b**, inversion of the alpha stereocenter at the piperidine ring also led to a loss in labeling, as seen for **5a,b** (Figure 1D).

With our optimized design **4b** in hand, we turned our attention to investigate the molecular basis of stereochemical recognition. We solved 1.8-Å and 1.2-Å resolution co-crystal structures of BTK(WT) and BTK(C481S), respectively, bound to **4b** (Figure 2A,B). In both structures, **4b** occupies the same ATP-competitive binding pocket as ibrutinib, with the pyrazolylpyrimidine group making the same polar contacts with the hinge region on BTK. For the wild-type protein, the β-lactone ring on **4b** is opened by Cys481 through nucleophilic attack of the thiol at the carbonyl. This gives rise to a thioester product, as confirmed by the continuous electron density (Figure 2A).

**Figure 2.**
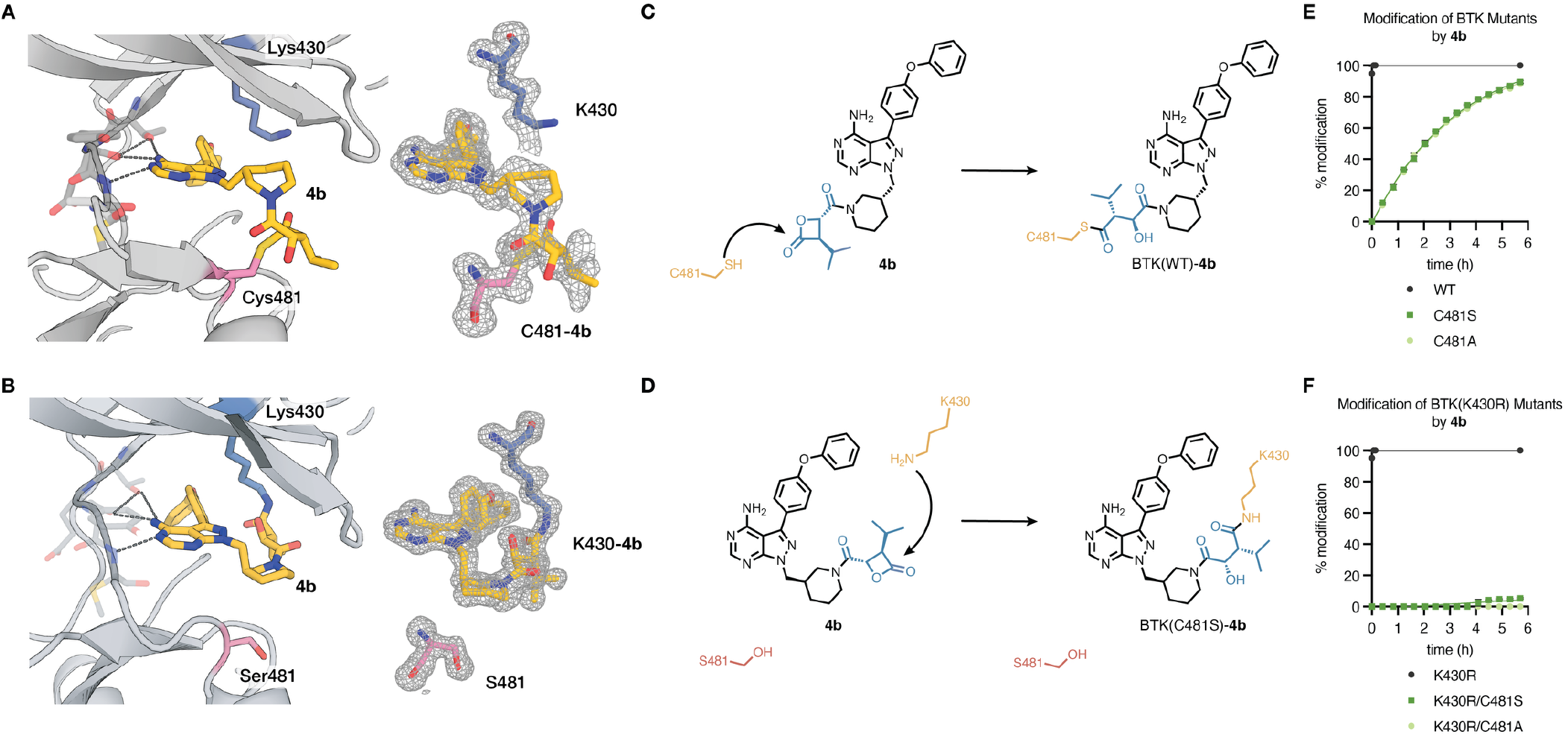
(A) Crystal structure of BTK(WT)•**4b** adduct. The covalent adduct is shown with an mF_o_–DF_c_ polder omit map in grey mesh (σ = 3.0), calculated by omitting ligand **4b**, Lys430, and Cys481. (B) Crystal structure of BTK(C481S)•**4b** adduct. (C,D) Schemes depicting the (C) reaction between compound **4b** and the Cys481 residue in BTK(WT) and (D) reaction between compound **4b** and Lys430 residue in BTK(C481S). (E) Time-course intact protein MS analysis of adduct formation by compound **4b** and recombinant BTK(WT), BTK(C481S), or BTK(C481A) KD. (F) Time-course intact protein MS analysis of adduct formation by compound **4b** and recombinant BTK(K430R) KD mutants.

Our finding suggests that **4b** can adopt distinct conformations to react with different nucleophilic residues in the ATP-binding pocket of BTK. In the wild-type protein, **4b** preferentially engages C481 due to the intrinsic high nucleophilicity of cysteines. If C481 is lost through mutation, K430 can be modified instead.

To understand the context-dependent chemoselectivity of the β-lactone electrophile, we measured the rate of adduct formation between different BTK mutants and **4b** through intact MS. In the presence of Cys481, as with wildtype or BTK(K430R) mutant, labeling was rapid and complete within 6 min. By contrast, labeling of BTK(C481S) and BTK(C481A) occurred slower but at comparable rates. This suggests that reactivity dominantly occurs through Lys430 (Figure 2E). We asked whether reaction with mutant serine (S481) is possible by preparing the double mutant BTK(K430R/C481S). We observed low but quantifiable levels of adduct formation (5%) in under 6 hours. Despite the low reactivity, we confirmed that the reaction was specific to S481, because we observed no covalent modification with the BTK(K430R/C481A) double mutant (Figure 2F). Thus, our study revealed that the β-lactone can serve as a versatile electrophile. In the ATP pocket of BTK, the rate of reactivity is consistent with covalent bond formation with the most reactive, available nucleophile, with the order of preference: Cys481, Lys430, Ser481.

To assess whether covalent adduct formation occurs in cells, we generated the alkyne-containing probe analogs of **4b** and its enantiomer **5a**, as **6** and **7**, respectively (Figure 3A).^30^ Ramos cells, a B-cell lymphoma cell line that expresses wild-type BTK, were treated with probe **6** or **7**, lysed and the lysate was subjected to copper-catalyzed azide-alkyne cycloaddition (CuAAC) with TAMRA-N_3_. Proteins were separated by SDS-PAGE and protein labeling was visualized by in-gel fluorescence scanning. We observed that probe analogs **6** and **7** both labeled wild-type BTK after a 1-h treatment in live cells (Figure S2), suggesting that the thioester adducts of beta lactone and C481 are stable in the context of a cellular proteome.

**Figure 3.**
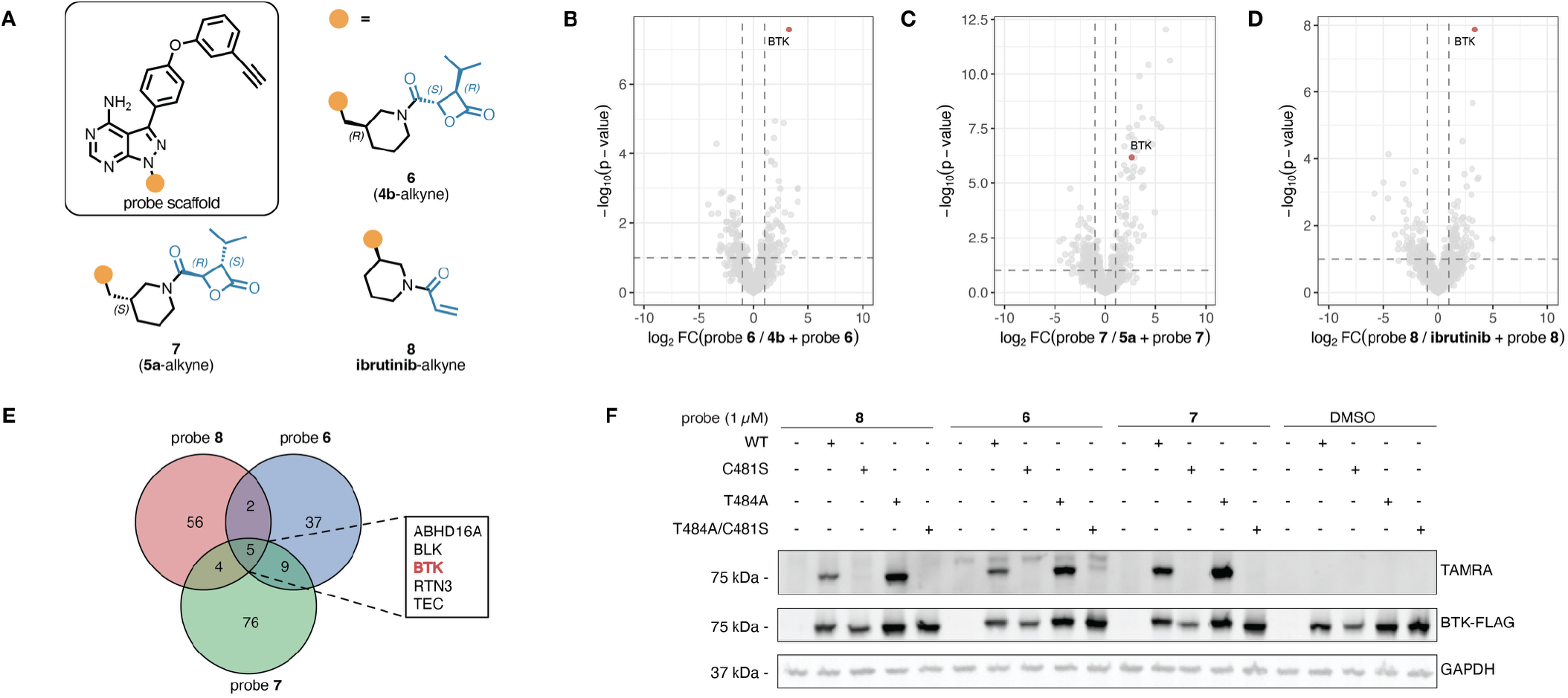
(A) Chemical structures of alkyne-containing probe analogs of **4b, 5a**, and ibrutinib. (B) Proteomic profiling of on-target proteins in Ramos cells. Cells were first treated by **4b** (20 µM) and then labeled by probe **6** (1 µM). Labeled proteins were subject to CuAAC and streptavidin pulldown. Enriched proteins were analyzed by label-free DIA quantitative proteomics by LC-MS/MS (C) Proteomic profiling in Ramos cells, first treated by **5a** (20 µM) and then labeled by probe **7** (1 µM). (D) Proteomic profiling in Ramos cells, first treated by ibrutinib (20 µM) and then labeled by probe **8** (1 µM). (E) Comparison of significantly competed proteins by probes **6-8.** Significant proteins defined as having a fold-change greater than 2 and *p-*value less than 0.05. (F) Labeling of overexpressed BTK mutants by probes **6** – **8** in HEK293T cells transfected with FLAG-tagged BTK mutants. Cells were treated with probes **6** – **8** (1 µM) for 4 h.

Next, we performed competition-based cysteine profiling proteomics by treating live Ramos cells with compound **4b, 5a**, or ibrutinib followed by cell lysis and treatment with desthiobiotin iodoacetamide. After tryptic digestion, probe-modified peptides were enriched for and quantified by label-free DIA quantitative MS. This workflow allows us to directly identify and quantify cellular target engagement with residue-level information. The site of reactivity was mapped to Cys481, consistent with the biochemical observations for wild-type BTK (Figure S3).

We then assessed the selectivity of these ligands by competition-based chemoproteomics with matched ligand-probe pairs, which allows for the identification of stoichiometric targets.^30^ Ramos cells were first treated with either compound **4b, 5a**, or ibrutinib, followed by the matched alkyne probe **6, 7**, or **8**, respectively. The labeled lysate was then subjected to CuAAC with biotin-azide for streptavidin enrichment, and protein targets were identified and quantified by label-free DIA quantitative MS. Encouragingly, the most significantly competed stoichiometric target of **4b** was BTK (Figure 3B), suggesting that this ligand is highly selective for BTK in cells. In comparison, the diastereomer, **5a**, and ibrutinib engaged several targets in addition to BTK (Figure 3C,D). Only five targets were selectively engaged by all three compounds including BTK as well as previously identified off-target kinases for ibrutinib, BLK and TEC, both of which contain a cysteine at the C481 equivalent position (Figure 3E).^30^ Probe **6** was also more selective for BTK compared to diastereomer **7** and ibrutinib probe **8** (Figure 3E).

Next, we aimed to assess the modification of drug-resistant mutant BTK(C481S) in the context of a cellular proteome. Under identical treatment conditions as before, we did not observe any BTK(C481S) labeling over a 24-hour incubation in HEK293T over-expressing BTK mutants (Figure S4). We hypothesized that the lack of labeling may be a combined effect from slower reaction kinetics and the competition with endogenous ATP in cells. To test this, we introduced the BTK(T474A) gatekeeper residue mutants in HEK293T cells. The T474A mutation has been previously shown to decrease BTK enzymatic activity while maintaining sensitivity to ATP-site competitive inhibitors^9,31^ by reducing K_m_(ATP).^32,33^ When these cells were treated with our probe analog **6**, we observe low but consistent labeling of the BTK(T474A/C481S) mutant. This labeling was not observed with BTK(C481S), or when the enantiomeric probe **7** was used (Figure 3F). We also observed this labeling in a dose-dependent manner (Figure S5A) that saturates within 2 h (Figure S5B). While our current compounds did not achieve 100% engagement of mutant BTK in cells, our data suggest that mutant BTK is ligandable by designed electrophilic compounds in a cellular context. Improvements on their reversible binding affinity as well electrophile positioning will likely enhance their cellular potency.

Our investigation on BTK shows three unique properties of β-lactone electrophiles: 1) they can react with multiple nucleophilic residues, including cysteine, lysine, and serine; 2) meanwhile, they are not broadly reactive compounds and can achieve excellent selectivity in cells; 3) chiral substituents on the electrophile can encode target specificity. These features prompted us to ask whether the stereochemistry of these β-lactone electrophiles provides an orthogonal means to achieve selective targeting of homologous kinases.

We synthesized enantiomeric β-lactone-containing analogs based on a general kinase inhibitor scaffold as found in the lysine-reactive general kinase probe, XO44 (Figure 4A).^34^ To this end, the fluorosulfonyl benzyl in XO44 was substituted with beta-lactone amides to yield compounds **9a,b**. These compounds contain terminal alkyne groups, allowing conjugation of fluorophores or biotin by CuAAC. As a control, we labeled cells with XO44.^35^ We observed that the β-lactone electrophile provided differential labeling compared to the electrophilic sulfonyl fluoride on XO44. Furthermore, we observed differences between the two β-lactone probe enantiomers (Fig 4B). Both electrophile choice and stereochemistry allowed for the selective targeting of kinase across the proteome.

**Figure 4.**
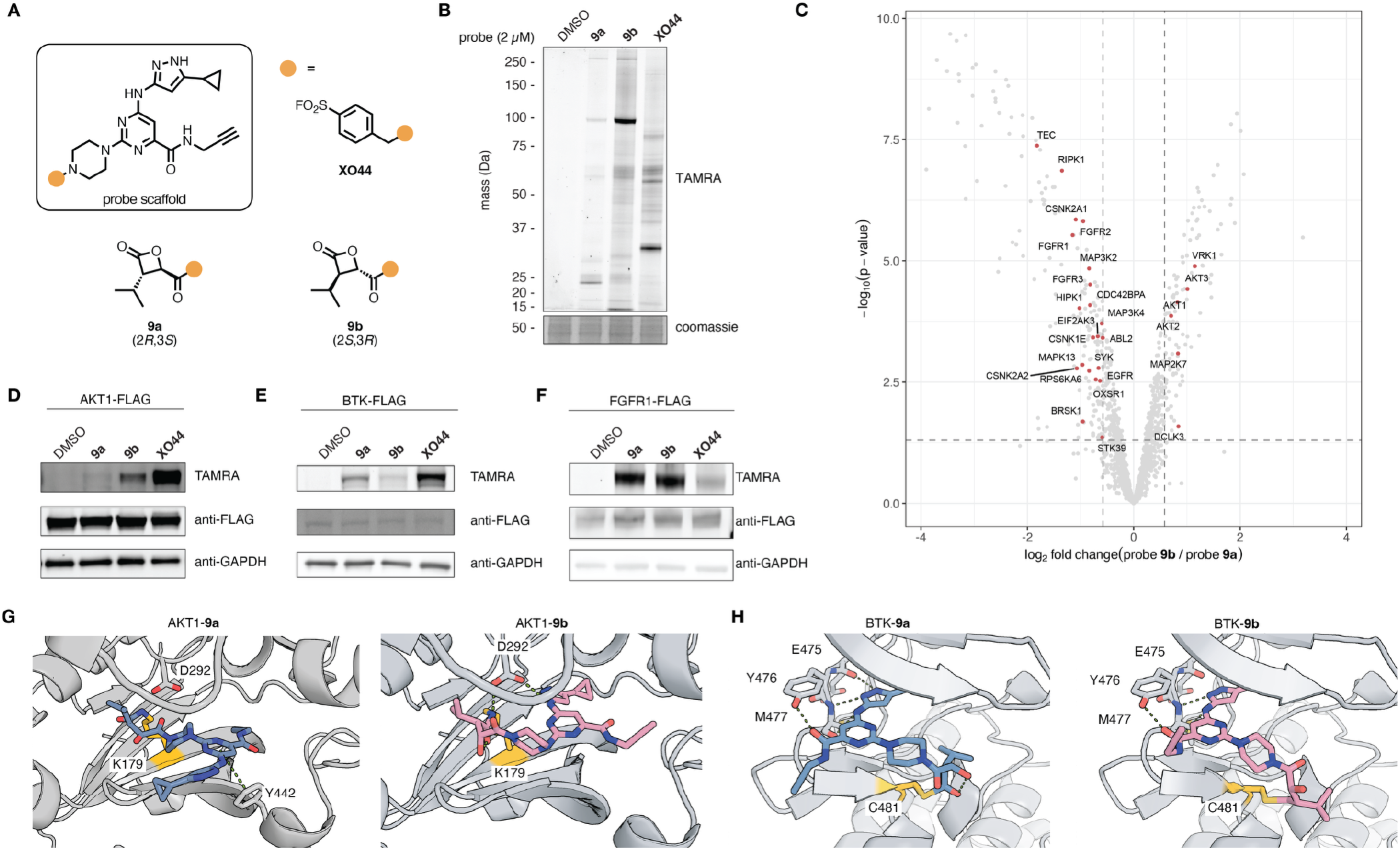
(A) Chemical structures of general kinase probes with sulfonyl fluoride (XO44) or β-lactone (**9a,b**) electrophiles. (B) Labeling of proteins by general kinase probes in live cells. HEK293T cells were treated with kinase probe **XO44, 9a**, or **9b** for 1 h. Cell lysates were subject to CuAAC with TAMRA-azide, resolved by SDS-PAGE, and scanned for fluorescence. (C) Proteomic profiling of cellular targets of probe **9** versus probe **10** in HEK293T cells. Cells were treated by probes (2 µM) for 1 h, after which cell lysates were subject to CuAAC with biotin-PEG3-azide and enriched by streptavidin pulldown. Enriched proteins were analyzed by TMT-based quantitative proteomics by LC-MS/MS. (D) – (F) Labeling of overexpressed (D) AKT, (E) BTK, (F) and FGFR by general kinase probes XO44, **9a**, and **9b** in HEK293T cells. Cells were treated by probes (2 µM) for 1 h. Cell lysates were subject to CuAAC with TAMRA-azide, resolved by SDS-PAGE, and scanned for fluorescence. (G,H) Predicted binding pose of probes **9a** (blue) and **9b** (pink) upon covalent docking with (G) Lys179 (yellow) in AKT1 and (H) Cys481 (yellow) in BTK. Key polar or pi-interactions shown in green.

To characterize the different targets, we treated HEK293T cells with our probes, enriched labeled proteins by biotin-streptavidin pulldown, and identified and quantified protein targets by tandem mass tag (TMT)-based quantitative mass spectrometry (Figure 4C). Consistent with the in-gel labeling, we observed targets selectively enriched by each of the three probes. Several kinases demonstrated preferential reactivity with the β-lactone electrophiles over the sulfonyl fluoride. Additionally, multiple kinases were enriched by one β-lactone enantiomer over the other (Figure S6).

Lastly, to explore the origins of the observed stereoselective engagement of AKT1 and BTK, we generated structural models of the putative covalent adducts through covalent docking Schrodinger Maestro. Probes **9a,b** were docked into the ATP-binding site of AKT1 (from PDB 4GV1) as amide adducts with Lys179 (Figure 4G) and the ATP-binding site of BTK (from PDB 5P9J) as thioester adducts with Cys481 (Figure 4H). The resulting energy-minimized poses suggested preferable interactions between **9b** and AKT1 compared to its antipode **9a**, including possible hydrogen bonds with Asp292. Both probes adopted similar poses in BTK with subtle differences in polar contacts with the hinge residues. Although these models should be interpreted cautiously, they provide a possible structural rationale for the observed stereochemical preferences.

## Conclusions

We report a chemical strategy to address the clinically observed BTK(C481S) resistance mutation by replacing the cysteine-reactive electrophile on an existing targeted covalent inhibitor, ibrutinib, with a β-lactone electrophile. We identified a β-lactone **4b**, which covalently engages both wild-type BTK and drug-resistant mutant BTK(C481S) through different mechanisms. While **4b** reacts with Cys481 in wild-type BTK, it instead engages Lys430 in BTK(C481S). Remarkably, the same β-lactone scaffold can engage wild-type and resistant BTK through distinct nucleophilic residues, revealing a strategy for overcoming resistance mutations that eliminate the canonical covalent target and providing a path toward other clinically observed Cys481 resistance variants in patients treated with second-generation BTK inhibitors.^10,11,36^

This use of stereochemistry to achieve target selectivity has previously been demonstrated with chiral phosphorus(V)^37^ and sulfur(VI) electrophiles^38^, as well as cysteine electrophiles coupled to chiral fragments.^39–43^ We extend this strategy to β-lactone electrophiles. Using a promiscuous kinase inhibitor equipped with enantiomeric β-lactones, we showed that each antipode preferentially captured distinct subsets of the kinome. This divergence highlights the capacity of chiral electrophiles to differentiate between structurally similar binding pockets and leverage stereochemical recognition as a selectivity filter for highly homologous kinases.

One limitation of the current study is that our compounds show low cellular activity. Nevertheless, the BTK(C481S) co-crystal structure provides a blueprint for improving ligand conformation, electrophile positioning, and Lys430 engagement. Consistent with prior studies showing that BTK Lys430 is chemically addressable,^44,45^ our work establishes β-lactones as a complementary strategy for targeting this residue, adding a structurally characterized electrophilic scaffold to the growing toolbox of lysine-targeting covalent chemistry.^46–48^

Collectively, our work introduces stereochemically encoded β-lactone electrophiles capable of dual cysteine- and lysine-targeting and reveals new opportunities for selective kinase targeting. More broadly, these findings demonstrate how stereochemical control of covalent reactivity can be harnessed to overcome resistance mutations and expand the scope of covalent drug discovery.

## ASSOCIATED CONTENT

### Supporting Information

Supplementary Figures 1-7, Supplementary Tables 1-4, detailed materials and methods, 1H spectra, and supplementary references (PDF).

Supplementary Table 5: Cysteine profiling proteomics of compounds **4b, 5a**, and ibrutinib in Ramos cells (XLSX).

Supplementary Table 6: Chemoproteomic profiling of Ramos cells by probes **6-8**. (XLSX).

Supplementary Table 7: Chemoproteomic profiling of HEK293T cells by probes **9a,b** and XO44. (XLSX).

### Accession Codes

Atomic coordinates and structure factors for the crystal structure of BTK(WT)•**4b** and BTK(C481S)•**4b** adducts have been deposited with the Protein Data Bank (PDB) with the accession numbers 12LE and 12LF, respectively.

## AUTHOR INFORMATION

### Author Contributions

CDW, ZZ conceived the project, designed the study, and wrote the manuscript. CDW, PEB, JR, EST, FJG performed experiments for the paper. JMM, MS provided critical insights into project experiments. The manuscript was written through contributions of all authors. All authors have read and approved the final version of the manuscript.

## ACKNOWLEDGMENT

This work is supported by Novartis Institutes for Biomedical Research and the Novartis-Berkeley Translational Chemical Biology Institute (NB-TCBI) and NIH/NCI 1R21CA280163-01. We thank Jay Nix and the staff at the Advanced Light Source beamline 8.2.2 for help with X-ray data collection and processing. CDW was partially supported by a Chemical Biology Training Grant from the NIH (066698). We thank Drs. Hasan Celik, Raynald Giovine, and Pines Magnetic Resonance Center’s Core NMR Facility (PMRC Core) for spectroscopic assistance. Instruments in the College of Chemistry NMR facility are partly supported by NIH S10OD024998. We thank A. Schepartz and members of the Zhang lab for providing comments and suggestions to this manuscript.

