## Supplemental Information for "Stereoselective Covalent Targeting of BTK(C481S) and Kinases with β-Lactone Electrophiles"

**Table of Contents**

I. Supplementary Figures and Tables
   a. Figure S1. Full structures of compounds.
   b. Figure S2. Labeling of proteins by probes **6 – 8** in Ramos cells.
   c. Figure S3. Cysteine profiling proteomics of Ramos cells with ibrutinib, compounds **4b,** and **5a**.
   d. Figure S4. Labeling of overexpressed BTK mutants in HEK293T cells.
   e. Figure S5. Dose-dependent and time-course labeling of BTK(T474A/C481S) mutants in HEK293T cells.
   f. Figure S6. Chemoproteomics of probes **9a,b** vs **XO44**.
   g. Figure S7. Predicted docking poses of probe **9a,b** versus AKT1.
   h. Table S1. X-ray crystallography data collection and refinement statistics.
   i. Table S2. List of proteins enriched by probes **6**, **7**, and **8** from competition proteomics.
   j. Table S3. Covalent docking scores and conformational energies of probes **9a,b** with AKT1 and BTK
   k. Table S4. List of antibodies.

II. Biological Methods

III. Chemical Methods

I. Supplementary Figures and Tables

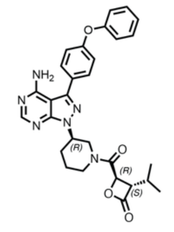

**1a**

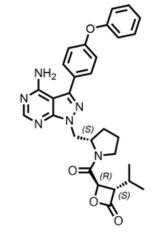

**2a**

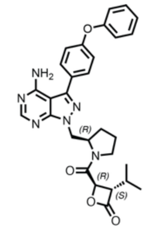

**3a**

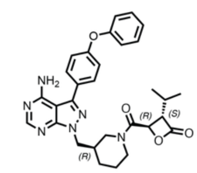

**4a**

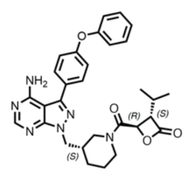

**5a**

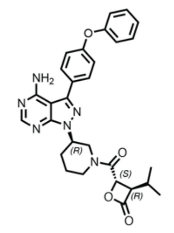

**1b**

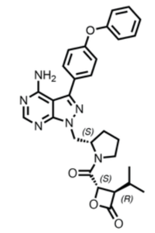

**2b**

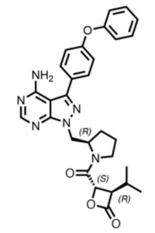

**3b**

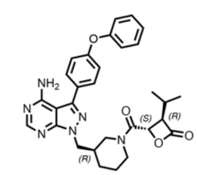

**4b**

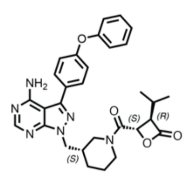

**5b**

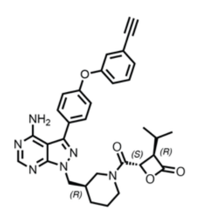

**6**
(**4b**-alkyne)

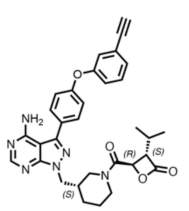

**7**
(**5a**-alkyne)

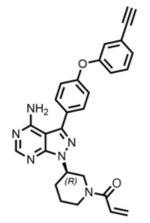

**8**
(ibrutinib-alkyne)

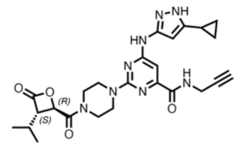

**9a**

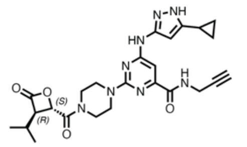

**9b**

**Figure S1**. Full structures of compounds used in this study.

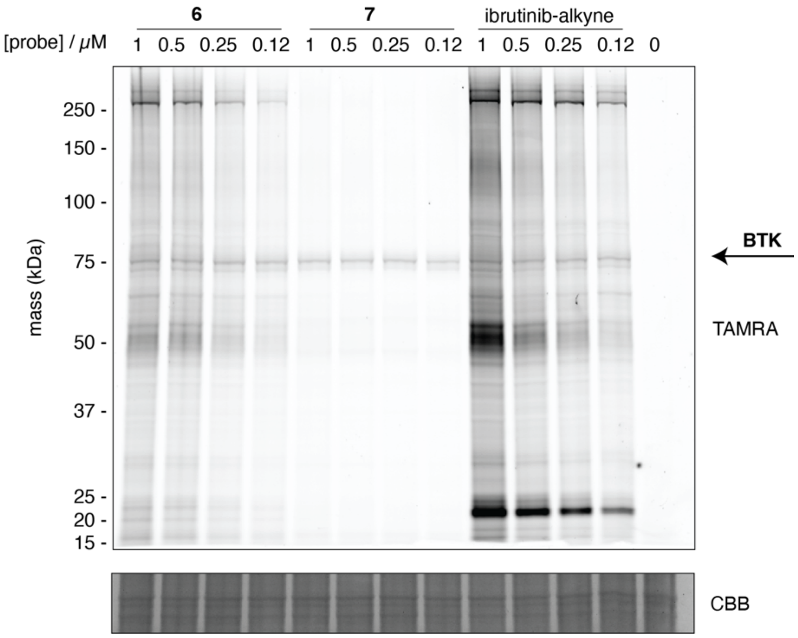

**Figure S2**. Labeling of proteins by probes **6 – 8** with varying concentrations in Ramos cells. Cells were treated with probes **6 – 8** for 1 h. Cell lysates were subject to CuAAC with TAMRA-azide, resolved by SDS-PAGE, and scanned for fluorescence.

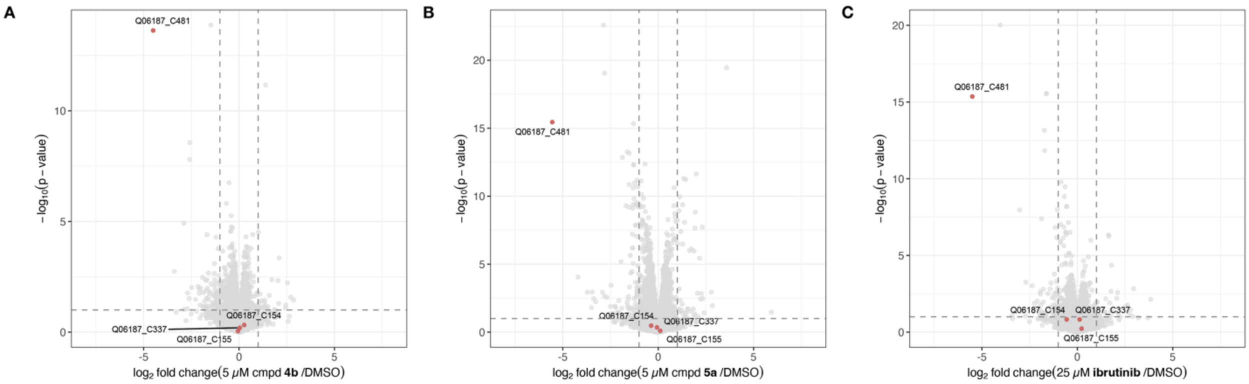

**Figure S3**. Cysteine profiling proteomics in Ramos cells. Cells were first treated by compound **4b** (5 μM), **5a** (5 μM), or **ibrutinib** (25 μM) and lysates incubated with iodoacetamide desthiobiotin. Labeled peptides were enriched and quantified by label-free DIA quantitative mass spectrometry. (n = 3 biologically independent samples).

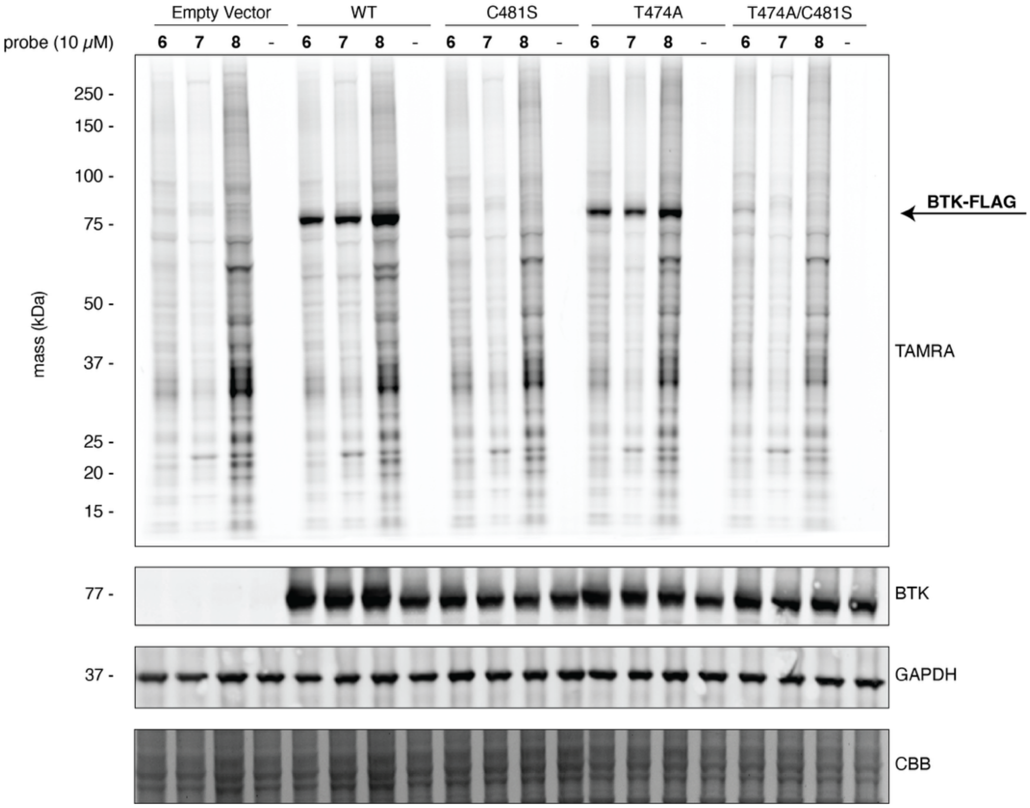

**Figure S4**. Labeling of proteins in HEK293T cells transfected with FLAG-tagged BTK mutants. Cells were treated with probes **6** – **8** (10 µM) or DMSO for 24 h. Cell lysates were subject to CuAAC with TAMRA-azide, resolved by SDS-PAGE, and scanned for fluorescence.

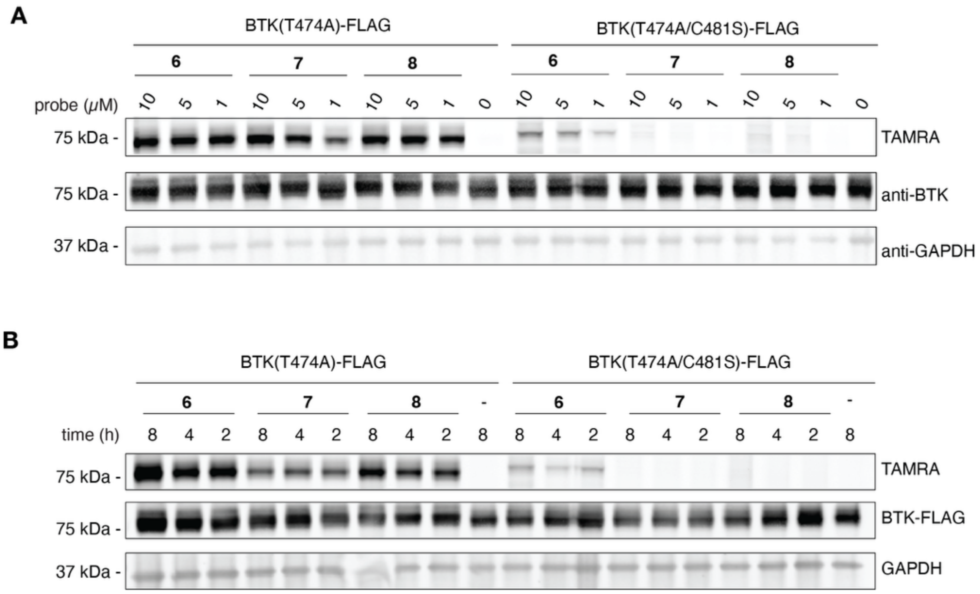

**Figure S5**. (A) Dose-dependent labeling of overexpressed BTK(T474A)-FLAG and BTK(T474A/C481S)-FLAG in HEK293T cells treated by probes **6** – **8** for 4 h. Cell lysates were subject to CuAAC with TAMRA-azide, resolved by SDS-PAGE, and scanned for fluorescence. (B) Time-course labeling of HEK293T cells transfected with BTK(T474A)-FLAG and BTK(T474A/C481S)-FLAG and treated by probes **6** – **8** (1 μM).

A

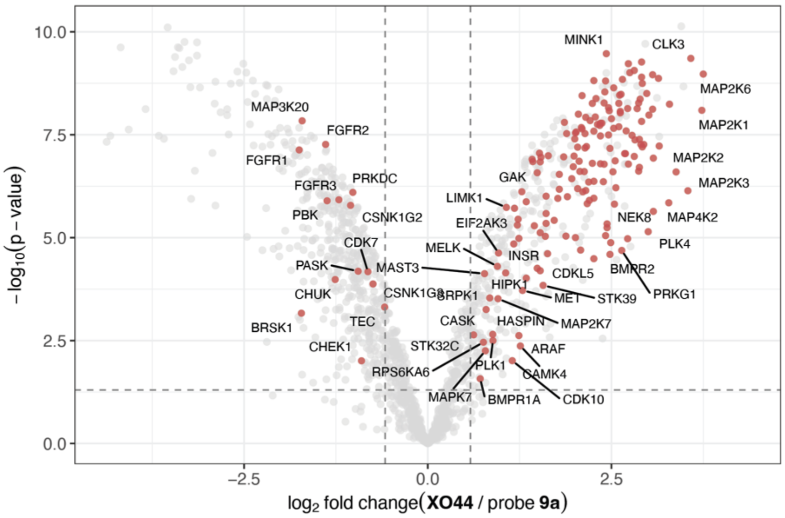

B

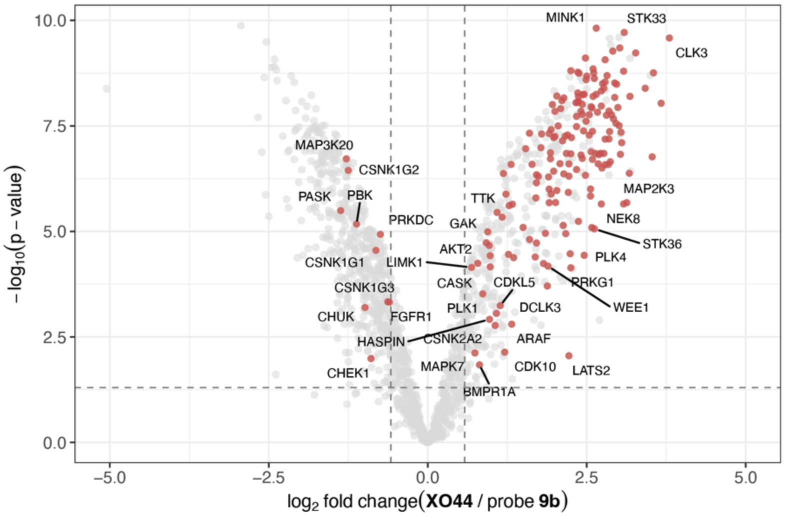

**Figure S6**. Proteomic profiling of cellular targets of (A) **XO44** versus probe **9** and (B) **XO44** probe **10** in HEK293T cells. Cells were treated by probes (2 μM) for 1 h, after which cell lysates were subject to CuAAC with biotin-PEG3-azide and enriched by streptavidin pulldown. Enriched proteins were analyzed by TMT-based quantitative proteomics by LC-MS/MS. Kinases are highlighted in red. (n = 3 biologically independent samples)

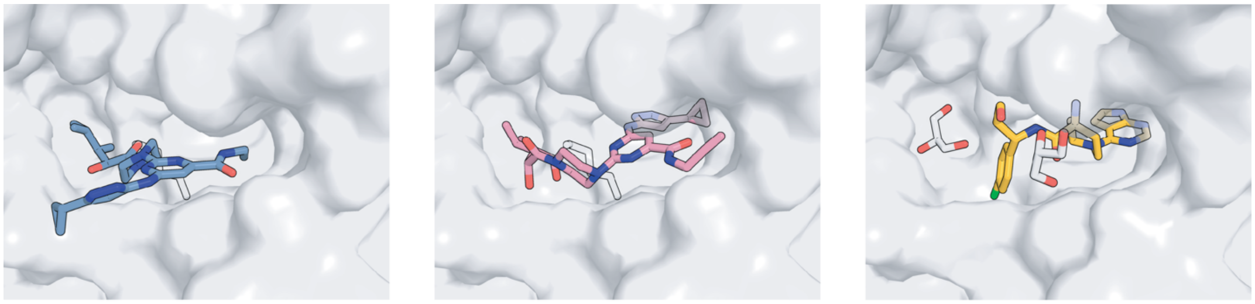

AKT1-**9a** AKT1-**9b** AKT1-**capivasertib**

**Figure S7**. Predicted binding poses of probes **9a** and **9b** from covalent docking with Lys(179) in AKT1, depicted as surface model, in comparison to capivasertib (PDB 4GV1).

**Table S1**. X-ray crystallography data collection and refinement statistics

| | **BTK(WT)•4b** | **BTK(C481S)•4b** |
|---|---|---|
| **Data collection** | | |
| Space group | $P\ 22_12_1$ | $P\ 2_12_12$ |
| Cell dimensions | | |
| *a*, *b*, *c* (Å) | 37.99 71.79 104.18 | 71.596 104.853 38.11 |
| α, β, γ (°) | 90 90 90 | 90 90 90 |
| Resolution (Å) | 59.11 - 1.77 (1.83 - 1.77) | 42.3 - 1.23 (1.24 - 1.23) |
| $R_{merge}$ | 0.0529 (0.1349) | 0.1873 (0.4572) |
| $I / \sigma I$ | 14.10 (7.20) | 4.83 (0.59) |
| Completeness (%) | 98.63 (99.40) | 93.97 (36.46) |
| Redundancy | 3.2 (3.2) | 6.3 (1.4) |
| **Refinement** | | |
| Resolution (Å) | | |
| No. reflections | 28200 (2798) | 79064 (1026) |
| $R_{work}$ | 0.1636 (0.1461) | 0.1825 (0.2991) |
| $R_{free}$ | 0.1954 (0.2123) | 0.2040 (0.2959) |
| No. atoms | | |
| Protein | 2223 | 2271 |
| Ligand/ion | 40 | 40 |
| Water | 363 | 363 |
| *B*-factors | | |
| Protein | 12.17 | 14.92 |
| Ligand/ion | 22.87 | 11.07 |
| Water | | |
| R.m.s. deviations | | |
| Bond lengths (Å) | 0.006 | 0.005 |
| Bond angles (°) | 0.83 | 0.86 |

*Values in parentheses are for highest-resolution shell.

**Table S2.** List of proteins enriched by probes **6**, **7**, and **8**. Significant proteins defined as having a fold-change greater than 2 and p-value less than 0.05.

*Overlapping targets*

| 6, 7, and 8 | 6 and 7 | 6 and 8 | 7 and 8 |
|---|---|---|---|
| RTN3 | CDIPT | TPM4 | LYPLA1 |
| ABHD16A | MT-ND5 | GUF1 | PSMB5 |
| TEC | PFKL | | C18orf32 |
| BLK | HMOX2 | | CTSZ |
| BTK | SELENOT | | |
| RTN3 | PAFAH2 | | |
| | ABHD6 | | |
| | PPT2 | | |
| | SELENOK | | |

*Probe specific targets*

| 6 only | 7 only | | 8 only | |
|---|---|---|---|---|
| PWWP4 | DEGS1 | MTDH | PDXK | INPP5B |
| GMDS | GNPAT | GPAT4 | MAP2K7 | CSTF2 |
| FLOT1 | CYB5B | COMTD1 | PMM2 | MAN1A1 |
| DDHD2 | SLC25A20 | CEND1 | CUTA | UBE2G1 |
| SUN1 | LY75 | TMEM256 | COL1A1 | GOLGA2 |
| HLA-B | LYPLA2 | PLA2G15 | APOA1 | AKAP13 |
| CAPNS1 | CD74 | ABHD11 | EPHX1 | UAP1 |
| ATP5MC1 | ASNS | DDX55 | COX7A2 | MAPK14 |
| GRK4 | CD37 | PIP4K2C | CD53 | HIBCH |
| IDH3G | CKB | ABHD3 | MPV17 | MARK2 |
| DHODH | CTSS | NUDT8 | MAPK9 | UBR7 |
| PRKAA1 | PSMB8 | TAF15 | COPS2 | ATAD1 |
| SCRIB | PSMB9 | ALG14 | BST2 | OTULIN |
| UAP1L1 | TSPO | ABHD17A | TMEM41B | DOCK10 |
| MRM1 | SOAT1 | ABHD14B | RETSAT | RANBP9 |
| OSBPL9 | NUDT1 | TMEM68 | NUP54 | DPH5 |
| PAFAH2 | PNPLA4 | PSMB7 | ABCA12 | PANK4 |
| TIMM29 | CPT1A | TXN2 | TMEM123 | ZC3H7B |
| BCL2L13 | ALDH3A2 | SELENOS | PRXL2B | |
| EHMT1 | LIMK1 | C17orf80 | PIGK | |
| PDLIM7 | PIP4K2B | DESI2 | APOBEC3D | |
| UFC1 | SLC35A1 | ALG1 | PAWR | |
| NCOA3 | REEP5 | MIS12 | CNDP2 | |

| | | | |
|---|---|---|---|
| OAS1 | PFKP | TMEM245 | VKORC1 |
| ITGA4 | TAP1 | FOXP1 | PRXL2A |
| HIKESHI | TMED1 | SCPEP1 | TMEM109 |

**Table S3.** Covalent docking and energy scores for compounds **9a,b** against AKT1 and BTK.

| Title | OPLS Version | cdock Affinity Score | Post-reactive Docking Score | Pre-reactive Docking Score | Relative Potential Energy (S-OPLS) | Prime Energy |
|---|---|---|---|---|---|---|
| AKT1-**9a** | OPLS4 | -2.68 | -2.68 | -4.176 | 22.058 | -13654.3 |
| AKT1-**9b** | OPLS4 | -4.304 | -4.304 | -4.93 | | -13677.09 |
| BTK-**9a** | OPLS4 | -9.11 | -9.11 | -8.313 | | -11116.43 |
| BTK-**9b** | OPLS4 | -8.439 | -8.439 | -8.166 | 28.792 | -11118.73 |

*All energy values are reported in kcal/mol.

**Table S4.** List of antibodies

| Target | Supplier | Identifier | Dilution |
|---|---|---|---|
| BTK | Cell Signaling Technology | 8547 | 1:1000 |
| GAPDH | ProteinTech | 60004-Ig-1 | 1:50000 |
| FLAG | Homemade (Clone M2) | | 0.5 µg/mL |

II. Biological Methods

**Cell Culture**

HEK293T cells were maintained in DMEM (Gibco 10566024) supplemented with 10% (v/v) heat-inactivated fetal bovine serum (VWR Avantor 76419-584). Ramos cells were maintained in RPMI 1640 (Gibco 11875093) supplemented with 10% (v/v) heat-inactivated fetal bovine serum (VWR Avantor 76419-584). Cells were passed for at least two generations after cryorecovery before they were used for assays. Cells were maintained in a humidified 37 ˚C incubator with 5% $CO_2$.

All cell lines tested mycoplasma negative using Mycoplasma Pro PCR Detection Kit (Applied Biological Materials Inc, G239).

When indicated, cells were treated with compound to a final DMSO concentration of 0.1%. HEK293T cells were detached with PBS (1 mL) and pelleted by centrifugation (1500 x g, 5 min) and pellets were washed once with ice-cold PBS (1 mL). Jurkat cells were pelleted by centrifugation (1500 x g, 5 min) and washed twice with ice-cold PBS (1 mL). Cells were lysed in lysis buffer (50 mM HEPES pH 7.5, 120 mM NaCl, 1% NP-40) on ice for 10 min and clarified by high-speed centrifugation (21000 x g, 10 min). Lysate concentrations were determined with protein BCA assay (Thermo Fisher, PIA53225) following manufacturer's protocols and adjusted to 1 mg/mL with additional lysis buffer.

**Recombinant protein expression and purification**

<u>BTK (wildtype), BTK(C481S) kinase domain</u>

Human BTK kinase domain (residues 389-659) containing mutations (M489A, R492A, E624A, K625A) was codon optimized, synthesized by Twist Biosciences and cloned into pFastBac vector using the Gibson Assembly method. The resulting construct contains N-terminal 6xHis tag and a Tobacco Etch Virus (TEV) protease cleavage site (ENLYFQG). Baculovirus was generated in Sf9 cells. 1 L of Sf9 cells were infected with P2 virus for 72 h. Cells were then pelleted by centrifugation (6500 x g, 15 min) and lysed in lysis buffer (20 mM HEPES pH 8, 200 mM NaCl, 5% glycerol, 1 mM TCEP, 5 mM imidazole). Lysate was clarified by high-speed centrifugation (19000 x g, 25 min). His-TEV-tagged protein was captured with Ni-NTA resin (Thermo Fisher #88222, 2 ml slurry per liter culture) at 4 °C for 1 h with constant end-to-end mixing. The loaded beads were then washed with lysis buffer and the protein was eluted with elution buffer (20 mM HEPES 8.0, 500 mM NaCl, 5% glycerol, 5 mM TCEP, 500 mM imidazole). To this protein solution, His-tagged TEV protease (0.025 mg TEV per milligram BTK protein) was added. The protein solution was diluted 1:10 v/v with Lysis Buffer and further purified with anion exchange chromatography (HiTrapQ column, Cytiva 17115301) using a NaCl gradient of 50 mM to 500 mM in 20 mM Tris pH 8, 5% glycerol buffer. Fractions containing pure BTK protein were pooled, concentrated, and flash frozen in liquid nitrogen.

**Intact protein mass spectrometry**

Purified BTK kinase domain (1 µM final) was incubated with compounds at 1 or 10 µM (1% v/v DMSO final) in 20 mM HEPES pH 7.5, 150 mM NaCl in a total volume of 100 µL. The extent of modification was assessed by electrospray MS using an Agilent Technologies 6545XT AdvanceBio LC/Q-TOF coupled to Agilent 1290 Infinity II LC HPLC eq

uipped with an ZORBAX RRHD Eclipse Plus C18 1.8 µm column (Agilent 959757-902). The mobile phase was a linear gradient of 10-98% acetonitrile/water + 0.05% formic acid. The spectra were processed in Agilent BioConfirm.

**Kinetic intact protein mass spectrometry**

Test compounds were prepared as 100× stock solutions in DMSO. Compounds were then diluted with SEC buffer (20 mM HEPES 7.5, 150 mM NaCl and 1 mM $MgCl_2$) to prepare a 2x stock. Recombinant proteins were diluted with SEC buffer to 2 µM. The compound solution was mixed with the protein solution at a 1:1 (v/v) ratio. The extent of modification was assessed by electrospray MS using an Agilent Technologies 6545XT AdvanceBio LC/Q-TOF coupled to Agilent 1290 Infinity II LC HPLC equipped with an ZORBAX RRHD Eclipse Plus C18 1.8 µm column (Agilent 959757-902). The mobile phase was a linear gradient of 10-98% acetonitrile/water + 0.05% formic acid. Injection time stamps were used to calculate elapsed time. The spectra were processed in Agilent BioConfirm.

**Crystallization**

Purified BTK(WT) and BTK(C481S) kinase domain was diluted to 50 µM in 20 mM HEPES pH 7.5, 150 mM NaCl and treated with 100 µM of compound **4B** to a final DMSO concentration of 1% overnight. 100% modification was confirmed by intact protein mass spectrometry. The BTK•**4B** complex was purified by size exclusion chromatography (Superdex200, 20 mM HEPES pH 8, 50 mM NaCl, 0.1 mM TCEP) and concentrated to 9 mg/mL.

For BTK(WT)•**4B**, 1 µL protein was mixed with 1 µL well buffer (0.1 M MIB, 27% PEG1500). The drop was seeded with crushed crystals. Crystals were grown at 20 ˚C in a 15-well plate using hanging-drop vapor diffusion method. Maximal crystal growth was achieved after 3 days. Crystals were transferred to a cryoprotectant solution (0.1 M MIB, 27% PEG1500, 25% glycerol), flash-frozen and stored in liquid nitrogen before X-ray diffraction.

For BTK(C481S)•**4B**, 1 µL protein was mixed with 1 µL well buffer (0.1 M MIB, 27% PEG1500). Crystals were grown at 20 ˚C in a 15-well plate using hanging-drop vapor diffusion method. Maximum crystal growth was observed after 14 days. Crystals were transferred to a cryoprotectant solution of 0.1 M MIB, 27% PEG1500, 25% glycerol, flash-frozen and stored in liquid nitrogen before X-ray diffraction.

0.1 M MIB: 0.025 M malonic acid, 0.0375 M imidazole, 0.0375 M boric acid

**X-Ray Data Collection and Structure Determination**

Datasets were collected at the Advanced Light Source Beamline 8.2.1 or 8.2.2. The dataset was indexed and integrated using iMOSFLM,[1] scaled with Aimless[2] and solved by molecular replacement using Phaser.[3] The crystal structure of ibrutinib-bound BTK kinase domain (PDB code: 5P9J) was used as the initial model. The structure was manually refined with Coot[4] and PHENIX.[5] Data collection and refinement statistics are listed in Table S1.

**Transfection of HEK293T cells**

HEK293T cells plated in 24-well plates (0.5e5 cells/well) and grown to 70% confluency. Cells were transfected with Lipofectamine 3000 (Invitrogen, L3000008) for 48 hours with 0.5 mg plasmid following manufacturer's protocols.

**Gel electrophoresis and immunoblot**

SDS-PAGE was run with Novex 4-12% Bis-Tris gel (Invitrogen) or SurePage 4-12% Bis-Tris gel (Genscript) in MOPS running buffer (Invitrogen) at 200 V following manufacturer's instructions. Protein bands were transferred onto 0.2-µM nitrocellulose membranes (Bio-Rad) using a wet-tank transfer apparatus (Bio-Rad) in 1x TOWBIN buffer with 10% methanol at 75V for 45 min. Membranes were blocked in 5% BSA-TBST for 1 h at 23 ˚C. Primary antibody binding was performed with the indicated antibodies diluted in 5% BSA-TBST at 4 ˚C for at least 16 h. After washing the membrane three times with TBST (5 min per wash), secondary antibodies (goat anti-rabbit IgG-IRDye 800 and goat anti-mouse IgG-IRDye 680, Li-COR) were added as solutions in 5% skim milk-TBST at dilutions recommended by the manufacturer. Secondary antibody binding was allowed to proceed for 1 h at 23 ˚C. The membrane was washed three times with TBST (5 min per wash) and imaged on a ChemiDoc MP Imaging System (BioRad).

**Chemical proteomic characterization of ibrutinib probes**

Ramos cells were seeded at 5e6 cells/mL in a Protein LoBind 96-well 2 mL deep well plate (Eppendorf) on the day of experiment. Cells were treated in triplicate with DMSO, 20µM Ibrutinib or compounds **4b** and **5a** and incubated for 1 h at 37°C, 5% $CO_2$. Subsequently, cells were treated with 1µM probes 6-8 and incubated for an additional 1 h at 37°C, 5% $CO_2$. Cells were harvested by spinning at 350xg for 3 min and washed with DPBS (Gibco) and lysed in the plate with 350 µL 50 mM HEPES pH 7.5, 150 mM NaCl, 1.5 mM $MgCl_2$, 0.8% NP-40, 1% SDS, 1x Halt Protease inhibitor (Thermo Scientific). Samples were heated at 95°C for 15 min and sonicated to sheer DNA. Lysate were prepped for copper-catalyzed azide-alkyne cycloaddition (CuAAC) by addition of 44 µL Click Mix (4 µL Biotin Picolyl Azide [Vector], 24 µL 1.7 mM TBTA in 4:1 tBuOH:DMSO, 8 µL 50 mM $CuSO_4$ in $H_2O$, 8 µL 50 mM TCEP in $H_2O$) per well and incubated for 1 h at RT with constant mixing. Samples were precipitated with addition of 1.6 mL cold acetone and stored at -80°C overnight. Precipitated proteins were collected by centrifugation (3000xg) for 30 min at RT, acetone was aspirated, and protein pellets were resuspended in 200 µL 50 mM HEPES pH 7.5, 1% SDS. Once reconstituted, the samples were diluted up to 900 µL with 50 mM HEPES pH 7.5 containing 0.2% SDS. 100 µL of Sera-Mag SpeedBeads Neutravidin-Coated Magnetic Particles (Cytiva) were added and incubated overnight at RT on an end-over-end mixer. Samples were washed on a KingFisher (Thermo) with 2x 50 mM HEPES pH 7.5, 0.2% SDS; 1x 10 mM DTT in 50 mM HEPES pH 8.0; 1x 50 mM iodoacetamide in 50 mM HEPES pH 8.0; 2x 50 mM HEPES pH 8.0; and beads were deposited in 200 µL 50 mM Ammonium Bicarbonate pH 8.0. 5 µg Trypsin/LysC (Promega) was added to each well containing beads and digested for 16 h at 37°C at 1000 rpm. Digested peptide supe was collected and concentrated by evaporation and reconstituted in 100 µL 2% acetonitrile, 0.1% formic acid for LC-MS/MS analysis. Samples were acquired on Astral Zoom Mass Spectrometer (Thermo) as described below. Raw files were processed with FragPipe 24.0 using the DIA_SpecLib_Quant workflow, using default parameters, searched against a 2024 reference human proteome FASTA file.[6,7]

**Cysteine Profiling Proteomics**

Ramos cells were seeded at 1e6 cells per well in a 12-well plate and cultured until confluent. Cells were then treated with DMSO, Ibrutinib, compound **4b**, or compound **5a** at 25 or 5 µM for 1 h in triplicate. Cells were washed and lysed in the well in 50 mM HEPES pH 7.5, 150 mM NaCl, 1.5 mM MgCl2, 0.8% NP-40, 0.1% nuclease (Thermo). Lysates were cleared by centrifugation at 3200 rcf for 10 minutes at 4 °C. Protein content of the resulting lysates were normalized, and 500 µg/sample was treated with desthiobiotin iodoacetamide (DBIA) at 500 µM for 1 h at room temperature. Lysates were reduced with 5 mM DTT for 20 minutes at room temperature and alkylated with 15mM iodoacetamide for 20 minutes at room temperature. Excess reagents were removed by cleanup through 3mg/sample Sera-Mag Speedbead carboxylate-modified [E3 and E7] magnetic particles (Cytiva), followed by on-bead digestion with 10 µg LysC/trypsin (Promega) overnight at 37 °C. Peptides were diluted two fold with 50 mM HEPES pH 8, 1% SDS, and probe modified peptides were enriched by incubating with 3mg/sample Sera-Mag SpeedBeads Neutravidin-Coated Magnetic Particles (Cytiva) for 90 min at room temperature with shaking. Beads were washed 2x with 800 µL 0.2% SDS, 2x with 800 µL PBS, and 1x 800 µL water. Peptides were eluted from the beads by incubation in 200 µL 50% acetonitrile, 0.1% trifluoroacetic acid buffer for 20 minutes at room temperature with agitation. Beads were removed and the eluted peptides were concentrated by evaporation. The resulting concentrated peptides were resuspended in 20 µL 2.5% formic acid for analysis by LC-MS/MS.

Samples were analyzed by nanoLC-MS/MS using a Vanquish Neo UHPLC system (Thermo) interfaced with an Orbitrap Astral Zoom Mass Spectrometer (Thermo). Samples (10 µL) were injected onto a C18 5 µm trapping column (300 µm x 5 mm) (Thermo) using 0.1% formic acid/2% acetonitrile in water at a flow rate of 3 μL/min. Trapped peptides were then introduced into the separation column and eluted at 3 μL/minute using a mobile phase A: 2% acetonitrile + 0.1% formic acid in water and a mobile phase B: 70% acetonitrile + 0.1% formic acid in water over a gradient of 5-55% B over 10 min. MS1 scans were acquired from m/z 380-980 at 240,000 mass resolution. Astral DIA scans were acquired from 150-2000 m/z, HCD with 30% collision and 5 Da isolation window. MS2 AGC target was 5.00e4 using a 3 ms maximum injection time. Raw files were processed using FragPipe 24.0 utilizing the chemprot-ABPP-IADTB workflow searched against a 2024 reference human proteome FASTA file.[6,7] Cleavage enzyme set to trypsin/chymotrypsin with 2 allowed missed cleavages. DBIA mass delta at cysteine set to 296.1848, max occurrences at 3. Carbidomethylation at cysteine set to 57.02146, max occurrences at 3. Standard DIA settings were used.

**Liquid chromatography tandem mass spectrometry analysis of proteins covalently modified by general kinase probes**

One day before treatment, HEK293T cells were added to 10-cm tissue culture dishes (4e6 cells/dish). The next day, cells were treated with 2 μM probe **9**, probe **10**, or **XO44** for 1 h. Each treatment condition was performed in biological triplicate. Cells were then placed on ice, and the media removed by aspiration. Cells were scraped in ice-cold PBS (10 mL) and pelleted by centrifugation (500*g*, 5 min). Cell pellets were washed with ice-cold PBS (1 x 1 mL). Cells were resuspended in PBS (500 μL) supplemented with protease and phosphatase inhibitors (cOmplete and phosSTOP; Roche) and lysed by probe-tip sonication. Concentrations of lysates were determined using a BCA protein assay (Thermo Fisher Scientific) and adjusted to 2 mg $ml^{-1}$ with additional PBS. Cell lysate (500 μL) were mixed with 5× Biotin-$N_3$ Click Master Mix and the reaction was allowed to proceed at 23 °C for 1 h. Proteins were then precipitated by addition of ice-cold methanol (600 μL) and chloroform (200 μL). Samples were vortexed and centrifuged, and both layers were carefully aspirated to leave a protein disk. The protein disk was resuspended in methanol, sonicated briefly in a water bath sonicator, and centrifuged. The supernatant was aspirated and the protein pellet was allowed to air dry for 10 min. Each pellet was redissolved in 8 M urea in PBS + 2% SDS (500 μL). To each sample, 25 μL of 200 mM DTT was added and samples were incubated at 65 ˚C for 15 min. Samples were cooled to 37 ˚C and 50 μL of 200 mM iodoacetamide was added. Samples were incubated at 37 ˚C in the dark. After 30 min, 50 μL of 20% SDS was added and samples were resuspended in 5 mL PBS and incubated with 100 μL of streptavidin agarose beads (Pierce, 20353, 50% v/v suspension, washed with PBS) for 1.5 h with constant end-to-end rotation. The beads were washed with PBS + 0.2% SDS (2 x 10 mL) and PBS (1 x 10 mL) and then transferred to low binding microcentrifuge tubes (Pierce) with PBS (1 mL). Beads were then washed with water (2 x 1 mL) and 20 mM HEPES pH 8 (2 x 1 mL). Each wash was performed by spinning down the resin (1500*g*, 2 min) and removing the supernatant. After the final wash, the beads were resuspended in 200 μL digestion buffer (20 mM HEPES, 1 mM $CaCl_2$, pH 8) with 500 ng trypsin, and the samples were incubated at 37 ˚C overnight with constant agitation (1000 r.p.m., orbital shaker). The beads were collected with spin filter columns (Pierce) and the tryptic peptides were collected into new low binding microcentrifuge tubes. 35 μL of acetonitrile was added to each sample. Samples were briefly vortexed and the tryptic peptides were labeled with 0.1 mg of TMT10plex™ Isobaric Label Reagents and Kit (Thermo Fisher Scientific, 90110) following manufacturer's instructions. After completion of labeling reaction and quenching with 0.5% hydroxylamine, each sample was acidified by 20 μL of formic acid. The acidified peptide mixtures were pooled in a single tube and dried by vacuum centrifugation (Genevac). The residue was resuspended in 5% acetonitrile + 0.1% formic acid and desalted with Omix C18 pipette tips (Agilent), eluting in 80% acetonitrile + 0.1% formic acid. Desalted peptides were dried by vacuum centrifugation (Genevac). TThe dried peptides were fractionated using a Pierce High pH Reverse-Phase Peptide Fractionation Kit (Thermo Fisher Scientific, 84868) following the manufacturer's instructions into 8 fractions which were concatenated into 4 fractions that were dried individually and resuspended in 20 μL of 0.5 formic acid. Peptides were Easy-Spray nano-HPLC column (Thermo Fisher Scientific, ES900; 150 mm length, 3 μM particle size, 100-Å particle size) over a 180-min gradient of 2%–37% acetonitrile/water + 0.1% formic acid and analyzed by a Orbitrap Exploris 480 mass spectrometer (precursor range: 350–1,200 *m*/*z*, charge state 2–5, MS1 Orbitrap resolution 120,000, max injection time auto, RF lens 50%; dynamic exclusion: 45 s; MS2 ion trap precursor isolation window 0.7 *m*/*z*; CID collision energy: 36%; constant modification of carbamidomethyl (Cys), TMT6plex (Lys and N-terminal) and variable modification of oxidation (Met). Peptides were searched against the SwissProt *Homo sapiens* reference proteome (accessed 2025.10.21), using FragPipe (21.1) with MSFragger (4.0).[6,7] Fold-enrichment was calculated as the ratio of the geometric means of peptide intensities of three biological replicates between the three conditions. *P* values were calculated using unpaired, two-tailed Student's *t*-test.

**Covalent Docking**

The X-ray crystal structure of the target protein was retrieved from the Protein Data Bank (AKT1: PDB 4GV1[8]; BTK: PDB 5P9J[9]). The structure was refined and prepared using the Protein Preparation Wizard with Maestro (14.4.133, Schrödinger Release 2025-2) and missing side chains and loops were modeled with Prime.[10] The protein was optimized at pH 7.0 ± 2.0 with PropKa and the protein was minimized with OPLS4 to a RMSD of all atoms to 0.30 Å. Ligands were prepared using the LigPrep module and minimized with OPLS4. Covalent docking was conducted through the CovDock workflow through Glide.[11] The receptor grid was centered on the nucleophilic residue and sized to the existing ligands in the crystal structures. Docking was performed using Pose Prediction and candidates within 2.5 kcal/mol from the energy minimum were selected for further refinement through Prime.

III. Chemistry Methods

**General Methods**

All reactions sensitive to air or moisture were conducted under nitrogen atmosphere in dry solvents under anhydrous conditions, unless otherwise noted. Air- and moisture-sensitive liquids were transferred via syringe. Solutions were concentrated by rotary evaporation at or below 40 °C. Analytical thin-layer chromatography (TLC) was performed using glass plates pre-coated with silica gel (0.25-mm, 60-Å pore size, Millipore Sigma) impregnated with a fluorescent indicator (254 nm). TLC plates were visualized by exposure to ultraviolet light (UV), then were stained by an 2% aqueous solution of potassium permanganate followed by brief heating by heat gun. Column chromatography was performed on silica gel (RediSep® Silver Silica Gel Disposable Flash Columns 4 grams from Teledyne ISCO, 692203304) or preparative high-performance liquid chromatography (prepHPLC, CombiFlash EZprep, C18 20x150mm column).

All solvents and reagents were used as obtained from commercial sources like Fisher Scientific, Sigma-Aldrich, Aaron Chemical, Ambeed, Sigma Aldrich, AK Scientific and Chemscene and used without further purification unless otherwise stated.

$^{1}$H NMR and $^{13}$C NMR spectra for characterization of new compounds and monitoring reactions were collected in $CDCl_e$ (Cambridge Isotope Laboratories, Cambridge, MA) on Bruker AVQ-400, AVB-400, AV-600, NEO-500, and NEO-501 MHz spectrometers. Proton chemical shifts are expressed in parts per million (ppm, δ scale) and are referenced to residual protium in the NMR solvent ($CHCl_3$: δ 7.26). Splitting patterns were indicated as follows: br, broad; s, singlet; d, doublet; t, triplet; q, quartet; m, multiplet; dd, doublet of doublets; dt, doublet of triplets. High-resolution mass spectrometry was performed on an Agilent Technologies 6545XT AdvanceBio LC/Q-TOF (ESI source) coupled to Agilent 1290 Infinity II LC HPLC.

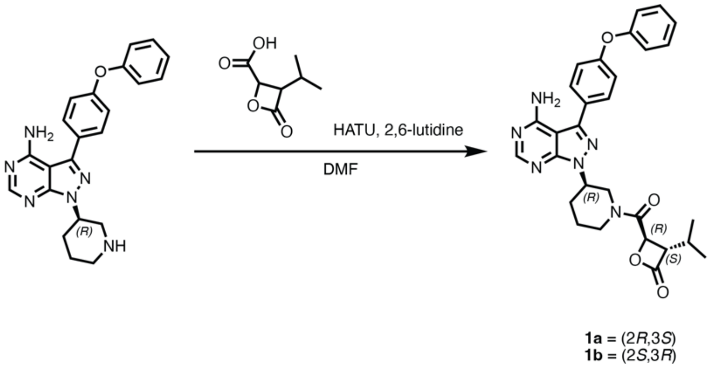

tert-butyl (2*R*,3*S*)-3-isopropyl-4-oxo-oxetane-2-carboxylate and tert-butyl (2*S*,3*R*)-3-isopropyl-4-oxo-oxetane-2-carboxylate

Prepared according to literature protocols.[12]

(3*S*,4*R*)-4-((*R*)-3-(4-amino-3-(3-phenoxyphenyl)-1*H*-pyrazolo[3,4-*d*]pyrimidin-1-yl)piperidine-1-carbonyl)-3-isopropyloxetan-2-one (**1a**)

To an oven dried vial, (2*R*,3*S*)-3-isopropyl-4-oxooxetane-2-carboxylic acid (25 mg, 0.16 mmol, 3.0 eq) and HATU (39 mg, 0.10 mmol, 2.0 eq) were dissolved in DMF (1 mL) and cooled to 0 ˚C. To this, 2,6-lutidine (25 µL, 0.22 mmol, 4.2 eq) was added and the reaction was stirred at 0 ˚C for 30 min. (*R*)-3-(3-phenoxyphenyl)-1-(piperidin-3-yl)-1*H*-pyrazolo[3,4-*d*]pyrimidin-4-amine (20 mg, 0.052 mmol, 1 eq) was added, the reaction was allowed to proceed at room temperature for 30 min. The reaction was diluted in DCM (1 mL) and washed with saturated sodium bicarbonate (2 mL). The aqueous layer was then extracted DCM (3 x 2 mL). The combined organic layers were washed with brine (2 mL), dried over sodium sulfate, and concentrated. The residue was purified by reverse-phase C18 HPLC. Elution was performed with a linear gradient of 10–95% acetonitrile–water + 0.1% formic acid, and the product-containing fractions were lyophilized to afford **1a** as a white solid (12 mg, 0.023 mmol, 44%).

**HRMS (ESI):** calculated for $C_{29}H_{30}N_6O_4^+$ $[M+H]^+$ 527.2401, found 527.2408.

**$^1$H NMR (500 MHz, $CDCl_3$)** (mixture of rotamers, 1:0.9 major:minor) δ / ppm = 8.27 (s, major rotamer, 1H), 8.25 (s, minor rotamer, 0.9H), 7.60 (dd, major + minor rotamer, *J* = 8.7, 6.4 Hz, 3.8H), 7.44 – 7.36 (m, major + minor rotamer, 3.8H), 7.23 – 7.04 (m, major + minor rotamer, 9.5H), 4.99 – 4.85 (m, major + minor rotamer, 1.9H), 4.81 (d, minor rotamer, *J* = 4.2 Hz, 0.9H), 4.78 (d, major rotamer, *J* = 4.2 Hz, 1H), 4.65 (dd, minor rotamer, *J* = 12.9, 4.3 Hz, 0.9H), 4.18 (dt, major rotamer *J* = 13.3, 4.6 Hz, 1H), 4.06 (dd, major rotamer *J* = 13.4, 4.5 Hz, 1H), 4.00 (ddt, major + minor rotamer, *J* = 12.8, 8.2, 4.4 Hz, 2.9H), 3.91 (dt, *J* = 14.1 Hz, 1H), 3.52 (dd, minor rotamer, *J* = 12.9, 10.3 Hz, 0.9H), 3.32 (ddd, minor rotamer, *J* = 14.3, 11.7, 3.0 Hz, 0.9H), 3.26 (ddd, major rotamer, *J* = 13.4, 10.1, 3.3 Hz, 1H), 2.36 (m, major + minor rotamer, 1.9H), 2.29 – 2.21 (m, major + minor rotamer, 1.9H), 2.21 – 2.01 (m, major + minor rotamer, 2.9H), 1.94 (m, major rotamer, 1H), 1.74 (m, major + minor rotamer, 1.9H), 1.13 (d, *J* = 6.7 Hz, 2.7H), 1.08 (d, *J* = 6.9 Hz, 2.7H), 1.03 (d, *J* = 6.8 Hz, 3H), 0.94 (d, *J* = 6.7 Hz, 3H).

(3*R*,4*S*)-4-((*R*)-3-(4-amino-3-(3-phenoxyphenyl)-1*H*-pyrazolo[3,4-*d*]pyrimidin-1-yl)piperidine-1-carbonyl)-3-isopropyloxetan-2-one (**1b**)

To an oven dried vial, (2*S*,3*R*)-3-isopropyl-4-oxooxetane-2-carboxylic acid (24.4 mg, 0.155 mmol, 3.00 Eq) and HATU (39.2 mg, 0.103 mmol, 2.00 Eq) were dissolved in DMF (1 mL) and cooled to 0 ˚C. To this, 2,6-lutidine (25 µL, 0.22 mmol, 4.0 eq) was added and the reaction was stirred at 0 ˚C for 30 min.

($R$)-3-(3-phenoxyphenyl)-1-(piperidin-3-yl)-1$H$-pyrazolo[3,4-$d$]pyrimidin-4-amine (19.9 mg, 0.519 mmol, 1.00 Eq) was added, the reaction was allowed to proceed at room temperature for 30 min. The reaction was diluted in DCM (1 mL) and washed with saturated sodium bicarbonate (2 mL). The organic layer was extracted DCM (3 x 2 mL). The combined organic layers were washed with brine (2 mL), dried over sodium sulfate, and concentrated. The residue was purified by reverse-phase C18 HPLC. Elution was performed with a linear gradient of 10–95% acetonitrile–water + 0.1% formic acid, and the product-containing fractions were lyophilized to afford **1b** as a white solid (16 mg, 0.051 mmol, 59%).

**HRMS (ESI):** calculated for $C_{29}H_{30}N_6O_4^+$ $[M+H]^+$ 527.2401, found 527.2401.

**$^1$H NMR (500 MHz, $CDCl_3$)** (mixture of rotamers 1:1) δ / ppm = 8.34 (s, 0.5H), 8.33 (s, 0.5H), 7.63 (d, $J$ = 7.6 Hz, 2H), 7.43 – 7.36 (m, 2H), 7.21 – 7.12 (m, 3H), 7.12 – 7.05 (m, 2H), 4.94 – 4.87 (m, 1H), 4.86 (d, $J$ = 4.3 Hz, 0.5H), 4.80 (d, $J$ = 4.2 Hz, 0.5H), 4.71 (dd, $J$ = 12.6, 4.3 Hz, 0.5H), 4.52 (d, $J$ = 13.2 Hz, 0.5H), 4.10 (d, $J$ = 3.8 Hz, 0.5H), 4.07 (dd, $J$ = 8.2, 4.1 Hz, 0.5H), 3.99 (dd, $J$ = 8.0, 4.2 Hz, 0.5H), 3.95 (d, $J$ = 13.5 Hz, 0.5H), 3.76 (dd, $J$ = 13.7, 10.6 Hz, 0.5H), 3.45 (dd, $J$ = 12.8, 10.7 Hz, 0.5H), 3.22 (td, $J$ = 12.7, 2.7 Hz, 0.5H), 2.97 – 2.87 (m, 1H), 2.45 (qd, $J$ = 12.4, 4.1 Hz, 0.5H), 2.40 – 2.31 (m, 0.5H), 2.30 – 2.23 (m, 1H), 2.15 (dp, $J$ = 7.9, 6.6 Hz, 1H), 2.08 – 1.97 (m, 1H), 1.80 (dt, $J$ = 41.0, 12.8 Hz, 1H), 1.13 – 1.04 (m, 6H). 1.12 (d, $J$ = 5.7 Hz, 1.5H), 1.11 (d, $J$ = 5.7 Hz, 1.5H), 1.08 (d, $J$ = 6.8 Hz, 1.5H), 1.05 (d, $J$ = 6.8 Hz, 1.5H).

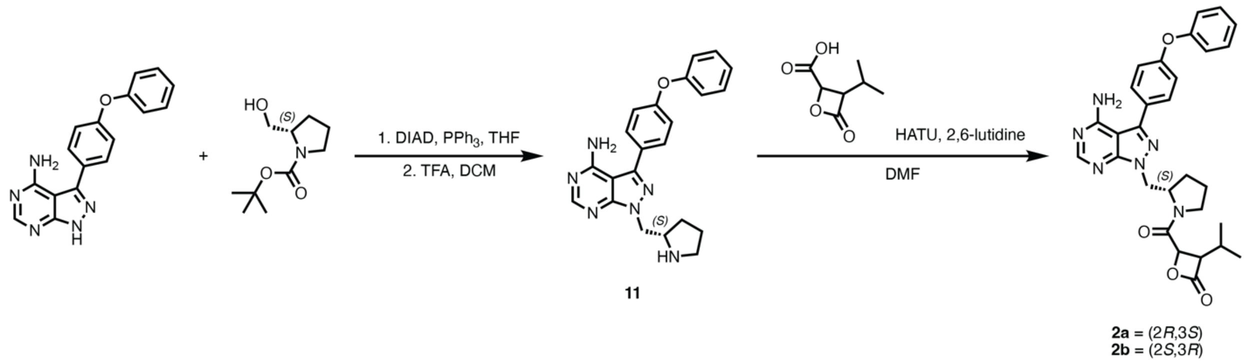

<u>($S$)-2-((4-amino-3-(4-phenoxyphenyl)-1H-pyrazolo[3,4-$d$]pyrimidin-1-yl)methyl)pyrrolidin-1-ium 2,2,2-trifluoroacetate (**11**)</u>

In an oven-dried 20-mL scintillation vial, 3-(4-phenoxyphenyl)-1H-pyrazolo[3,4-$d$]pyrimidin-4-amine (102.9 mg, 1.000 Eq, 339.2 µmol), tert-butyl ($S$)-2-(hydroxymethyl)pyrrolidine-1-carboxylate (103.7 mg, 1.519 Eq, 0.5152 mmol), and triphenylphosphine (137.6 mg, 1.546 Eq, 524.6 µmol) were dissolved in THF (3.4 mL). To this solution, diisopropyl azodicarboxylate (104 mg, 0.100 mL, 1.52 Eq, 514 µmol) was added slowly and the reaction was stirred at room temperature for 16 h. The reaction was concentrated and purified by flash column chromatography (1% methanol in DCM) to afford the product as a clear oil. This oil was redissolved in DCM (2 mL) and TFA (2 mL) and allowed to stand overnight. The solution was then concentrated, and the product was triturated out with diethyl ether to afford the **11** as a white crystalline substance (65.5 mg, 0.339 mmol, 39%).

**HRMS (ESI):** calculated for $C_{22}H_{23}N_6O^+$ $[M+H]^+$ 387.1928, found 387.1938.

**$^1$H NMR (500 MHz, $CDCl_3$):** δ / ppm = 8.25 (s, 1H), 7.57 – 7.51 (m, 2H), 7.45 – 7.37 (m, 2H), 7.25 – 7.18 (m, 1H), 7.16 – 7.11 (m, 2H), 7.11 – 7.05 (m, 2H), 6.35 (s, 1H), 4.85 (dd, $J$ = 15.0, 8.0 Hz, 1H), 4.76 (dd, $J$ = 14.9, 3.9 Hz, 1H), 4.29 (s, 1H), 3.34 (t, $J$ = 7.3 Hz, 2H), 2.33 (td, $J$ = 12.5, 7.0 Hz, 1H), 2.10 (m, 2H), 1.87 (dq, $J$ = 13.1, 8.7 Hz, 1H).

(3*S*,4*R*)-4-((*S*)-2-((4-amino-3-(4-phenoxyphenyl)-1H-pyrazolo[3,4-*d*]pyrimidin-1-yl)methyl)pyrrolidine-1-carbonyl)-3-isopropyloxetan-2-one (**2a**)

In an oven dried 1-dram vial, the (2*R*,3*S*)-3-isopropyl-4-oxooxetane-2-carboxylic acid (1.4 mg, 1 Eq, 8.9 µmol) and HATU (6.5 mg, 1.9 Eq, 17 µmol) were dissolved in DCM (100 µL) and cooled to 0 ˚C. To this, 2,6-lutidine (5.4 mg, 5.8 µL, 5.7 Eq, 0.050 mmol) was added and the solution was allowed to stir on ice for 30 minutes. (*S*)-2-((4-amino-3-(4-phenoxyphenyl)-1H-pyrazolo[3,4-*d*]pyrimidin-1-yl)methyl)pyrrolidin-1-ium 2,2,2-trifluoroacetate (7.2 mg, 1.6 Eq, 14 µmol) was then added and the solution was stirred at room temperature for 30 minutes. The reaction was diluted in DCM (1 mL) and washed with saturated sodium bicarbonate (2 mL). The aqueous layer was extracted with DCM (3 x 2 mL). The combined organic layers were washed with brine (2 mL), dried over sodium sulfate, and concentrated. The residue was purified by reverse-phase C18 HPLC. Elution was performed with a linear gradient of 10–95% acetonitrile–water + 0.1% formic acid, and the product-containing fractions were lyophilized to afford **2a** as a white solid (0.8 mg, 0.009 mmol, 20%).

**HRMS (ESI):** calculated for $C_{29}H_{31}N_6O_4^+$ $[M+H]^+$ 527.2401, found 527.2412.

**$^1$H NMR (500 MHz, $CDCl_3$):** (mixture of rotamers, 1:0.4 major:minor) δ / ppm = 8.33 (s, major + minor rotamer, 1.4 H), 7.64 (d, major + minor rotamer, *J* = 8.5, 2.8H), 7.39 (t, major + minor rotamer, *J* = 7.8 Hz, 2.8H), 7.20 – 7.10 (m, major + minor rotamer, 4.2H), 7.12 – 7.06 (m, major + minor rotamer 2.8H), 5.86 (br s, major + minor rotamer, 2.8H), 4.90 (d, minor rotamer, *J* = 4.1 Hz, 0.4H), 4.83 (dd, major rotamer, *J* = 13.9, 5.4 Hz, 1H), 4.69 – 4.64 (m, major + minor 1.4H), 4.65 (d, major rotamer, *J* = 4.2 Hz, 1H), 4.61 (dd, major rotamer, *J* = 13.9, 4.0 Hz, 1H), 4.55 – 4.42 (m, minor rotamer, 0.8H), 3.99 (dd, minor rotamer, *J* = 7.7, 4.0 Hz, 0.4H), 3.93 (dd, major rotamer, *J* = 8.2, 4.2 Hz, 1H), 3.66 – 3.57 (m, minor rotamer, 0.4H), 3.53 – 3.38 (m, major + minor rotamer, 2.4H), 2.19 – 1.92 (m, major + minor, 4.2H), 1.85 – 1.80 (m, major + minor rotamer, 1.4H), 1.70 – 1.63 (m, major + minor rotamer, 1.4H), 1.11 (d, major rotamer, *J* = 6.7 Hz, 3H), 1.07 (d, minor rotamer, *J* = 6.6 Hz, 1.2H), 1.07 (d, major rotamer, *J* = 6.8 Hz, 3H), 1.00 (d, minor rotamer, *J* = 6.7 Hz, 1.2H).

(3*R*,4*S*)-4-((*S*)-2-((4-amino-3-(4-phenoxyphenyl)-1H-pyrazolo[3,4-*d*]pyrimidin-1-yl)methyl)pyrrolidine-1-carbonyl)-3-isopropyloxetan-2-one (**2b**)

In an oven dried 1-dram vial, (2*S*,3*R*)-3-isopropyl-4-oxooxetane-2-carboxylic acid (1.4 mg, 1 Eq, 8.9 µmol) and HATU (6.5 mg, 1.9 Eq, 0.017 mmol) were dissolved in DMF (100 µL) and cooled to 0 ˚C. To this, 2,6-lutidine (5.4 mg, 5.8 µL, 5.7 Eq, 0.050 mmol) was added and the solution was allowed to stir on ice for 30 minutes. (*S*)-2-((4-amino-3-(4-phenoxyphenyl)-1H-pyrazolo[3,4-*d*]pyrimidin-1-yl)methyl)pyrrolidin-1-ium 2,2,2-trifluoroacetate (6.3 mg, 1.4 Eq, 0.013 mmol) was then added and the solution was stirred at room temperature for 30 minutes. The reaction was diluted in DCM (1 mL) and washed with saturated sodium bicarbonate (2 mL). The organic layer was extracted DCM (3 x 2 mL). The combined organic layers were washed with brine (2 mL), dried over sodium sulfate, and concentrated. The residue was purified by reverse-phase C18 HPLC. Elution was performed with a linear gradient of 10–95% acetonitrile–water + 0.1% formic acid, and the product-containing fractions were lyophilized to afford **2b** as a white solid (1.0 mg, 0.0089 mmol, 21%).

**HRMS (ESI):** calculated for $C_{29}H_{31}N_6O_4^+$ $[M+H]^+$ 527.2401, found 527.2410.

**$^1$H NMR (500 MHz, $CDCl_3$)**: (mixture of rotamers, 1:0.4 major:minor) δ / ppm = 8.36 (s, minor rotamer, 0.4H), 8.35 (s, major rotamer, 1H), 7.68 – 7.59 (m, major + minor rotamer, 2.8H), 7.43 – 7.36 (m, major + minor rotamer, 2.8H), 7.22 – 7.12 (m, major + minor rotamer, 4.2H), 7.11 – 7.06 (m, major +

minor rotamer, 2.8H), 5.55 (br s, major + minor rotamer, 2.8H), 5.09 (d, minor rotamer, *J* = 4.0 Hz, 0.4H), 4.87 (dd, major rotamer, *J* = 14.1, 4.8 Hz, 1H), 4.71 – 4.65 (m, major + minor rotamer, 1.4 H), 4.66 (d, major rotamer, *J* = 4.1 Hz, 1H), 4.63 – 4.53 (m, major + minor rotamer, 1.4H), 4.45 (dd, minor rotamer, *J* = 13.8, 8.5 Hz, 0.4H), 3.94 (dd, major rotamer, *J* = 8.0, 4.0 Hz, 1H), 3.86 (dd, minor rotamer, *J* = 7.6, 4.0 Hz, 0.4H), 3.68 – 3.62 (m, minor rotamer, 0.4H), 3.59 – 3.54 (m, major _ minor rotamer, 1.4H), 3.31 – 3.26 (m, major rotamer, 1H), 2.13 – 1.98 (m, major + minor rotamer, 4.2H), 1.90 – 1.79 (m, major + minor rotamer, 1.4H), 1.50 – 1.41 (m, major + minor rotamer, 1.4H), 1.10 (d, minor rotamer, *J* = 6.8 Hz, 1.2H), 1.08 (d, major rotamer, *J* = 6.7 Hz, 3H), 1.07 (d, minor rotamer, *J* = 7.4, 1.2H), 1.01 (d, major rotamer, *J* = 6.7 Hz, 3H).

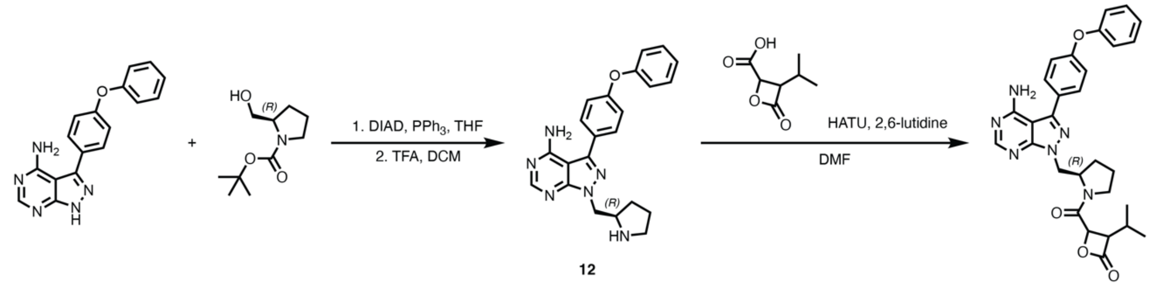

12

**3a** = (2*R*,3*S*)
**3b** = (2*S*,3*R*)

<u>(*R*)-2-((4-amino-3-(4-phenoxyphenyl)-1H-pyrazolo[3,4-*d*]pyrimidin-1-yl)methyl)pyrrolidin-1-ium 2,2,2-trifluoroacetate (**12**)</u>

In an oven-dried 20-mL scintillation vial, 3-(4-phenoxyphenyl)-1H-pyrazolo[3,4-*d*]pyrimidin-4-amine (103.5 mg, 1.000 Eq, 0.3412 mmol), triphenylphosphine (129.7 mg, 1.449 Eq, 0.4945 mmol), and tert-butyl (*R*)-2-(hydroxymethyl)pyrrolidine-1-carboxylate (104.2 mg, 1.517 Eq, 0.5177 mmol) were dissolved in anhydrous THF (3.4 mL). To the reaction, diisopropyl azodicarboxylate (104 mg, 100 µL, 1.51 Eq, 0.514 mmol) was added slowly and the reaction was stirred at room temperature for 16 h. The reaction was concentrated and purified by flash column chromatography (1% methanol in DCM) and fractions containing the product were dried down to afford a clear oil. This oil was redissolved in DCM (2 mL) and TFA (2 mL) and allowed to stand overnight. The solution was then concentrated, and the product was triturated out with diethyl ether to afford **12** as a white solid (36.3 mg, 0.0725 mmol, 21 %).

**$^1$H NMR (500 MHz, CDCl$_3$):** δ / ppm = 8.26 (s, 1H), 7.58 – 7.51 (m, 2H), 7.46 – 7.38 (m, 2H), 7.26 – 7.19 (m, 1H), 7.17 – 7.11 (m, 2H), 7.11 – 7.06 (m, 2H), 6.35 (s, 1H), 4.88 – 4.75 (m, 2H), 4.30 (m, 1H), 3.37 (m, 2H), 2.35 (dq, *J* = 13.2, 6.4 Hz, 1H), 2.11 (m, 2H), 1.86 (dq, *J* = 13.2, 8.8 Hz, 1H).

<u>(3*S*,4*R*)-4-((*R*)-2-((4-amino-3-(4-phenoxyphenyl)-1H-pyrazolo[3,4-*d*]pyrimidin-1-yl)methyl)pyrrolidine-1-carbonyl)-3-isopropyloxetan-2-one (**3a**)</u>

In an oven dried 1-dram vial, (2*R*,3*S*)-3-isopropyl-4-oxooxetane-2-carboxylic acid (5.69 mg, 1.20 Eq, 0.0360 mmol) and HATU (13.7 mg, 1.20 Eq, 0.0360 µmol) were dissolved in DMF (0.3 mL) and the mixture was cooled to 0 ˚C. To this, 2,6-lutidine (16.1 mg, 17.4 µL, 5 Eq, 0.150 mmol) was added and the solution was allowed to stir on ice for 30 minutes. (*R*)-2-((4-amino-3-(4-phenoxyphenyl)-1H-pyrazolo[3,4-*d*]pyrimidin-1-yl)methyl)pyrrolidin-1-ium 2,2,2-trifluoroacetate (15.0 mg, 1.00 Eq, 0.0300 mmol) was then added to the reaction and mixture was stirred at room temperature for 30 minutes. The reaction was diluted in DCM (1 mL) and washed with saturated sodium bicarbonate (2 mL). The organic layer was extracted DCM (3 x 2 mL). The combined organic layers were washed with brine (2 mL), dried over sodium sulfate, and concentrated. The residue was purified by reverse-phase C18 HPLC. Elution was performed with a linear gradient of 10–95% acetonitrile–water + 0.1% formic acid, and the product-containing fractions were lyophilized to afford the **3a** as a white solid (2.5 mg, 0.030 mmol, 16%).

**HRMS (ESI):** calculated for $C_{29}H_{31}N_6O_4^+$ [M+H]$^+$ 527.2401, found 527.2422.

**$^1$H NMR (500 MHz, CDCl$_3$)**: (mixture of rotamers, 1:0.25 major:minor) δ / ppm = 8.36 (s, minor rotamer, 0.25H), 8.35 (s, major rotamer, 1H), 7.68 – 7.59 (m, major + minor rotamer, 2.5H), 7.43 – 7.36 (m, major + minor rotamer, 2.5H), 7.22 – 7.12 (m, major + minor rotamer, 3.75H), 7.11 – 7.06 (m, major + minor rotamer, 2.5H), 5.60 (br s, major + minor rotamer, 2H), 5.09 (d, minor rotamer, *J* = 4.1 Hz, 0.25H), 4.87 (dd, major rotamer, *J* = 14.1, 4.8 Hz, 1H), 4.71 – 4.65 (m, major + minor rotamer, 1.25H), 4.66 (d, major rotamer, *J* = 4.1 Hz, 1H), 4.63 – 4.53 (m, major + minor rotamer, 1.25H), 4.45 (dd, minor rotamer, *J* = 13.8, 8.5 Hz, 0.25H), 3.94 (dd, major rotamer, *J* = 7.9, 4.1 Hz, 1H), 3.86 (dd, minor rotamer, *J* = 7.6, 4.0 Hz, 0.25H), 3.70 – 3.61 (m, minor rotamer, 0.5H), 3.61 – 3.52 (m, major rotamer, 1H), 3.28 (ddd, major rotamer, *J* = 9.8, 8.5, 4.2 Hz, 1H), 2.21 – 1.95 (m, major + minor rotamer, 3.75H), 1.90 – 1.79 (m, major + minor rotamer, 1.25H), 1.50 – 1.41 (m, major + minor rotamer, 1.25H), 1.10 (d, minor rotamer, *J* = 6.8 Hz, 0.75H), 1.08 (d, *J* = 6.7 Hz, 3H), 1.07 (d, *J* = 7.4 Hz, 1H), 1.01 (d, *J* = 6.7 Hz, 3H).

<u>(3*R*,4*S*)-4-((*R*)-2-((4-amino-3-(4-phenoxyphenyl)-1H-pyrazolo[3,4-*d*]pyrimidin-1-yl)methyl)pyrrolidine-1-carbonyl)-3-isopropyloxetan-2-one (**3b**)</u>

In an oven dried 1-dram vial, (2*S*,3*R*)-3-isopropyl-4-oxooxetane-2-carboxylic acid (1.4 mg, 1.0 Eq, 8.9 μmol) and HATU (6.5 mg, 1.9 Eq, 0.017 mmol) were dissolved in DMF (100 μL) and the mixture was cooled to 0 ˚C. To this, 2,6-lutidine (5.4 mg, 5.8 μL, 5.7 Eq, 0.050 mmol) was added and the solution was allowed to stir on ice for 30 minutes. (*R*)-2-((4-amino-3-(4-phenoxyphenyl)-1H-pyrazolo[3,4-*d*]pyrimidin-1-yl)methyl)pyrrolidin-1-ium 2,2,2-trifluoroacetate (5.5 mg, 1.2 Eq, 0.011 mmol) was then added to the reaction and mixture was stirred at room temperature for 30 minutes. The reaction was diluted in DCM (1 mL) and washed with saturated sodium bicarbonate (2 mL). The organic layer was extracted DCM (3 x 2 mL). The combined organic layers were washed with brine (2 mL), dried over sodium sulfate, and concentrated. The residue was purified by reverse-phase C18 HPLC. Elution was performed with a linear gradient of 10–95% acetonitrile–water + 0.1% formic acid, and the product-containing fractions were lyophilized to afford the **3b** as a white solid (1.5 mg, 0.0089 mmol, 32%).

**HRMS (ESI):** calculated for $C_{29}H_{31}N_6O_4^+$ [M+H]$^+$ 527.2401, found 527.2405.

**$^1$H NMR (500 MHz, CDCl$_3$):** (mixture of rotamers, 1:0.4 major:minor) δ / ppm = 8.36 (s, 1.4H), 7.64 (d, major + minor rotamer, *J* = 8.6, 2.8H), 7.42 – 7.36 (m, major + minor rotamer, 2.8H), 7.22 – 7.13 (m, major + minor rotamer, 4.2H), 7.12 – 7.06 (m, major + minor rotamer, 2.8H), 5.60 (br s, major + minor rotamer, 2.4H), 4.90 (d, minor rotamer, *J* = 4.0 Hz, 0.4H), 4.83 (dd, major rotamer, *J* = 13.8, 5.5 Hz, 1H), 4.69 – 4.64 (m, major + minor 1.4H), 4.65 (d, major rotamer, *J* = 4.3 Hz, 1H), 4.61 (dd, major rotamer, *J* = 13.9, 4.0 Hz, 1H), 4.50 – 4.47 (m, minor rotamer, 0.8H), 3.99 (dd, minor rotamer, *J* = 7.7, 4.1 Hz, 0.4H), 3.93 (dd, major rotamer, *J* = 8.2, 4.2 Hz, 1H), 3.66 – 3.59 (m, minor rotamer, 0.4H), 3.53 – 3.38 (m, major + minor rotamer, 2.4H), 2.17 – 2.13 (m, major + minor, 1.4 H), 2.05 – 1.96 (m, major + minor rotamer, 2.8H), 1.85 – 1.80 (m, minor rotamer, 1.4H), 1.68 – 1.64 (m, major rotamer, 1.4H), 1.11 (d, *J* = 6.7 Hz, 3H), 1.07 (d, major + minor rotamer, *J* = 6.7 Hz, 4.2H), 1.00 (d, *J* = 6.7 Hz, 1.2H).

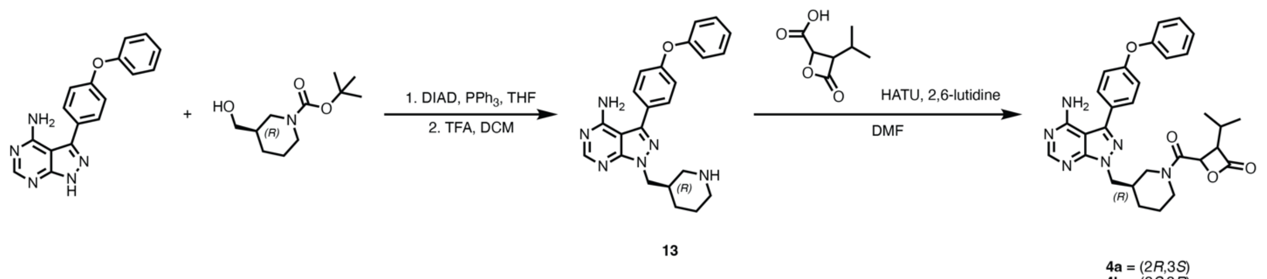

<u>(*R*)-3-((4-amino-3-(4-phenoxyphenyl)-1H-pyrazolo[3,4-*d*]pyrimidin-1-yl)methyl)piperidin-1-ium 2,2,2-trifluoroacetate (**13**)</u>

In an oven-dried 20 mL scintillation vial, 3-(4-phenoxyphenyl)-1H-pyrazolo[3,4-*d*]pyrimidin-4-amine (115.1 mg, 1.000 Eq, 379.5 µmol), tert-butyl (*R*)-3-(hydroxymethyl)piperidine-1-carboxylate (99.9 mg, 1.22 Eq, 464 µmol), and triphenylphosphine (106.6 mg, 1.071 Eq, 406.4 µmol) were dissolved in anhydrous THF (3.8 mL). Diisopropyl azodicarboxylate (81 mg, 78 µL, 1.1 Eq, 0.40 mmol) was added slowly and the reaction was stirred at room temperature for 16 h. The reaction was concentrated and purified by flash column chromatography in a gradient from 0-10% MeOH in DCM. The resulting oil was dissolved in DCM (2 mL) and TFA (2 mL) and allowed to sit for 3 h. The volatiles were removed by rotary evaporation and the residue was triturated with diethyl ether to afford the **13** as a white solid (169.6 mg, 0.3400 mmol, 97%).

**HRMS (ESI):** calculated for $C_{23}H_{25}N_6O^+$ $[M+H]^+$ 401.2085, found 401.2102.

**$^1$H NMR (500 MHz, $CDCl_3$):** δ / ppm = 8.26 (s, 1H), 7.57 (d, *J* = 8.4 Hz, 2H), 7.45 – 7.38 (m, 2H), 7.25 – 7.18 (m, 1H), 7.17 (d, *J* = 8.4 Hz, 2H), 7.12 – 7.07 (m, 2H), 6.32 (br s, 1H), 4.51 – 4.34 (m, 2H), 3.45 – 3.37 (m, 2H), 2.84 – 2.79 (m, 3H), 2.64 – 2.59 (m, 1H), 1.97 (d, *J* = 13.2 Hz, 2H), 1.91 – 1.79 (m, 1H), 1.37 – 1.30 (m, 1H).

<u>(3*S*,4*R*)-4-((*R*)-3-((4-amino-3-(4-phenoxyphenyl)-1H-pyrazolo[3,4-*d*]pyrimidin-1-yl)methyl)piperidine-1-carbonyl)-3-isopropyloxetan-2-one (**4a**)</u>

In a 1-dram vial, (2*S*,3*R*)-3-isopropyl-4-oxooxetane-2-carboxylic acid (5.22 mg, 1.90 Eq, 0.0330 mmol) and HATU (20.0 mg, 3.02 Eq, 0.0526 mmol) were dissolved in DMF (0.1 mL). To this solution, 2,6-dimethylpyridine (19.6 mg, 21.1 µL, 5.52 Eq, 0.182 mmol) was added and the mixture was cooled to 0 ˚C and stirred on ice for 20 min. To this solution, a mixture of **13** (7.7 mg, 1.0 Eq, 0.018 mmol) and 2,6-dimethylpyridine (7.4 mg, 8.0 µL, 3.9 Eq, 0.069 mmol) in DMF (0.1 mL) was added and the reaction was allowed to warm to room temperature and stir for 1 h. The reaction mixture was diluted in DCM (2 mL) and washed with 1x saturated sodium bicarbonate. The organic layer was filtered through a sodium sulfate, concentrated and the residue was purified by reverse-phase C18 HPLC. Elution was performed with a linear gradient of 10–95% acetonitrile–water + 0.1% formic acid, and the product-containing fractions were lyophilized to afford the **4a** as a white solid (1.8 mg, 0.015 mmol, 22%).

**HRMS (ESI):** calculated for $C_{30}H_{33}N_6O_4^+$ $[M+H]^+$ 541.2558, found 541.2581

**$^1$H NMR (600 MHz, $CDCl_3$):** (mixture of rotamers, 1:0.7 major:minor) δ / ppm = 8.35 (s, major rotamer, 1H), 8.33 (s, minor rotamer, 1H), 7.68 – 7.62 (m, major + minor rotamer, 3.4H), 7.41 – 7.38 (m, major + minor rotamer, 0.8 Hz, 3.4H), 7.21 – 7.13 (m, major + minor rotamer, 5.1H), 7.11 – 7.06 (m, major + minor rotamer, 3.4H), 5.98 (br s, major + minor rotamer, 2H), 4.78 (d, major rotamer, *J* = 4.2 Hz, 1H), 4.74 (d, minor rotamer, *J* = 4.1 Hz, 0.7H), 4.42 – 4.34 (m, major + minor rotamer, 3.4H), 4.34 – 4.28 (m, minor rotamer, 0.7H), 4.09 (dt, major rotamer, *J* = 12.4, 3.8 Hz, 1H), 3.95 (dd, minor rotamer, *J* = 8.1, 4.2 Hz, 0.7H), 3.90 (dd, major rotamer, *J* = 8.3, 4.2 Hz, 1H), 3.76 (d, minor rotamer, *J* = 13.9 Hz, 0.7H),

3.70 (dd, major rotamer, *J* = 13.6, 3.6 Hz, 1H), 3.23 – 3.15 (m, minor rotamer, 0.7H), 3.13 (dd, major rotamer, *J* = 13.5, 9.0 Hz, 1H), 3.07 (td, major rotamer, *J* = 11.8, 3.3 Hz, 1H), 2.81 (dt, minor rotamer, *J* = 14.4, 9.6 Hz, 0.7H), 2.39 – 2.31 (m, major + minor rotamer, 1.7H), 2.17 – 2.09 (m, major + minor rotamer, 1.7H), 1.98 – 1.91 (m, major + minor rotamer, 2.7H), 1.91 – 1.84 (m, major + minor rotamer, 1.7H), 1.81 – 1.75 (m, minor rotamer, 1.4H), 1.61 – 1.54 (m, minor rotamer, 1.4H), 1.50 (m, major + minor rotamer, 1.7H), 1.45 – 1.34 (m, major rotamer, 1H), 1.10 (d, minor rotamer, *J* = 6.7 Hz, 2.1H), 1.04 (d, minor rotamer, *J* = 6.7 Hz, 2.1H), 1.01 (d, major rotamer, *J* = 6.7 Hz, 3H), 0.95 (d, major rotamer, *J* = 6.7 Hz, 3H).

(3*R*,4*S*)-4-((*R*)-3-((4-amino-3-(4-phenoxyphenyl)-1H-pyrazolo[3,4-*d*]pyrimidin-1-yl)methyl)piperidine-1-carbonyl)-3-isopropyloxetan-2-one (**4b**)

In a 1-dram vial, (2*R*,3*S*)-3-isopropyl-4-oxooxetane-2-carboxylic acid (7.38 mg, 1.20 Eq, 0.0466 mmol) and HATU (20.0 mg, 1.35 Eq, 0.0526 mmol) were dissolved in DMF (0.1 mL). To this solution, 2,6-dimethylpyridine (12.5 mg, 13.5 µL, 3.00 Eq, 0.117 mmol) was added and the mixture was cooled to 0 ˚C and stirred on ice for 20 min. To this solution, a mixture of **13** (20.0 mg, 1.00 Eq, 0.0389 mmol) and 2,6-dimethylpyridine (12.5 mg, 13.5 µL, 3.00 Eq, 0.117 mmol) in DMF (0.1 mL) was added and the reaction was allowed to warm to room temperature and stir for 1 h. The reaction mixture was diluted in DCM (2 mL) and washed with 1x saturated sodium bicarbonate. The organic layer was filtered through a sodium sulfate, concentrated and the residue was purified by reverse-phase C18 HPLC. Elution was performed with a linear gradient of 10–95% acetonitrile–water + 0.1% formic acid, and the product-containing fractions were lyophilized to afford the **4a** as a white solid (5.0 mg, 0.039 mmol, 24%).

**HRMS (ESI)**: calculated for $C_{30}H_{33}N_6O_4^+$ $[M+H]^+$ 541.2558, found 541.2569.

**$^1$H NMR (600 MHz, $CDCl_3$):** (mixture of rotamers, major:minor 1:0.8) δ / ppm = 8.36 (s, minor rotamer, 0.8H), 8.33 (s, major rotamer, 1H), 7.68 – 7.62 (m, major + minor rotamer, 3.6H), 7.43 – 7.36 (m, major + minor rotamer, 3.6H), 7.21 – 7.13 (m, major + minor rotamer, 5.4H), 7.11 – 7.06 (m, major + minor rotamer, 3.6H), 5.92 (br s, 2H), 4.74 (d, major rotamer, *J* = 4.2 Hz, 1H), 4.68 (d, minor rotamer, *J* = 4.2 Hz, 0.8H), 4.43 – 4.28 (m, major + minor rotamer, 5.4H), 3.98 (dd, major rotamer, *J* = 8.2, 4.2 Hz, 1H), 3.89 (dd, minor rotamer, *J* = 8.0, 4.2 Hz, 0.8H), 3.86 – 3.78 (m, major + minor rotamer, 1.8H), 3.12 – 3.03 (m, major + minor rotamer, 1.8H), 2.91 – 2.84 (m, minor rotamer, 0.8H), 2.69 (dd, major rotamer, *J* = 13.2, 10.7 Hz, 1H), 2.39 – 2.31 (m, major + minor rotamer, 2.6H), 2.18 – 2.09 (m, major rotamer, 1H), 1.95 – 1.77 (m, major + minor rotamer, 3.6H), 1.63 – 1.50 (m, major + minor rotamer, 1.8H), 1.45 – 1.31 (m, major + minor rotamer, 1.8H), 1.10 (d, major rotamer, *J* = 6.7 Hz, 3H), 1.03 (d, major rotamer, *J* = 6.7 Hz, 2.4H), 1.00 (d, minor rotamer, *J* = 6.7 Hz, 2.4H), 0.94 (d, minor rotamer, *J* = 6.7 Hz, 2.1H).

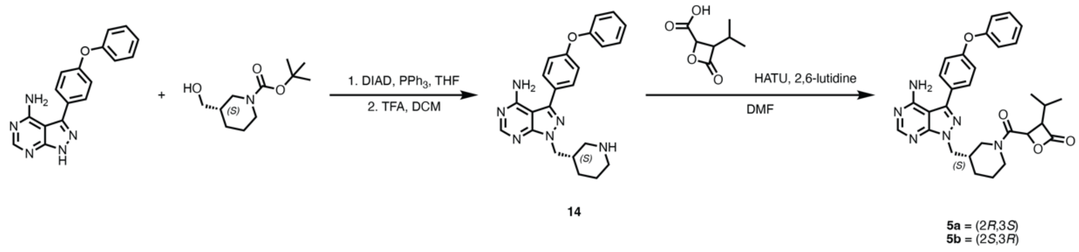

(*S*)-3-((4-amino-3-(4-phenoxyphenyl)-1H-pyrazolo[3,4-*d*]pyrimidin-1-yl)methyl)piperidin-1-ium 2,2,2-trifluoroacetate (**14**)

In an oven-dried 20 mL scintillation vial, 3-(4-phenoxyphenyl)-1H-pyrazolo[3,4-*d*]pyrimidin-4-amine (99.2 mg, 1.00 Eq, 0.327 mmol), tert-butyl (*S*)-3-(hydroxymethyl)piperidine-1-carboxylate (99.2 mg, 1.41 Eq, 0.461 mmol), and triphenylphosphine (109.6 mg, 1.280 Eq, 0.4179 mmol) were dissolved in anhydrous THF (3.3 mL). Diisopropyl azodicarboxylate (81 mg, 78 µL, 1.2 Eq, 0.40 mmol) was added slowly and the reaction was stirred at room temperature for 16 h. The reaction was concentrated and purified by flash column chromatography in a gradient from 0-10% MeOH in DCM. The resulting oil was dissolved in DCM (2 mL) and TFA (2 mL) and allowed to sit for 3 h. The volatiles were removed by rotary evaporation and the residue was triturated with diethyl ether to afford the **14** as a white solid (99.1 mg, 0.327 mmol, 61%).

**HRMS (ESI):** calculated for $C_{23}H_{25}N_6O^+$ $[M+H]^+$ 401.2085, found 401.2103.

**$^1$H NMR (500 MHz, CDCl$_3$):** δ / ppm = 8.24 (s, 1H), 7.57 (d, *J* = 8.5 Hz, 2H), 7.42 (t, *J* = 7.8 Hz, 2H), 7.21 (t, *J* = 7.4 Hz, 1H), 7.16 (d, *J* = 8.5 Hz, 2H), 7.09 (d, *J* = 7.6 Hz, 2H), 6.32 (s, 1H), 4.43 (qd, *J* = 14.3, 6.3 Hz, 2H), 3.38 (dd, *J* = 27.4, 12.4 Hz, 2H), 2.81 – 2.77 (m, 2H), 2.63 – 2.59 (m, 1H), 1.98 – 1.82 (m, 3H), 1.38 – 1.27 (m, 1H).

(3*S*,4*R*)-4-((*S*)-3-((4-amino-3-(4-phenoxyphenyl)-1H-pyrazolo[3,4-*d*]pyrimidin-1-yl)methyl)piperidine-1-carbonyl)-3-isopropyloxetan-2-one (**5a**)

In a 1-dram vial, (2*S*,3*R*)-3-isopropyl-4-oxooxetane-2-carboxylic acid (5.22 mg, 1.00 Eq, 0.0330 mmol) and HATU (20.0 mg, 1.59 Eq, 0.0526 mmol) were dissolved in DMF (0.1 mL). To this solution, 2,6-dimethylpyridine (19.6 mg, 21.1 µL, 5.52 Eq, 0.182 mmol) was added and the mixture was cooled to 0 ˚C and stirred on ice for 20 min. To this solution, a mixture of **14** (9.1 mg, 1.0 Eq, 0.018 mmol) and 2,6-dimethylpyridine (7.4 mg, 8.0 µL, 3.9 Eq, 0.069 mmol) in DMF (0.1 mL) was added and the reaction was allowed to warm to room temperature and stir for 1 h. The reaction mixture was diluted in DCM (2 mL) and washed with 1x saturated sodium bicarbonate. The organic layer was filtered through a sodium sulfate, concentrated and the residue was purified by reverse-phase C18 HPLC. Elution was performed with a linear gradient of 10–95% acetonitrile–water + 0.1% formic acid, and the product-containing fractions were lyophilized to afford the **5a** as a white solid (2.8 mg, 0.018 mmol, 29%).

**HRMS (ESI):** calculated for $C_{30}H_{33}N_6O_4^+$ $[M+H]^+$ 541.2558, found 541.2561.

**$^1$H NMR (600 MHz, CDCl$_3$):** (mixture of rotamers, major:minor 1:0.8) δ / ppm = 8.36 (s, minor rotamer, 0.8H), 8.33 (s, major rotamer, 1H), 7.66 – 7.63 (m, major + minor rotamer, 3.6H), 7.41 – 7.37 (m, major + minor rotamer, 3.6H), 7.21 – 7.13 (m, major + minor rotamer, 5.4H), 7.11 – 7.06 (m, major + minor rotamer, 3.6H), 5.93 (br s, 3.6H), 4.74 (d, major rotamer, *J* = 4.2 Hz, 1H), 4.68 (d, minor rotamer, *J* = 4.2 Hz, 0.8H), 4.43 – 4.28 (m, major + minor rotamer, 5.4H), 3.98 (dd, major rotamer, *J* = 8.2, 4.2 Hz, 1H), 3.89 (dd, minor rotamer, *J* = 8.0, 4.2 Hz, 0.8H), 3.82 (t, major + minor rotamer, *J* = 12.5 Hz, 1.8H), 3.15 – 3.02 (m, major + minor rotamer, 1.8H), 2.91 – 2.83 (m, minor rotamer, 0.8H), 2.69 (dd, major rotamer, *J* = 13.2, 10.7 Hz, 1H), 2.38 – 2.32 (m, major + minor rotamer, 2.6H), 2.18 – 2.08 (m, major rotamer, 1H), 1.95 – 1.79 (m, major + minor rotamer, 3.6H), 1.63 – 1.47 (m, major + minor rotamer, 1.8H), 1.46 – 1.34 (m, major + minor rotamer, 1.8H), 1.10 (d, major rotamer, *J* = 6.7 Hz, 3H), 1.03 (d, major rotamer, *J* = 6.7 Hz, 3H), 1.00 (d, minor rotamer, *J* = 6.7 Hz, 2.4H), 0.94 (d, minor rotamer, *J* = 6.8 Hz, 2.4H).

(3*R*,4*S*)-4-((*S*)-3-((4-amino-3-(4-phenoxyphenyl)-1H-pyrazolo[3,4-*d*]pyrimidin-1-yl)methyl)piperidine-1-carbonyl)-3-isopropyloxetan-2-one (**5b**)

In a 1-dram vial, (2*R*,3*S*)-3-isopropyl-4-oxooxetane-2-carboxylic acid (4.42 mg, 1.33 Eq, 0.0279 mmol) and HATU (12.0 mg, 1.50 Eq, 0.0315 mmol) were dissolved in DMF (0.1 mL). To this solution, 2,6-dimethylpyridine (11 mg, 12 µL, 5.0 Eq, 0.11 mmol) was added and the mixture was cooled to 0 ˚C and stirred on ice for 20 min. To this solution, a mixture of **14** (10.8 mg, 1.00 Eq, 0.0210 mmol) and 2,6-dimethylpyridine (2.3 mg, 2.5 µL, 1.0 Eq, 0.022 mmol) in DMF (0.1 mL) was added and the reaction was allowed to warm to room temperature and stir for 1 h. The reaction mixture was diluted in DCM (2 mL) and washed with 1x saturated sodium bicarbonate. The organic layer was filtered through a sodium sulfate, concentrated and the residue was purified by reverse-phase C18 HPLC. Elution was performed with a linear gradient of 10–95% acetonitrile–water + 0.1% formic acid, and the product-containing fractions were lyophilized to afford the **5b** as a white solid (2.5 mg, 0.021 mmol, 22%).

**HRMS (ESI)**: calculated for $C_{30}H_{33}N_6O_4^+$ $[M+H]^+$ 541.2558, found 541.2575.

**$^1$H NMR (600 MHz, CDCl$_3$):** (mixture of rotamers, 1:0.7 major:minor) δ / ppm = 8.35 (s, major rotamer, 1H), 8.33 (s, minor rotamer, 1H), 7.67 – 7.63 (m, major + minor rotamer, 3.4H), 7.39 (t, major + minor rotamer, *J* = 7.8 Hz, 3.4H), 7.21 – 7.13 (m, major + minor rotamer, 5.1H), 7.11 – 7.06 (m, major + minor rotamer, 3.4H), 5.95 (br s, major + minor rotamer, 3.4H), 4.78 (d, major rotamer, *J* = 4.1 Hz, 1H), 4.74 (d, minor rotamer, *J* = 4.1 Hz, 0.7H), 4.42 – 4.34 (m, major + minor rotamer, 3.4H), 4.34 – 4.28 (m, minor rotamer, 0.7H), 4.10 – 4.07 (m, major rotamer, 1H), 3.95 (dd, minor rotamer, *J* = 8.3, 4.2 Hz, 0.7H), 3.90 (dd, major rotamer, *J* = 8.4, 4.2 Hz, 1H), 3.77 (d, minor rotamer, *J* = 13.9 Hz, 0.7H), 3.70 (d, major rotamer, *J* = 13.6 Hz, 1H), 3.23 – 3.06 (m, major + minor rotamer, 2.7H), 2.81 (dd, minor rotamer, *J* = 13.2, 10.1 Hz, 0.7H), 2.39 – 2.32 (m, major + minor rotamer, 1.7H), 2.15 – 2.09 (m, major + minor rotamer, 1.7H), 1.96 – 1.92 (m, major + minor rotamer, 2.7H), 1.88 – 1.86 (m, major + minor rotamer, 1.7H), 1.79 – 1.76 (m, minor rotamer, 1.4H), 1.59 – 1.56 (m, minor rotamer, 1.4H), 1.52 – 1.46 (m, major + minor rotamer, 1.7H), 1.41 – 1.35 (m, major rotamer, 1H), 1.10 (d, minor rotamer, *J* = 6.6 Hz, 2.1H), 1.04 (d, minor rotamer, *J* = 6.7 Hz, 2.1H), 1.01 (d, major rotamer, *J* = 6.6 Hz, 4H), 0.95 (d, major rotamer, *J* = 6.5 Hz, 3H).

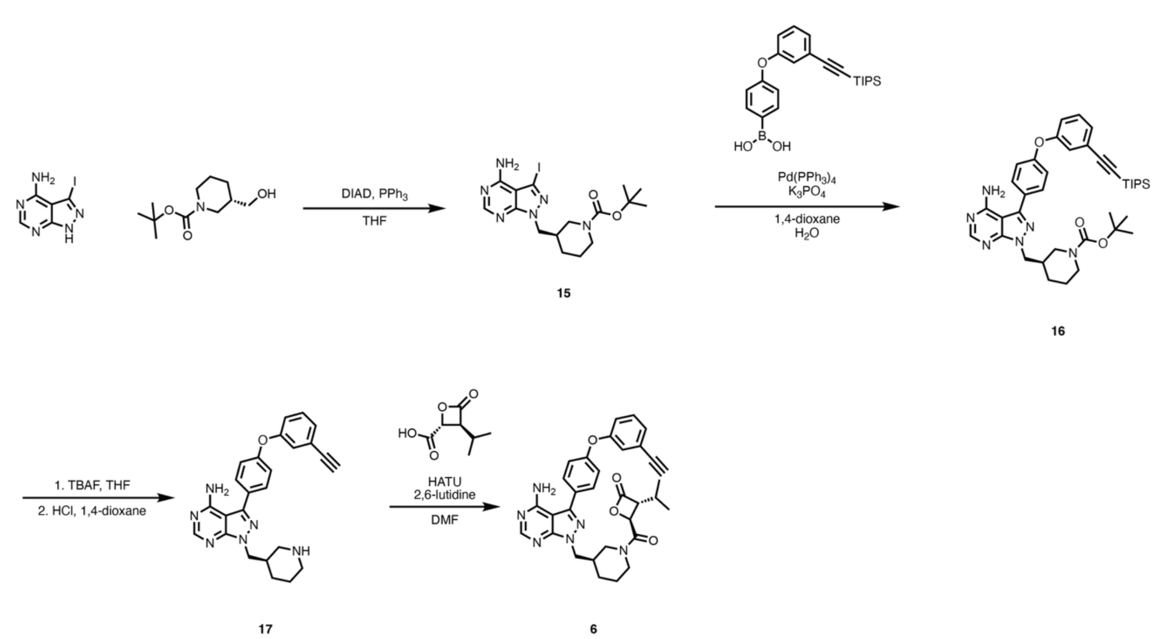

<u>(4-(3-((triisopropylsilyl)ethynyl)phenoxy)phenyl)boronic acid</u>

Synthesized according to literature procedures.[13]

<u>tert-butyl (*R*)-3-((4-amino-3-iodo-1H-pyrazolo[3,4-*d*]pyrimidin-1-yl)methyl)piperidine-1-carboxylate (**15**)</u>

tert-butyl (*R*)-3-(hydroxymethyl)piperidine-1-carboxylate (253.2 mg, 1.465 Eq, 1.176 mmol), 3-iodo-1H-pyrazolo[3,4-*d*]pyrimidin-4-amine (209.6 mg, 1.000 Eq, 803.0 µmol), and triphenylphosphine (297.5 mg, 1.413 Eq, 1.134 mmol) were added to an oven-dried vial and dissolved in THF (7 mL). Diisopropyl azodicarboxylate (250 mg, 240 µL, 1.54 Eq, 1.23 mmol) was added slowly and the reaction was stirred at room temperature for 16 h. The reaction was concentrated and the residue was purified by flash column chromatography (0-5% gradient of MeOH in DCM). The fractions containing product were combined to yield **15** as a yellow solid (275 mg, 0.803 mmol, 75%).

**HRMS (ESI):** calculated for $C_{16}H_{24}IN_6O_2^+$ $[M+H]^+$ 459.1000, found 459.1051.

**$^1$H NMR (500 MHz, $CDCl_3$):** δ / ppm = 8.30 (s, 1H), 6.30 (br s, 2H), 4.25 (d, *J* = 7.3 Hz, 2H), 3.86 (d, *J* = 13.4 Hz, 2H), 2.81 (ddd, *J* = 13.7, 10.9, 2.8 Hz, 1H), 2.71 – 2.68 (m, 1H), 2.24 – 2.21 (m, 1H), 1.71 – 1.65 (m, 2H), 1.44 – 1.42 (m, 1H), 1.39 (s, 9H), 1.24 - 1.17 (m, 1H).

<u>tert-butyl (*R*)-3-((4-amino-3-(4-(3-((triisopropylsilyl)ethynyl)phenoxy)phenyl)-1H-pyrazolo[3,4-*d*]pyrimidin-1-yl)methyl)piperidine-1-carboxylate (**16**)</u>

A 20-ml vial was charged with (4-(3-((triisopropylsilyl)ethynyl)phenoxy)phenyl)boronic acid (79.7 mg, 1.00 Eq, 0.0202 mmol), compound **15** (106.5 mg, 1.150 Eq, 0.2324 mmol) and a magnetic stir bar.1,4-Dioxane (1.5 mL) was added and the mixture was stirred briefly. To this solution, potassium phosphate (1.5 mL, 0.50 molar, 3.7 Eq, 0.75 mmol) was added as an aqueous solution. A stream of nitrogen was bubbled through the solution through a 22-gauge needle for 10 min.
Tetrakis(triphenylphosphine)palladium(0) (23 mg, 0.10 Eq, 0.020 mmol) was then added and the mixture

was heated to 100 ˚C for 18 h. The reaction mixture was cooled to 23 ˚C and then acidified with 1 N HCl (3 mL), until pH paper indicated a pH of ~5. The mixture was extracted with ethyl acetate (3 x 5 mL), and the combined organic layers were dried over sodium sulfate, filtered, and concentrated. The residue was purified by flash column chromatography (gradient of 0-10% MeOH in DCM) to afford the **16** as a brown oil which was used directly in the next step.

**$^1$H NMR (400 MHz, $CDCl_3$):** δ / ppm = δ 8.37 (s, 1H), 7.70 – 7.64 (m, 2H), 7.35 – 7.27 (m, 2H), 7.21 – 7.12 (m, 3H), 7.03 (dt, *J* = 7.4, 2.2 Hz, 1H), 5.57 (br s, 2H), 4.36 – 4.29 (m, 2H), 3.92 – 3.88 (m, 2H), 2.87 – 2.73 (m, 2H), 2.31 – 2.27 (m, 1H), 1.77 – 1.69 (m, 3H), 1.46 – 1.40 (m, 2H), 1.38 (s, 9H), 1.25 (t, *J* = 11.1 Hz, 2H), 1.12 (d, *J* = 1.8 Hz, 18H).

(*R*)-3-(4-(3-ethynylphenoxy)phenyl)-1-(piperidin-3-ylmethyl)-1H-pyrazolo[3,4-*d*]pyrimidin-4-amine (**17**)

Compound **16** was dissolved in THF (1 mL), and the solution was treated with 1 M TBAF (106 mg, 405 µL, 2.00 Eq, 0.405 mmol) in THF. The reaction was stirred for 1 hours at room temperature. The reaction was diluted with EtOAc (1 mL), washed with brine (3 x 2 mL), and dried over sodium sulfate in a packed glass pipette. The volatiles were removed to produce an orange oil which was then redissolved in DCM (0.8 mL). To this solution, 1 M HCl in 1,4-dioxane (0.8 mL) was added and the mixture was stirred at room temperature for 16 h. The volatiles were removed and the product was triturated out in diethyl ether and dried down. This residue was then resuspended in ethyl acetate (1 mL) and 1 N HCl (1 mL). The product-containing aqueous layer was separated and basified with 1 M NaOH until the pH was over 8. The product was extracted with ethyl acetate and dried down to yield **17** as a light brown solid (59.0 mg, 0.202 mmol, 69% over 2 steps).

**HRMS (ESI):** calculated for $C_{25}H_{25}N_6O^+$ $[M+H]^+$ 425.2084, found 425.2076.

**$^1$H NMR (500 MHz, $CDCl_3$):** δ / ppm = δ 8.36 (s, 1H), 7.67 (d, *J* = 8.6 Hz, 2H), 7.36 – 7.28 (m, 2H), 7.20 – 7.14 (m, 2H), 7.11 – 7.06 (m, 1H), 5.59 (br s, 2H), 4.35 (dd, *J* = 6.8, 3.6 Hz, 2H), 3.18 – 3.15 (m, 2H), 3.10 (s, 1H), 2.66 (td, *J* = 12.0, 3.1 Hz, 2H), 2.59 (t, *J* = 11.6 Hz, 1H), 2.46 – 2.42 (m, 1H), 1.87 – 1.85 (d, *J* = 13.5 Hz, 1H), 1.80 – 1.77 (m, 1H) 1.72 – 1.67 (m, 1H).

(3*R*,4*S*)-4-((*R*)-3-((4-amino-3-(4-(3-ethynylphenoxy)phenyl)-1H-pyrazolo[3,4-*d*]pyrimidin-1-yl)methyl)piperidine-1-carbonyl)-3-isopropyloxetan-2-one (**6**)

In an oven dried 1-dram vial, (2*S*,3*R*)-3-isopropyl-4-oxooxetane-2-carboxylic acid (4.47 mg, 1.20 Eq, 0.028.3 mmol) and HATU (10.7 mg, 1.19 Eq, 0.0283 mmol) were dissolved in DMF (0.2 mL). The solution was cooled to 0 ˚C and 2,6-lutidine (12.6 mg, 13.6 µL, 4.98 Eq, 0.118 mmol) was added slowly. The mixture was allowed to stir on ice for 20 minutes after which **17** (10.0 mg, 1.00 Eq, 0.0236 mmol) was added and the reaction was allowed to proceed at room temperature for 16 h. The reaction mixture was diluted with DCM (1 mL) and washed with saturated sodium bicarbonate (1 x 1 mL). The aqueous layer was extracted with DCM (3 x 2 mL). The combined organic layers were washed with brine (2 mL), dried over sodium sulfate, and concentrated. The residue was purified by reverse-phase C18 HPLC. Elution was performed with a linear gradient of 10–95% acetonitrile–water + 0.1% formic acid, and the product-containing fractions were lyophilized to afford the **6** as a white solid (2.9 mg, 0.024 mmol, 22%).

**HRMS (ESI):** calculated for $C_{32}H_{33}N_6O_4^+$ $[M+H]^+$ 565.2558, found 565.2538.

**$^1$H NMR (600 MHz, $CDCl_3$):** (mixture of rotamers, 1:0.8 major:minor) δ / ppm = 8.34 (s, minor rotamer, 0.8H), 8.30 (s, major rotamer, 1H), 7.67 (dd, major + minor rotamer, J = 8.5, 6.1 Hz, 3.6H), 7.35 – 7.32 (m, major + minor rotamer, 1.8H) 7.31 – 7.29 (m, major + minor rotamer, 1.8H), 7.20 – 7.13 (m, major + minor rotamer, 5.4H), 7.10 – 7.08 (m, major + minor rotamer, 1.8H), 6.07 (br s, major + minor rotamer, 1.8H), 4.74 (d, major rotamer, J = 4.2 Hz, 1H), 4.68 (d, minor rotamer, J = 4.2 Hz, 0.8H), 4.43 – 4.28 (m, 5.4H), 3.98 (dd, major rotamer, J = 8.3, 4.2 Hz, 1H), 3.90 (dd, minor rotamer, J = 8.0, 4.2 Hz, 0.8H), 3.83 (d, J = 12.2 Hz, 1.8H), 3.10 (s, minor rotamer, 0.8H), 3.10 (s, major rotamer, 1H), 3.13 – 3.03 (m, major + minor rotamer, 1.8H), 2.92 – 2.85 (m, minor rotamer, 0.8H), 2.70 (dd, J = 13.2, 10.6 Hz, 1H), 2.38 – 2.33 (m, major + minor rotamer, 1.8H), 2.16 – 2.09 (m, major rotamer, 1H), 1.96 – 1.91 (m, minor rotamer, 0.8H), 1.90 – 1.78 (m, major + minor rotamer, 3.6H), 1.63 – 1.48 (m, major + minor rotamer, 1.8H), 1.45 – 1.29 (m, major + minor rotamer, 1.8H), 1.10 (d, major rotamer, J = 6.7 Hz, 3H), 1.03 (d, major rotamer, J = 6.7 Hz, 3H), 1.00 (d, minor rotamer, J = 6.7 Hz, 2.4H), 0.95 (d, minor rotamer, J = 6.8 Hz, 2.4H).

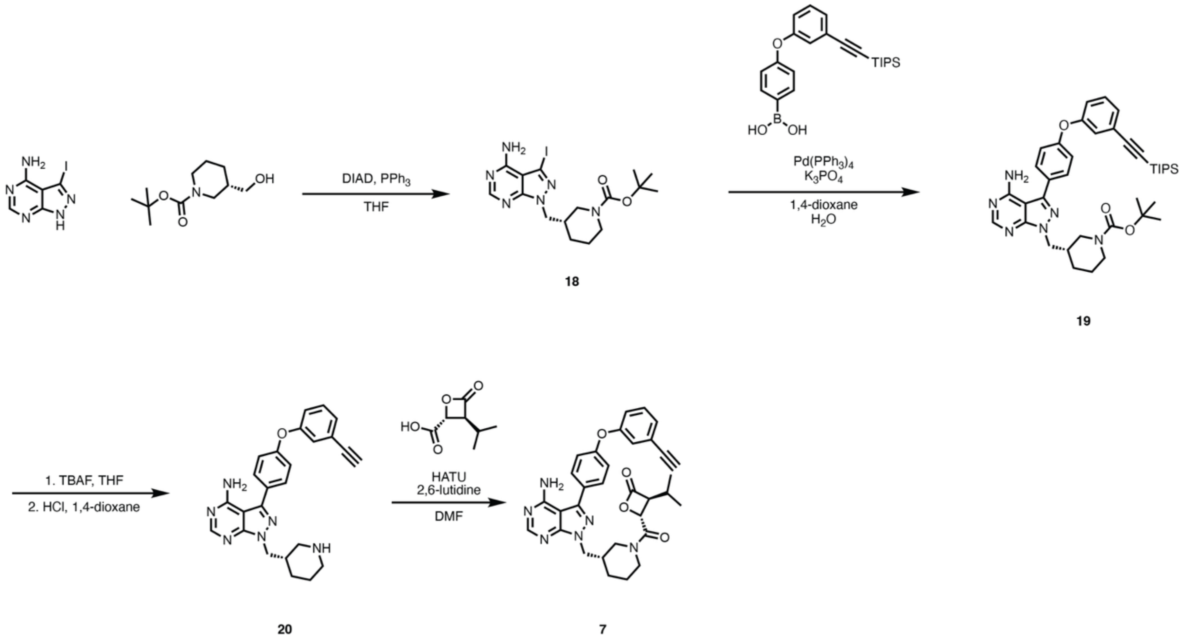

<u>tert-butyl (*S*)-3-((4-amino-3-iodo-1H-pyrazolo[3,4-*d*]pyrimidin-1-yl)methyl)piperidine-1-carboxylate (**18**)</u>

3-iodo-1H-pyrazolo[3,4-*d*]pyrimidin-4-amine (218.9 mg, 1.000 Eq, 838.6 μmol), triphenylphosphine (314.9 mg, 1.432 Eq, 1.201 mmol), and tert-butyl (*S*)-3-(hydroxymethyl)piperidine-1-carboxylate (256.3 mg, 1.420 Eq, 1.190 mmol) were added to an oven-dried vial and dissolved in THF (7 mL). Diisopropyl azodicarboxylate (250 mg, 240 μL, 1.47 Eq, 1.23 mmol) was added slowly and the reaction was stirred at RT overnight. reaction was stirred at room temperature for 16 h. The reaction was concentrated and the residue was purified by flash column chromatography (0-5% gradient of MeOH in DCM). The fractions containing product were combined to afford **18** as a yellow solid (271 mg, 0.839 mmol, 71%).

**HRMS (ESI):** calculated for $C_{16}H_{24}IN_6O_2^+$ [M+H]$^+$ 459.1000, found 459.1050.

**$^1$H NMR (500 MHz, CDCl$_3$):** δ / ppm = 8.33 (s, 1H), 5.87 (br s, 2H), 4.26 (dd, *J* = 7.3, 1.8 Hz, 2H), 3.87 (d, *J* = 13.3 Hz, 1H), 2.82 (ddd, *J* = 13.6, 11.1, 2.9 Hz, 1H), 2.71 – 2.68 (m, 1H), 2.24 – 2.21 (m, 1H), 1.71 – 1.67 (m, 2H), 1.44 – 1.42 (m, 1H), 1.39 (s, 9H), 1.24 - 1.17 (m, 1H).

tert-butyl (*S*)-3-((4-amino-3-(4-(3-((triisopropylsilyl)ethynyl)phenoxy)phenyl)-1H-pyrazolo[3,4-*d*]pyrimidin-1-yl)methyl)piperidine-1-carboxylate (**19**)

A 20-ml vial was charged with (4-(3-((triisopropylsilyl)ethynyl)phenoxy)phenyl)boronic acid (81.4 mg, 1.00 Eq, 206 μmol), **18** (103 mg, 1.09 Eq, 0.225 mmol) and a magnetic stir bar. 1,4-Dioxane (1.5 mL) was added and the mixture was stirred briefly. To this solution, 0.5 M potassium phosphate (1.5 mL, 3.7 Eq, 0.75 mmol) was added as an aqueous solution. A stream of nitrogen was bubbled through the solution through a 22-gauge needle for 10 min. Tetrakis(triphenylphosphine)palladium(0) (24 mg, 0.10 Eq, 0.021 mmol) was then added and the mixture was heated to 100 ˚C for 18 h. The reaction mixture was cooled to 23 ˚C and then acidified with 1 N HCl (3 mL), until pH paper indicated a pH of ~5. The mixture was extracted with ethyl acetate (3 x 5 mL) and the combined organic layers were dried over sodium sulfate, filtered, and concentrated. The residue was purified by flash column chromatography (gradient of 0-10% MeOH in DCM) to afford **19** as a brown oil which was used directly in future steps.

**$^1$H NMR (400 MHz, $CDCl_3$):** δ / ppm = δ 8.38 (s, 1H), 7.70 – 7.65 (m, *J* = 8.6 Hz, 2H), 7.35 – 7.27 (m, 2H), 7.21 – 7.12 (m, 3H), 7.03 (dt, *J* = 7.3, 2.3 Hz, 1H), 5.68 (br s, 2H), 4.34 – 4.32 (m, 2H), 3.91 – 3.87 (m, 2H), 2.91 – 2.60 (m, 2H), 2.31 – 2.28 (m, 1H), 1.80 – 1.67 (m, 3H), 1.39 (s, 11H), 1.24 (dd, *J* = 17.8, 7.2 Hz, 1H), 1.12 (d, *J* = 1.7 Hz, 20H).

(*S*)-3-(4-(3-ethynylphenoxy)phenyl)-1-(piperidin-3-ylmethyl)-1H-pyrazolo[3,4-*d*]pyrimidin-4-amine (**20**)

Compound **19** was dissolved in THF (1 mL) and the solution was treated with 1M TBAF in THF (106 mg, 405 μL, 2.00 Eq, 0.405 mmol). The reaction was stirred for 1 hour at room temperature. The reaction was diluted with EtOAc (1 mL), washed with brine (3 x 2 mL), and dried over sodium sulfate in a packed glass pipette. The volatiles were removed to produce an orange oil which was then redissolved in DCM (0.8 mL). To this solution, 1 M HCl in 1,4-dioxanes (0.8 mL, 4 Eq., 0.8 mmol) was added and the mixture was stirred at room temperature for 16 h. The volatiles were removed and the product was triturated out in diethyl ether and dried down. This residue was then resuspended in ethyl acetate (1 mL) and 1 N HCl (1 mL). The product-containing aqueous layer was separated and basified with 1 M NaOH until the pH was over 8. The product was extracted with ethyl acetate and dried down to yield **20** as a light brown solid (57.8 mg, 0.208 mmol, 60% over 2 steps).

**HRMS (ESI):** calculated for $C_{25}H_{25}N_6O^+$ $[M+H]^+$ 425.2084, found 425.2076.

**$^1$H NMR (500 MHz, $CDCl_3$):** δ / ppm = 8.38 (s, 1H), 7.67 (d, *J* = 8.6 Hz, 2H), 7.37 – 7.27 (m, 2H), 7.19 – 7.14 (m, 3H), 7.08 (ddd, *J* = 8.1, 2.6, 1.2 Hz, 1H), 5.50 (br s, 2H), 4.32 (qd, *J* = 13.8, 7.3 Hz, 2H), 3.10 (s, 1H), 3.02 – 2.94 (m, 2H), 2.60 (td, *J* = 11.7, 2.9 Hz, 1H), 2.48 (dd, *J* = 12.2, 10.0 Hz, 1H), 2.29 – 2.24 (m, 1H), 1.85 – 1.66 (m, 2H), 1.51 – 1.43 (m, 1H), 1.31 – 1.23 (m, 1H).

(3*S*,4*R*)-4-((*S*)-3-((4-amino-3-(4-(3-ethynylphenoxy)phenyl)-1H-pyrazolo[3,4-*d*]pyrimidin-1-yl)methyl)piperidine-1-carbonyl)-3-isopropyloxetan-2-one (**7**)

In an oven dried 1-dram vial, (2*R*,3*S*)-3-isopropyl-4-oxooxetane-2-carboxylic acid (4.47 mg, 1.20 Eq, 0.0283 mmol) and HATU (10.7 mg, 1.19 Eq, 0.0281 mmol) were dissolved in DMF (0.2 mL). The solution was cooled to 0 ˚C and 2,6-lutidine (12.6 mg, 13.6 μL, 4.98 Eq, 0.118 mmol) was added slowly. The mixture was allowed to stir on ice for 20 minutes after which **20** (10.0 mg, 1.00 Eq, 0.0236 mmol) was added and the reaction was allowed to proceed at room temperature for 16 h. The reaction mixture was diluted with DCM (1 mL) and washed with saturated sodium bicarbonate (1 x 1 mL). The aqueous layer was extracted with DCM (3 x 2 mL). The combined organic layers were washed with brine (2 mL), dried over sodium sulfate, and concentrated. The residue was purified by reverse-phase C18 HPLC.

Elution was performed with a linear gradient of 10–95% acetonitrile–water + 0.1% formic acid, and the product-containing fractions were lyophilized to afford the **7** as a white solid (3.2 mg, 0.023 mmol, 24%).

**HRMS (ESI)**: calculated for $C_{32}H_{33}N_6O_4^+$ $[M+H]^+$ 565.2558, found 565.2525.

**$^1$H NMR (600 MHz, CDCl$_3$):** (mixture of rotamers, 1:0.8 major:minor) δ / ppm = 8.35 (s, minor rotamer, 0.8H), 8.32 (s, major rotamer, 1H), 7.68 – 7.66 (m, major + minor rotamer, 3.6H), 7.36 – 7.32 (m, major + minor rotamer, 1.8H) 7.31 – 7.28 (m, major + minor rotamer, 1.8H), 7.20 – 7.15 (m, major + minor rotamer, 5.4H), 7.10 – 7.08 (m, major + minor rotamer, 1.8H), 6.01 (br s, major + minor rotamer, 1.8H), 4.74 (d, major rotamer, J = 4.2 Hz, 1H), 4.68 (d, minor rotamer, J = 4.2 Hz, 0.8H), 4.43 – 4.28 (m, 5.4H), 3.98 (dd, major rotamer, J = 8.3, 4.2 Hz, 1H), 3.90 (dd, minor rotamer, J = 8.0, 4.2 Hz, 0.8H), 3.83 (d, J = 12.2 Hz, 1.8H), 3.10 (s, minor rotamer, 0.8H), 3.10 (s, major rotamer, 1H), 3.11 – 3.04 (m, major + minor rotamer, 1.8H), 2.90 – 2.85 (m, minor rotamer, 0.8H), 2.70 (dd, J = 13.2, 10.6 Hz, 1H), 2.38 – 2.33 (m, major + minor rotamer, 1.8H), 2.16 – 2.09 (m, major rotamer, 1H), 1.96 – 1.91 (m, minor rotamer, 0.8H), 1.90 – 1.80 (m, major + minor rotamer, 3.6H), 1.63 – 1.48 (m, major + minor rotamer, 1.8H), 1.45 – 1.29 (m, major + minor rotamer, 1.8H), 1.10 (d, major rotamer, J = 6.7 Hz, 3H), 1.03 (d, major rotamer, J = 6.7 Hz, 3H), 1.00 (d, minor rotamer, J = 6.7 Hz, 2.4H), 0.95 (d, minor rotamer, J = 6.8 Hz, 2.4H).

(*R*)-1-(3-(4-amino-3-(4-(3-ethynylphenoxy)phenyl)-1H-pyrazolo[3,4-*d*]pyrimidin-1-yl)piperidin-1-yl)prop-2-en-1-one (**8**, **ibrutinib-alkyne**)

Synthesized according to literature procedure.[13]

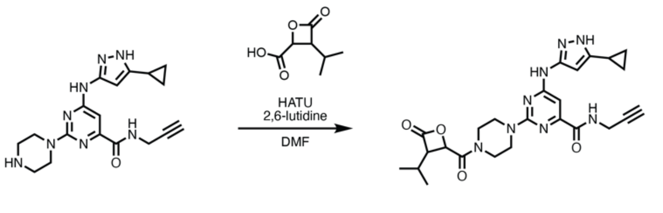

**9a** = (2*R*,3*S*)
**9b** = (2*S*,3*R*)

6-((5-cyclopropyl-1H-pyrazol-3-yl)amino)-2-(4-((2*R*,3*S*)-3-isopropyl-4-oxooxetane-2-carbonyl)piperazin-1-yl)-N-(prop-2-yn-1-yl)pyrimidine-4-carboxamide (**9a**)

In an oven dried 1-dram vial, (2*R*,3*S*)-3-isopropyl-4-oxooxetane-2-carboxylic acid (1.4 mg, 1.0 Eq, 0.0089 mmol) and HATU (6.5 mg, 1.9 Eq, 0.017 mmol) were dissolved in DMF (0.1 mL). The solution was cooled to 0 ˚C and 2,6-lutidine (5.4 mg, 5.8 µL, 5.7 Eq, 0.050 mmol) was added slowly. The mixture was allowed to stir on ice for 20 minutes after which 4-(4-((5-cyclopropyl-1H-pyrazol-3-yl)amino)-6-(prop-2-yn-1-ylcarbamoyl)pyrimidin-2-yl)piperazin-1-ium 2,2,2-trifluoroacetate (8.5 mg, 2.0 Eq, 18 µmol) (8.5 mg, 2.0 Eq, 0.018 mol) was added and the reaction was allowed to proceed at room temperature for 16 h. The reaction mixture was diluted with DCM (1 mL) and washed with saturated sodium bicarbonate (1 x 1 mL). The aqueous layer was extracted with DCM (3 x 2 mL). The combined organic layers were washed with brine (2 mL), dried over sodium sulfate, and concentrated. The residue was purified by reverse-phase C18 HPLC. Elution was performed with a linear gradient of 10–95% acetonitrile–water + 0.1% formic acid, and the product-containing fractions were lyophilized to afford **9a** as a white solid (0.5 mg, 0.0089 mmol, 10%).

**HRMS (ESI):** calculated for $C_{25}H_{31}N_8O_4^+$ $[M+H]^+$ 507.2463, found 507.2480.

**$^{1}$H NMR (500 MHz, $CDCl_3$):** δ / ppm = 7.94 (t, *J* = 5.7 Hz, 1H), 6.92 (s, 1H), 6.15 (s, 1H), 4.83 (d, *J* = 4.2 Hz, 1H), 4.24 (dd, *J* = 5.8, 2.6 Hz, 2H), 4.10 – 4.07 (m, 1H) 4.07 (dd, *J* = 8.2, 4.1 Hz, 1H), 3.97 – 3.90 (m, 1H), 3.77 – 3.66 (m 4H), 3.61 – 3.51 (m, 2H), 2.28 (t, *J* = 2.6 Hz, 1H), 2.23 – 2.14 (m, 1H), 1.96 – 1.91 (m, 1H), 1.15 (d, *J* = 6.6 Hz, 3H), 1.09 (d, *J* = 6.7 Hz, 3H), 1.04 – 0.96 (m, 2H), 0.75 (dt, *J* = 6.7, 4.7 Hz, 2H).

6-((5-cyclopropyl-1H-pyrazol-3-yl)amino)-2-(4-((2*S*,3*R*)-3-isopropyl-4-oxooxetane-2-carbonyl)piperazin-1-yl)-N-(prop-2-yn-1-yl)pyrimidine-4-carboxamide (**9b**)

In an oven dried 1-dram vial, (2*S*,3*R*)-3-isopropyl-4-oxooxetane-2-carboxylic acid (1.4 mg, 1.0 Eq, 0.0089 mmol) and HATU (6.5 mg, 1.9 Eq, 0.017 mmol) were dissolved in DMF (0.1 mL). The solution was cooled to 0 ˚C and 2,6-lutidine (5.4 mg, 5.8 µL, 5.7 Eq, 0.050 mmol) was added slowly. The mixture was allowed to stir on ice for 20 minutes after which 4-(4-((5-cyclopropyl-1H-pyrazol-3-yl)amino)-6-(prop-2-yn-1-ylcarbamoyl)pyrimidin-2-yl)piperazin-1-ium 2,2,2-trifluoroacetate (8.5 mg, 2.0 Eq, 18 µmol) (8.5 mg, 2.0 Eq, 0.018 mmol) was added and the reaction was allowed to proceed at room temperature for 16 h. The reaction mixture was diluted with DCM (1 mL) and washed with saturated sodium bicarbonate (1 x 1 mL). The aqueous layer was extracted with DCM (3 x 2 mL). The combined organic layers were washed with brine (2 mL), dried over sodium sulfate, and concentrated. The residue was purified by reverse-phase C18 HPLC. Elution was performed with a linear gradient of 10–95% acetonitrile–water + 0.1% formic acid, and the product-containing fractions were lyophilized to afford **9b** as a white solid (0.5 mg, 0.0089 mmol, 10%).

**HRMS (ESI):** calculated for $C_{25}H_{31}N_8O_4^+$ $[M+H]^+$ 507.2463, found 506.2471.

**$^{1}$H NMR (500 MHz, $CDCl_3$):** δ / ppm = 7.95 (t, *J* = 5.6 Hz, 1H), 6.91 (s, 1H), 6.17 (s, 1H), 4.83 (d, *J* = 4.2 Hz, 1H), 4.24 (dd, *J* = 5.7, 2.6 Hz, 2H), 4.11 – 4.07 (m, 1H), 4.07 (dd, *J* = 8.2, 4.2 Hz, 1H), 3.98 – 3.92 (m, 1H), 3.78 – 3.67 (m, 4H), 3.61 – 3.51 (m, 2H), 2.28 (t, *J* = 2.6 Hz, 1H), 2.25 – 2.15 (m, 1H), 1.94 – 1.99 (m, 1H), 1.15 (d, *J* = 6.7 Hz, 3H), 1.09 (d, *J* = 6.7 Hz, 3H), 1.07 – 0.96 (m, 2H), 0.76 (dt, *J* = 6.7, 4.6 Hz, 2H).

$^{1}$H-NMR (500 MHz, $CDCl_3$) of **1a**

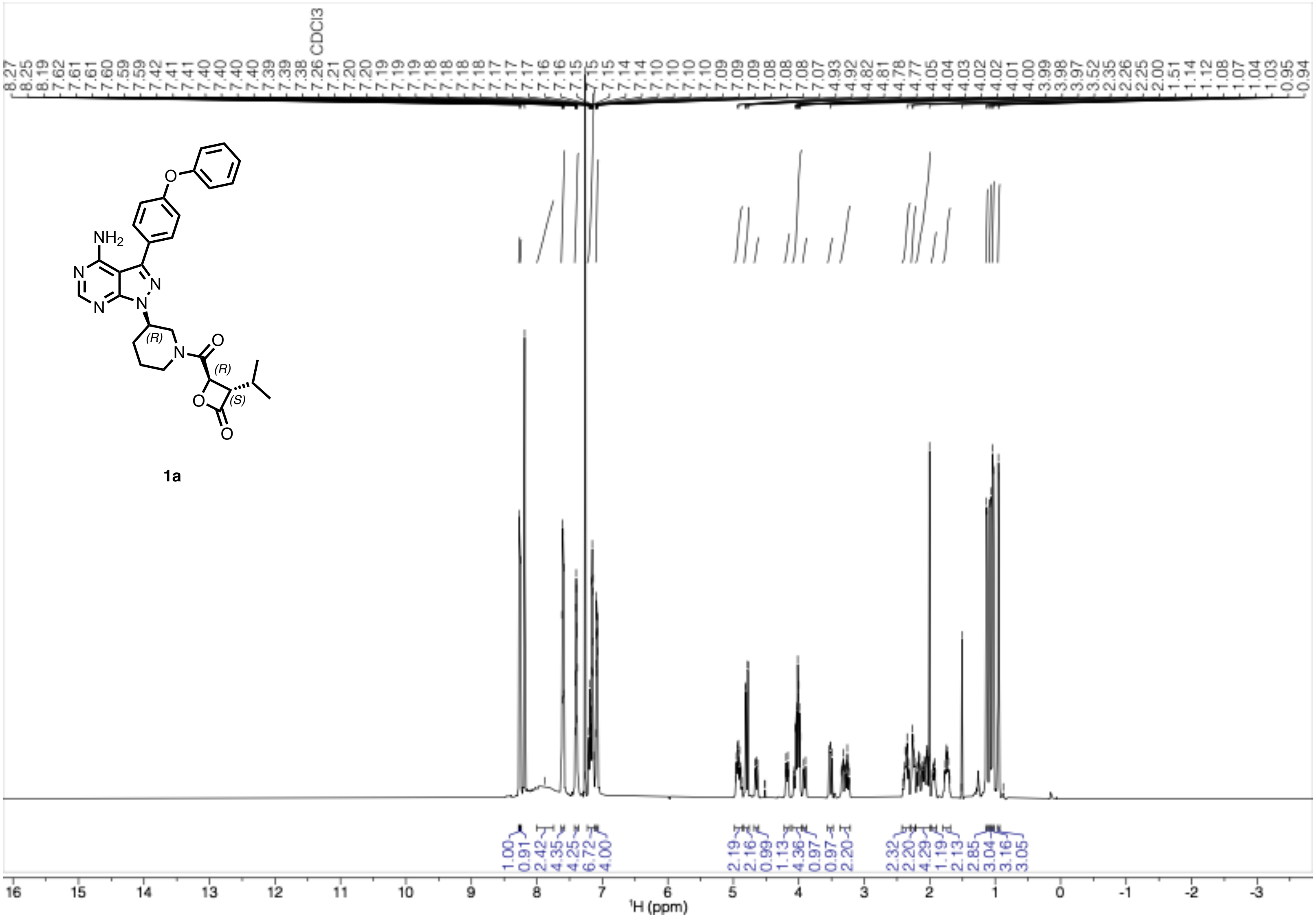

$^{1}H$-NMR (500 MHz, $CDCl_3$) of **1b**

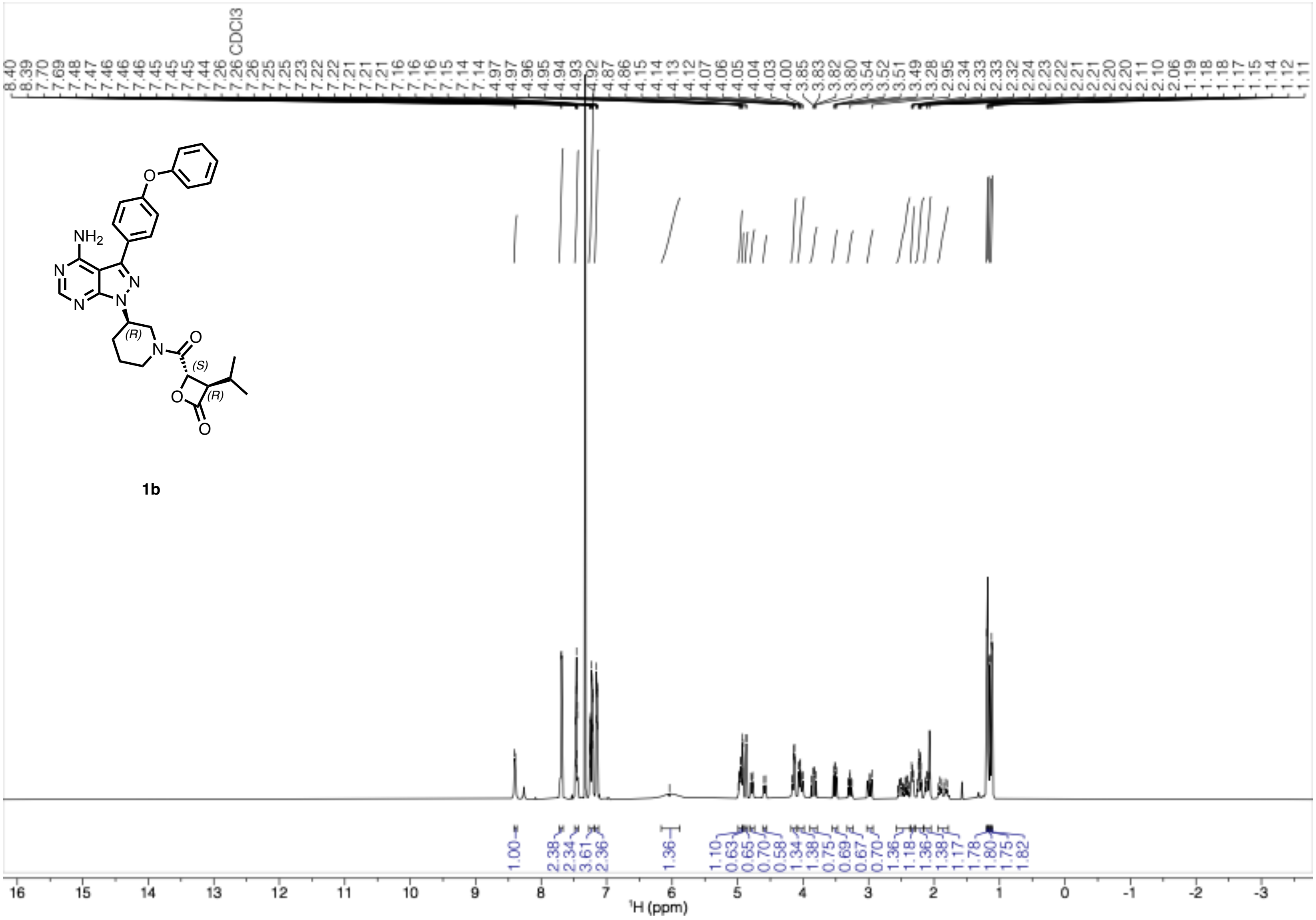

[1]H-NMR (500 MHz, CDCl3) of **2a**

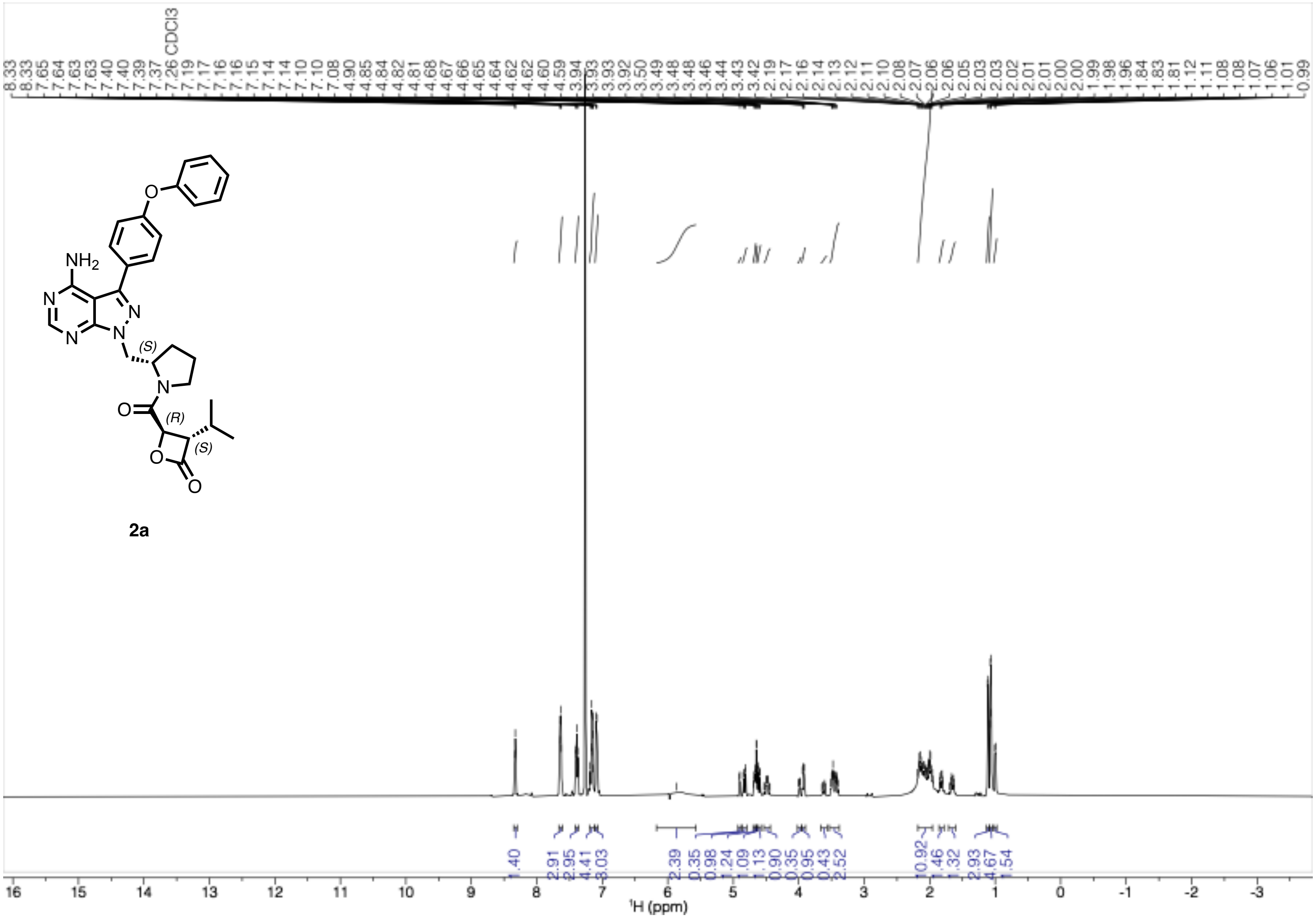

[1]H-NMR (500 MHz, $CDCl_3$) of **2b**

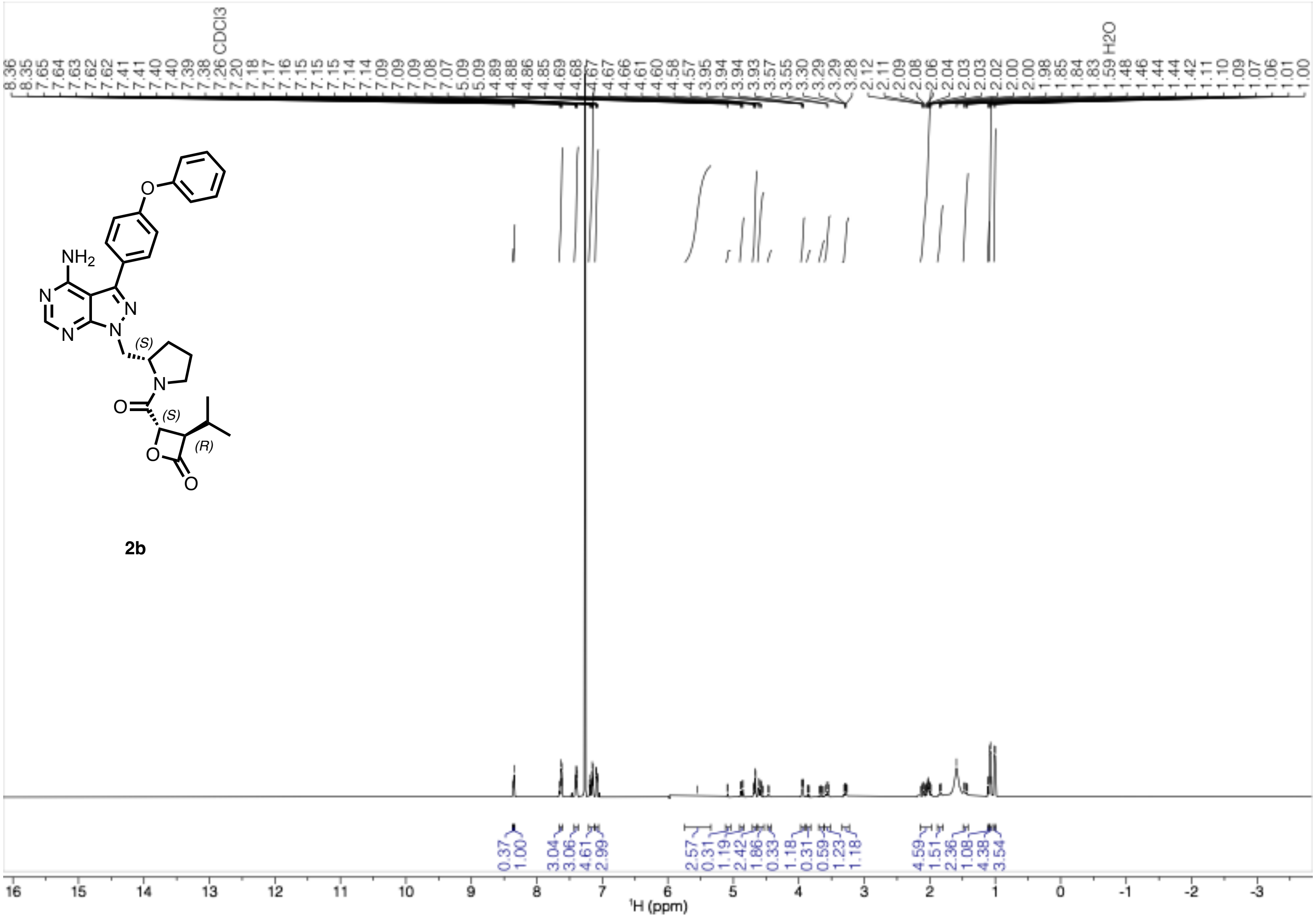

$^{1}$H-NMR (500 MHz, $CDCl_3$) of **3a**

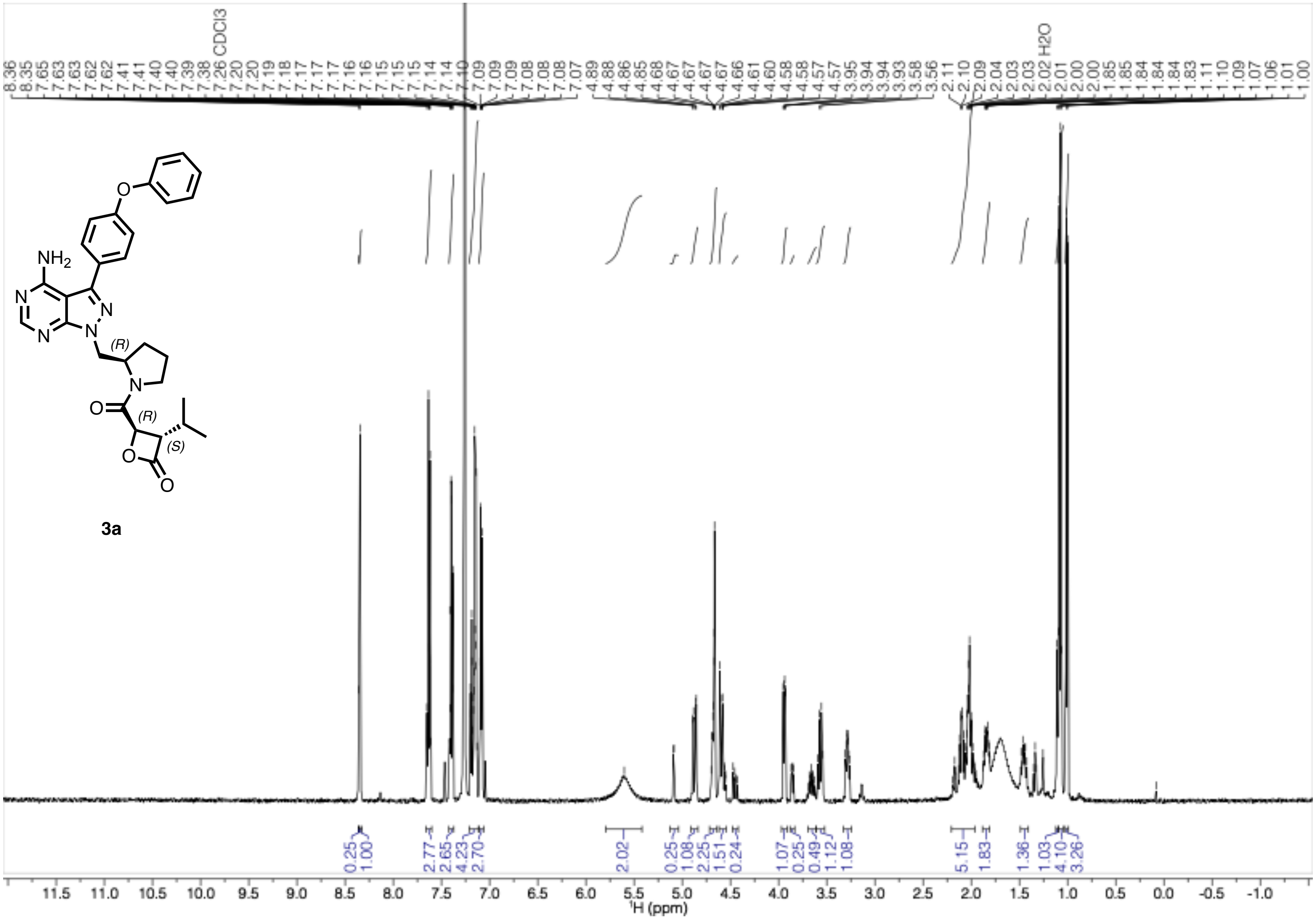

$^{1}H$-NMR (500 MHz, $CDCl_3$) of **3b**

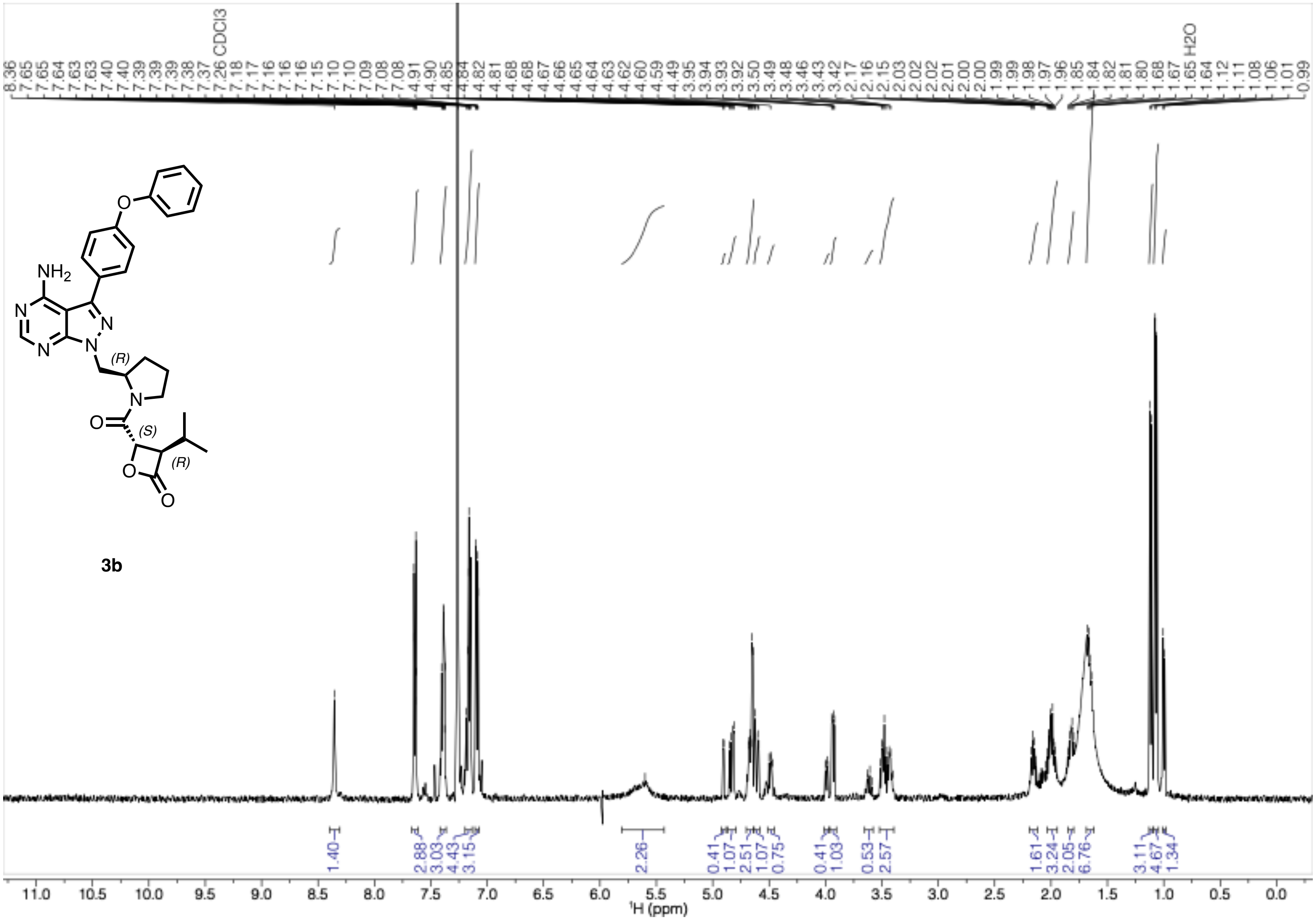

$^{1}$H-NMR (600 MHz, $CDCl_3$) of **4a**

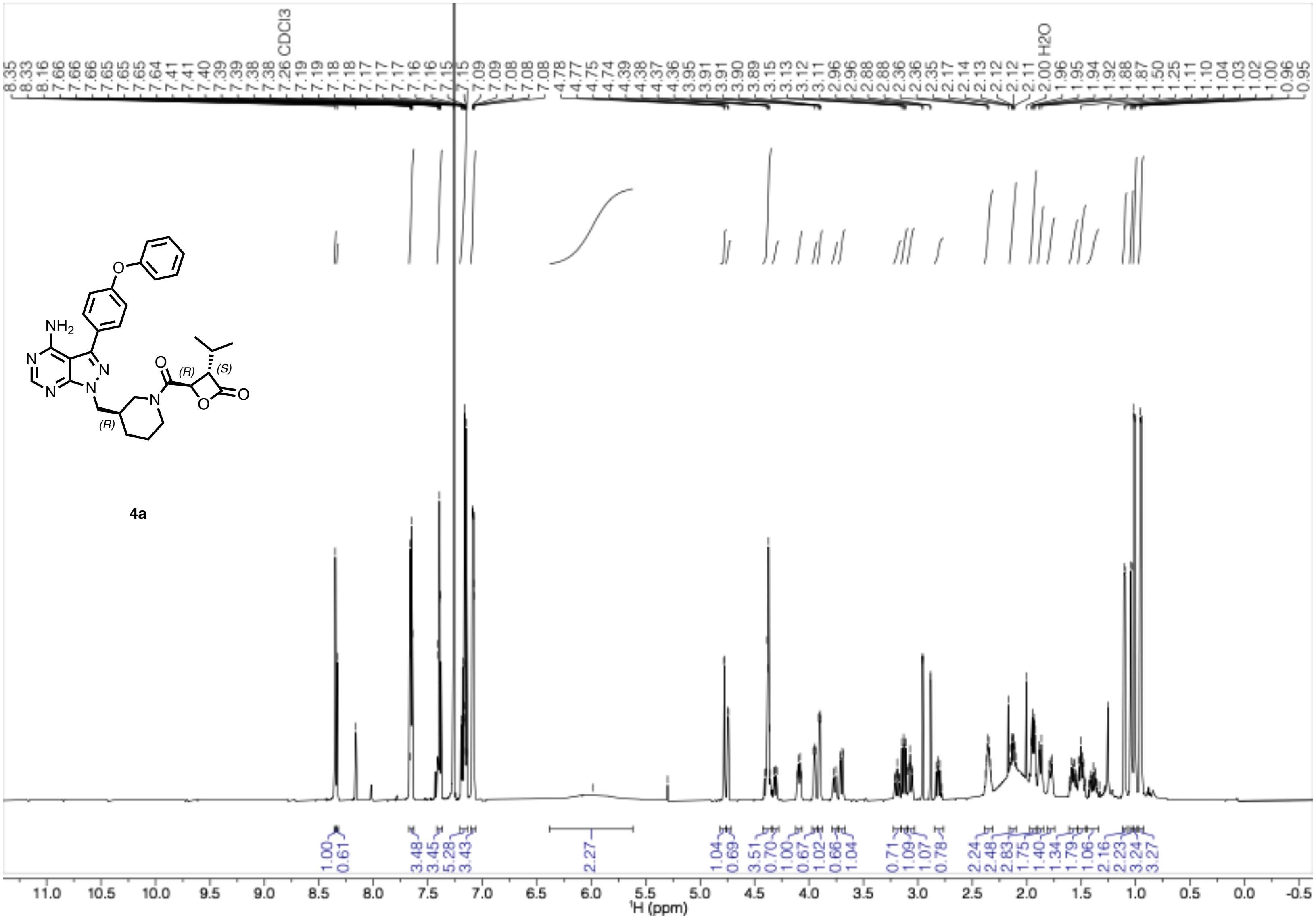

$^{1}H$-NMR (600 MHz, $CDCl_3$) of **4b**

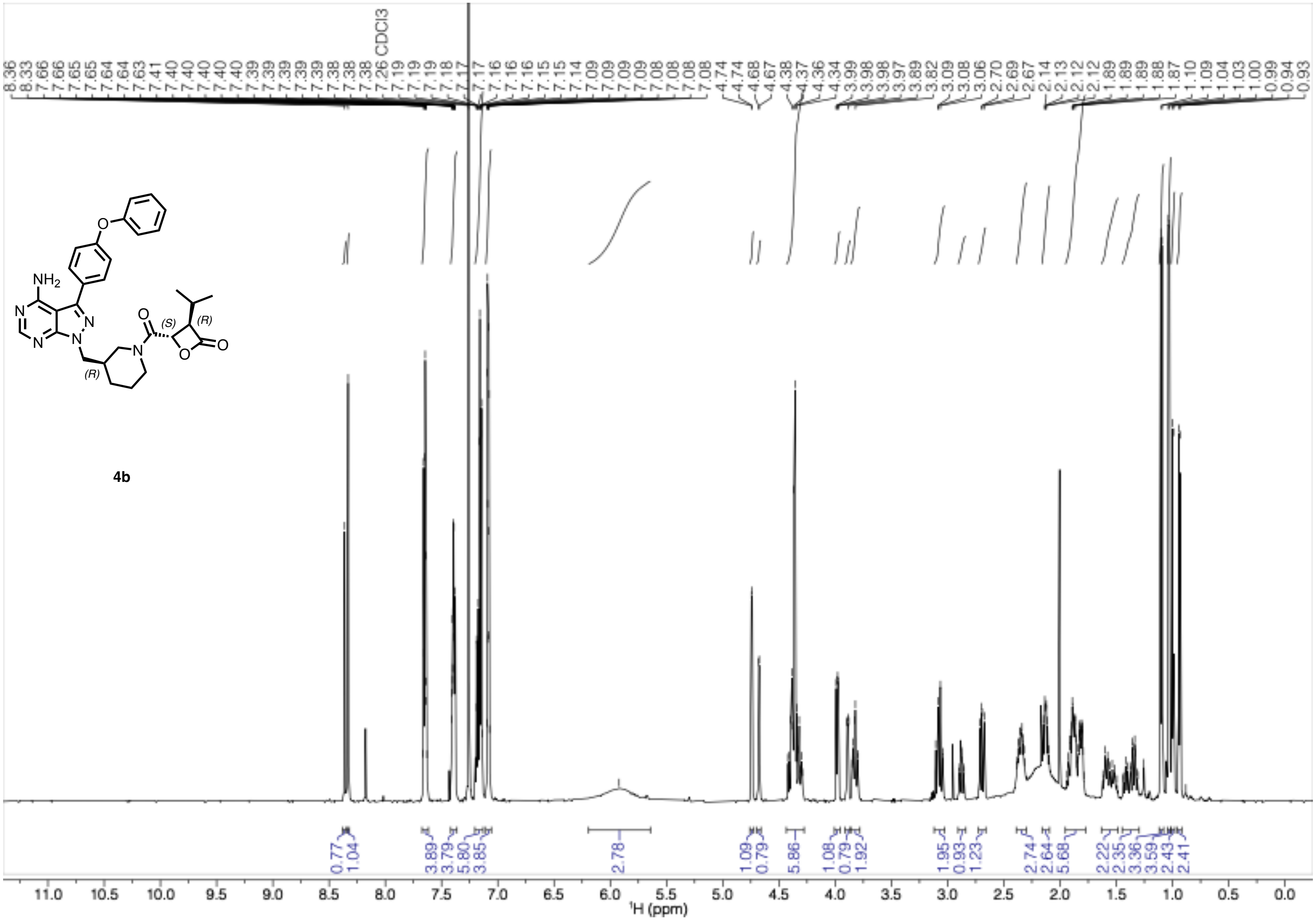

$^{1}$H-NMR (600 MHz, $CDCl_3$) of **5a**

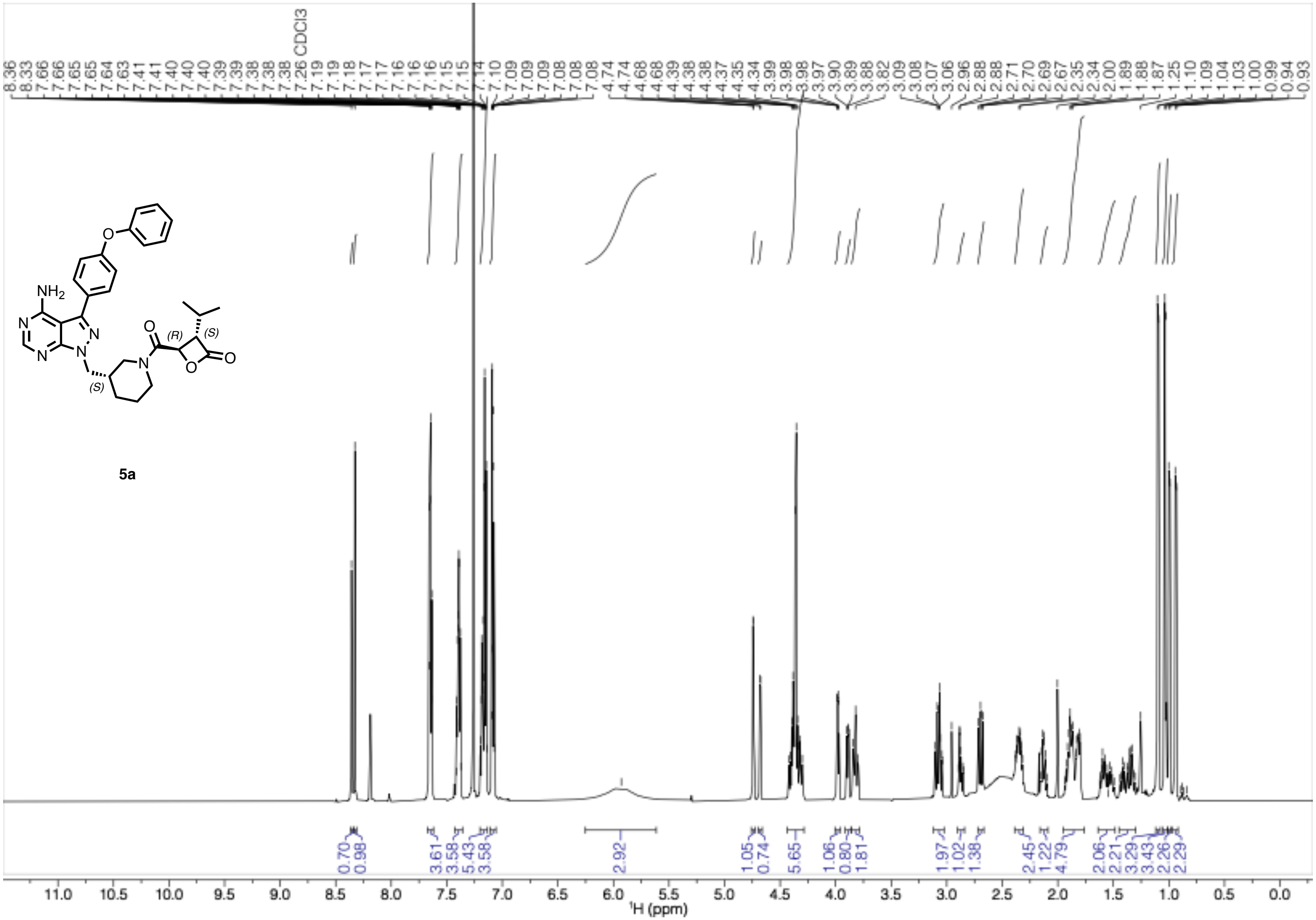

$^{1}$H-NMR (600 MHz, $CDCl_3$) of **5b**

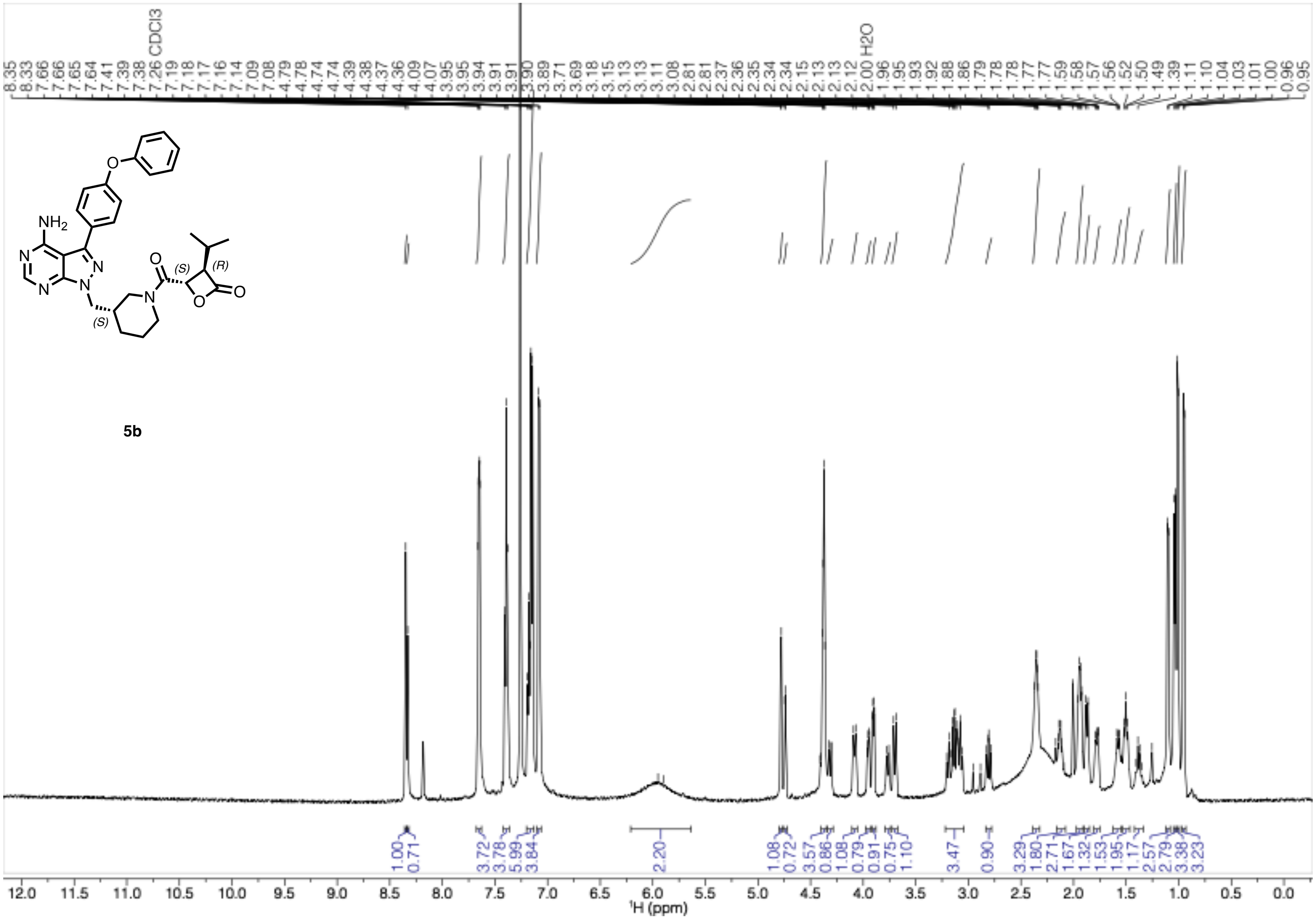

[1]H-NMR (600 MHz, $CDCl_3$) of **6**

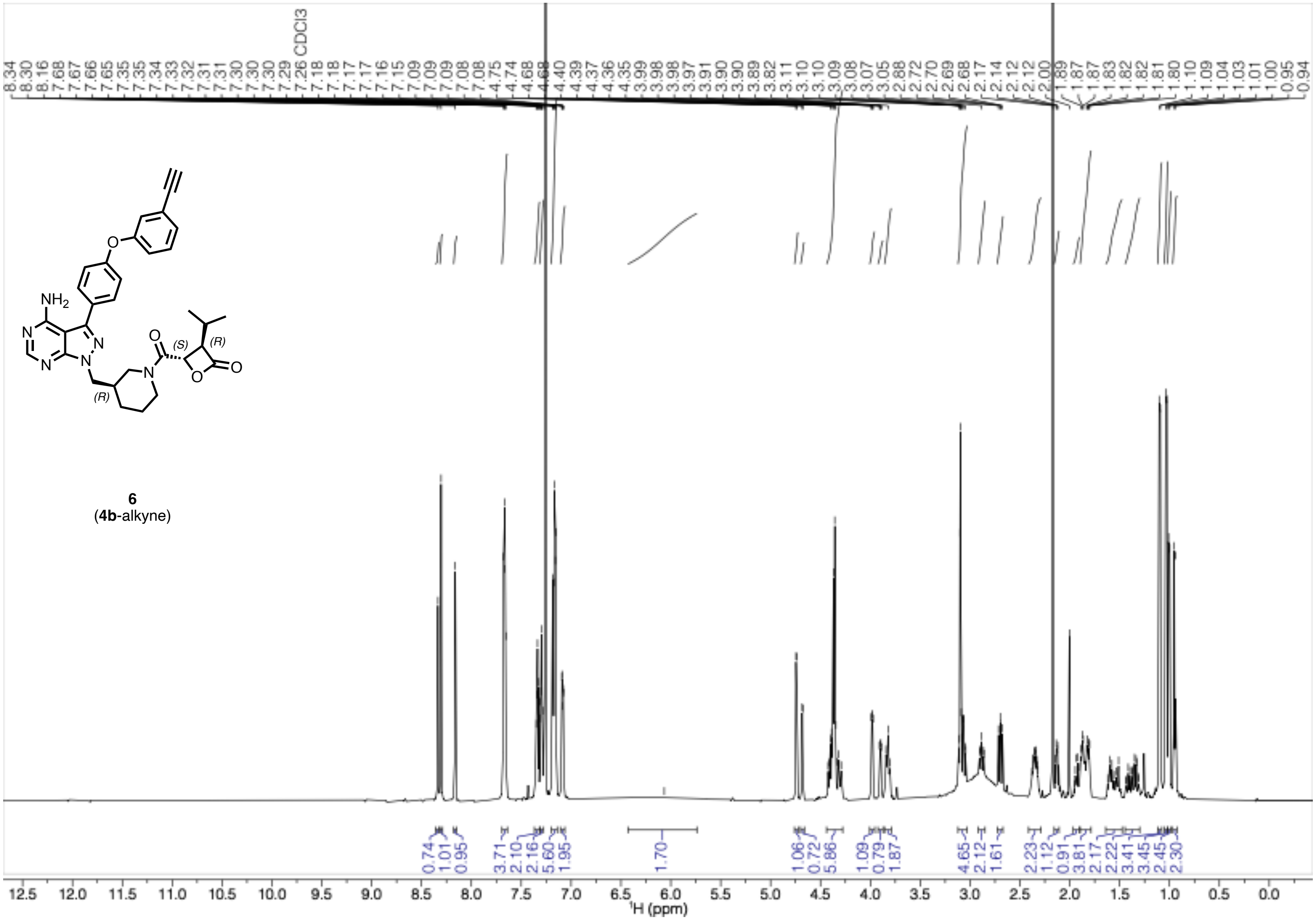

$^{1}$H-NMR (600 MHz, $CDCl_3$) of **7**

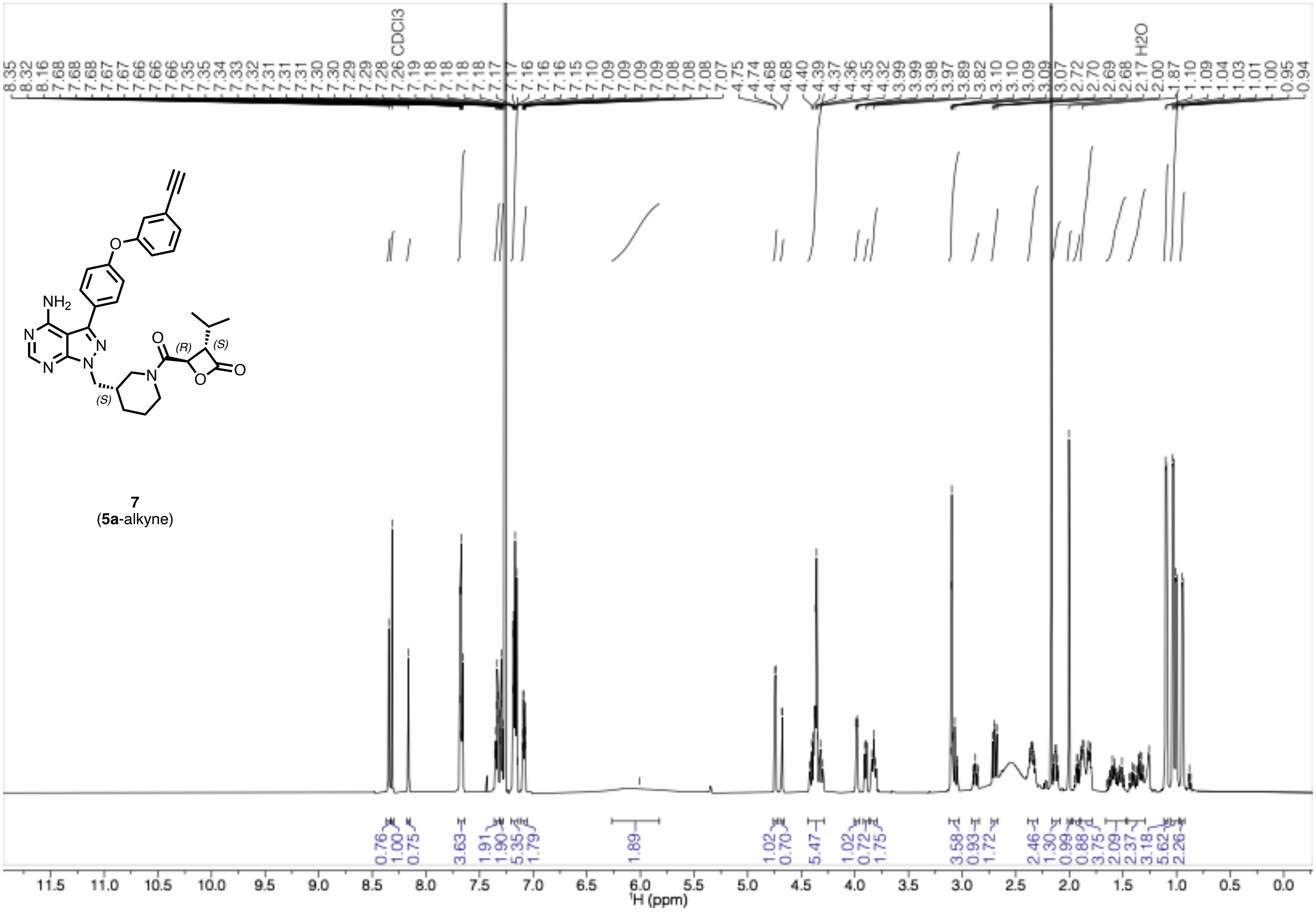

<sup></sup>[1]H-NMR (500 MHz, $CDCl_3$) of **9a**

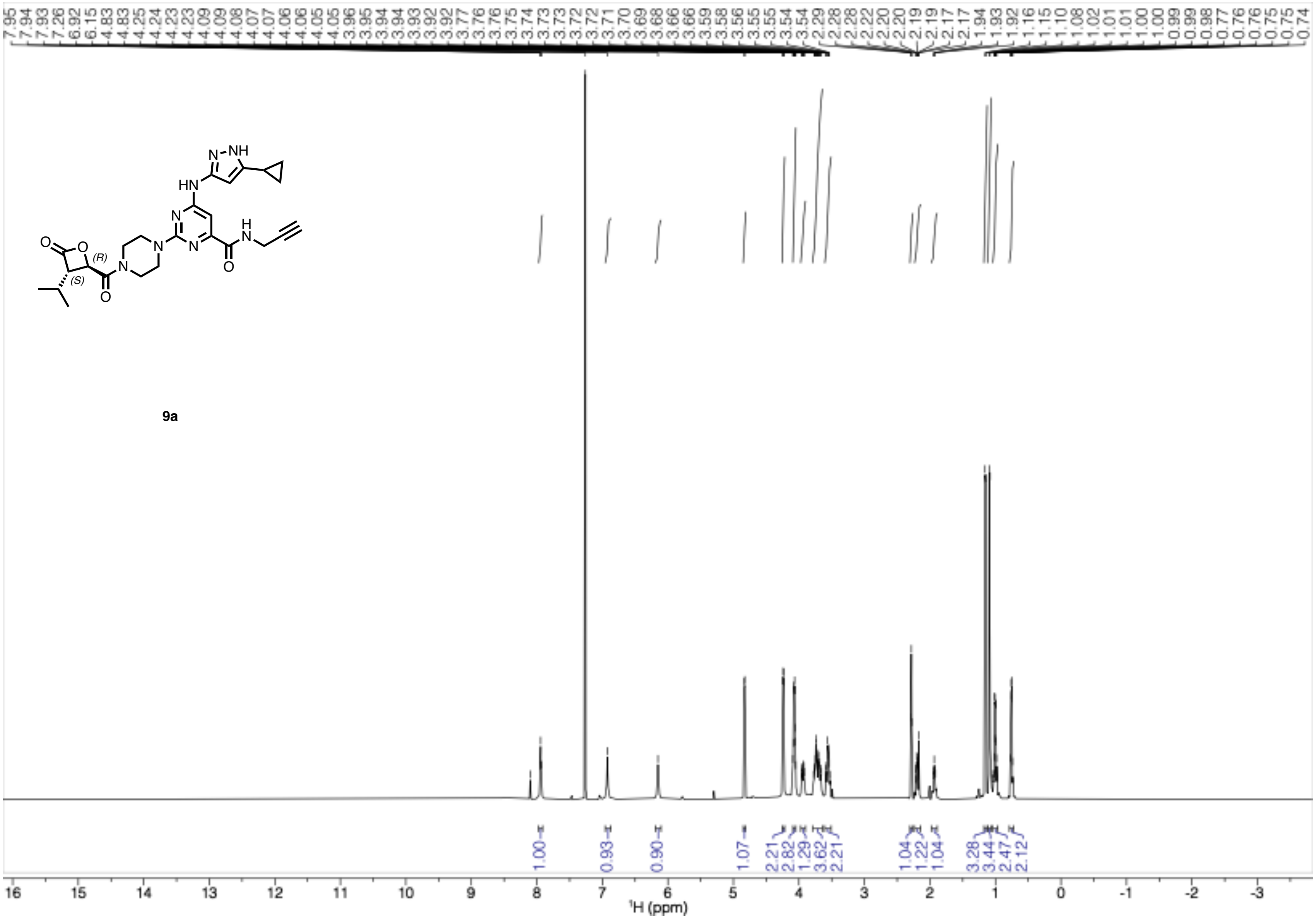

9a

[1]H-NMR (500 MHz, CDCl₃) of **9b**

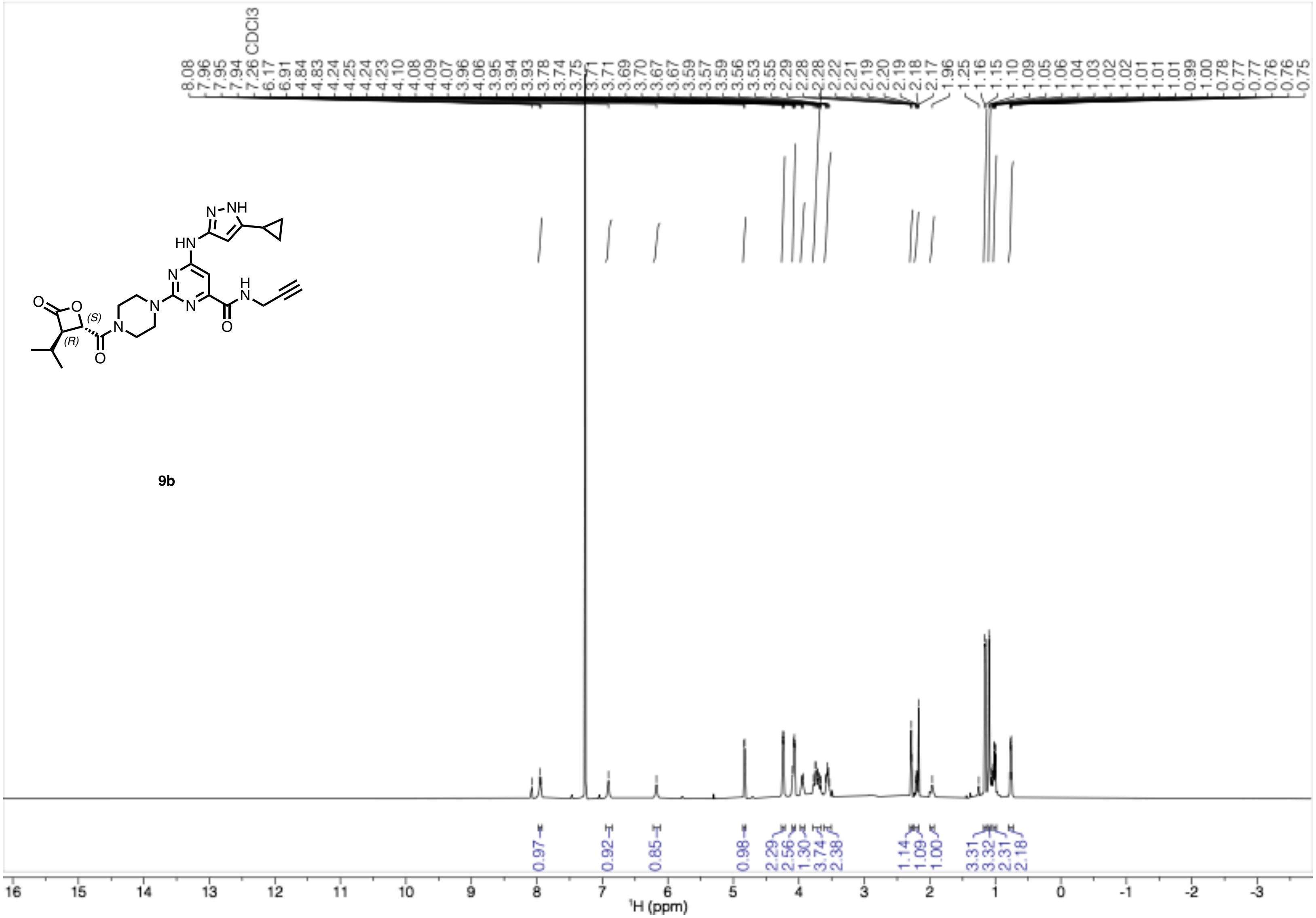

[1]H-NMR (500 MHz, CDCl3) of **11**

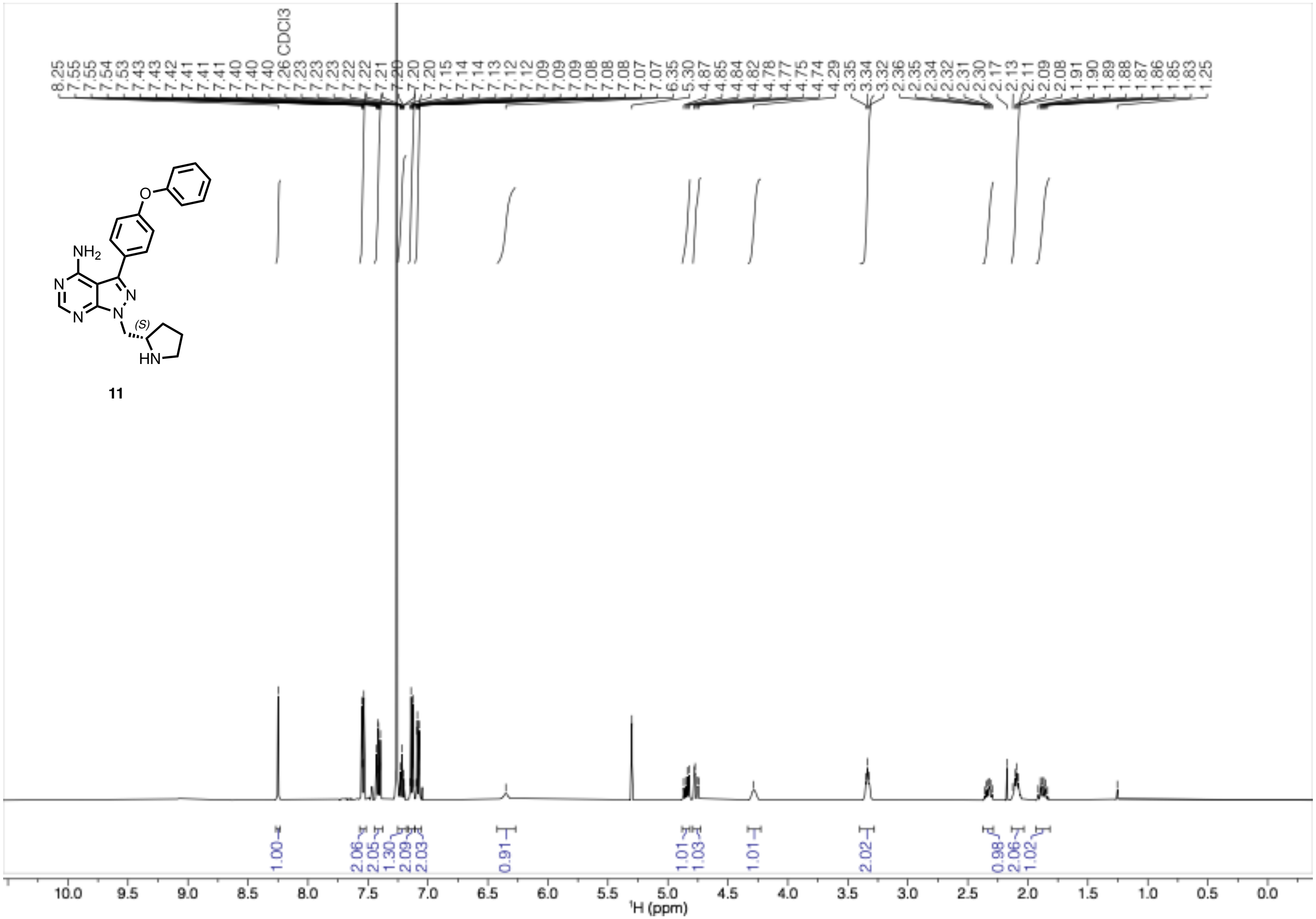

$^{1}$H-NMR (500 MHz, $CDCl_3$) of **12**

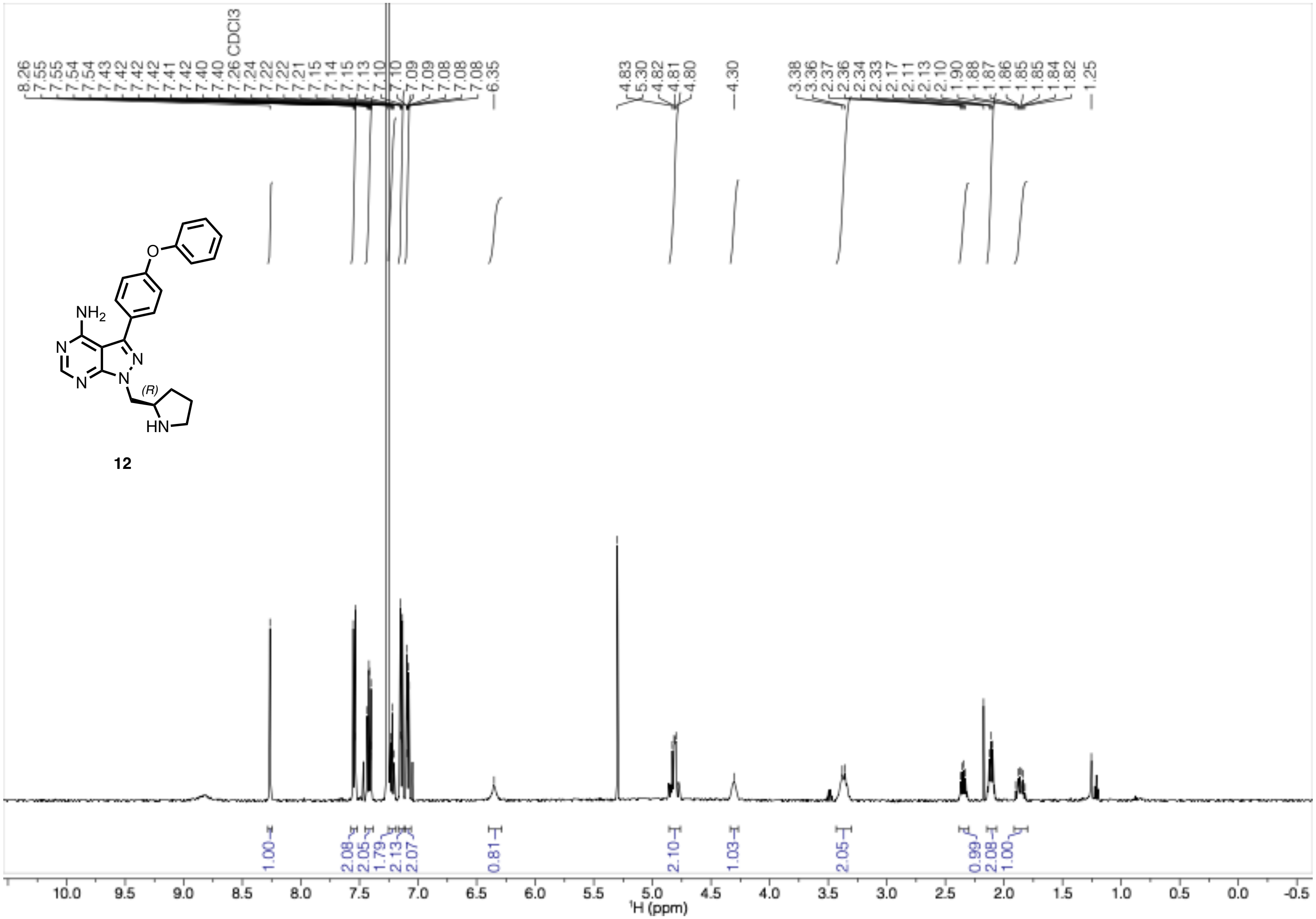

[1]H-NMR (500 MHz, $CDCl_3$) of **13**

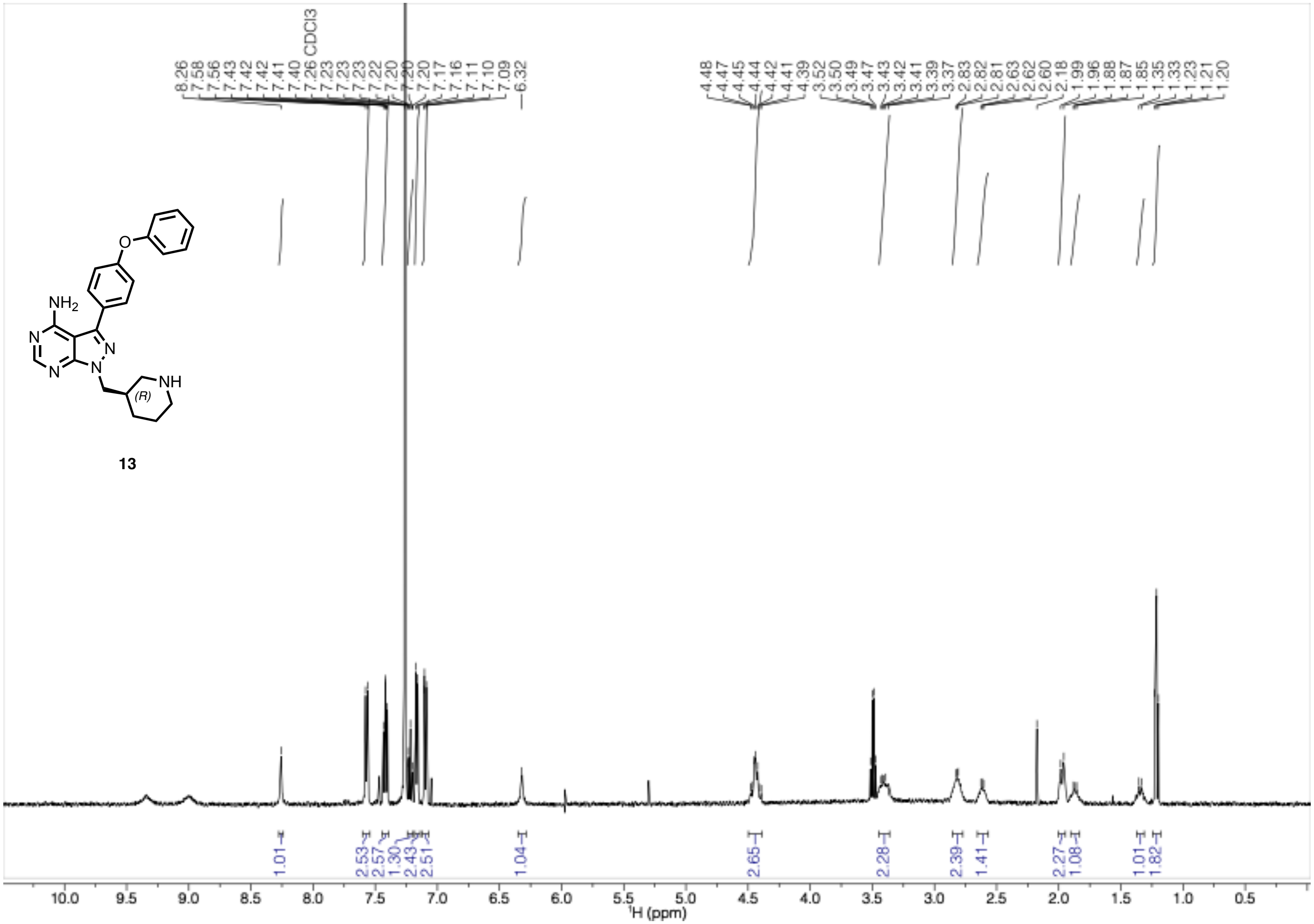

[1]H-NMR (500 MHz, $CDCl_3$) of **14**

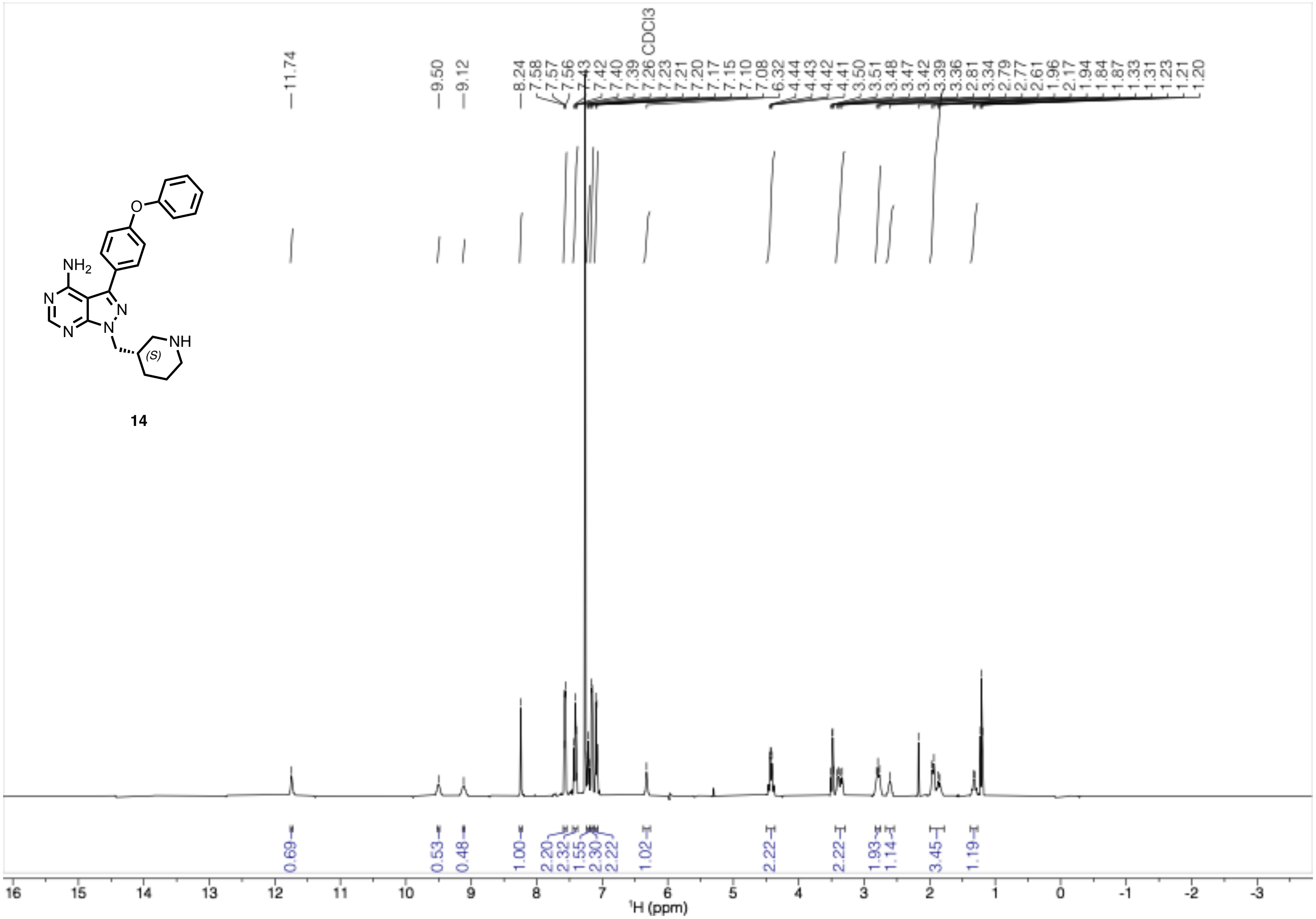

$^{1}$H-NMR (500 MHz, $CDCl_3$) of **15**

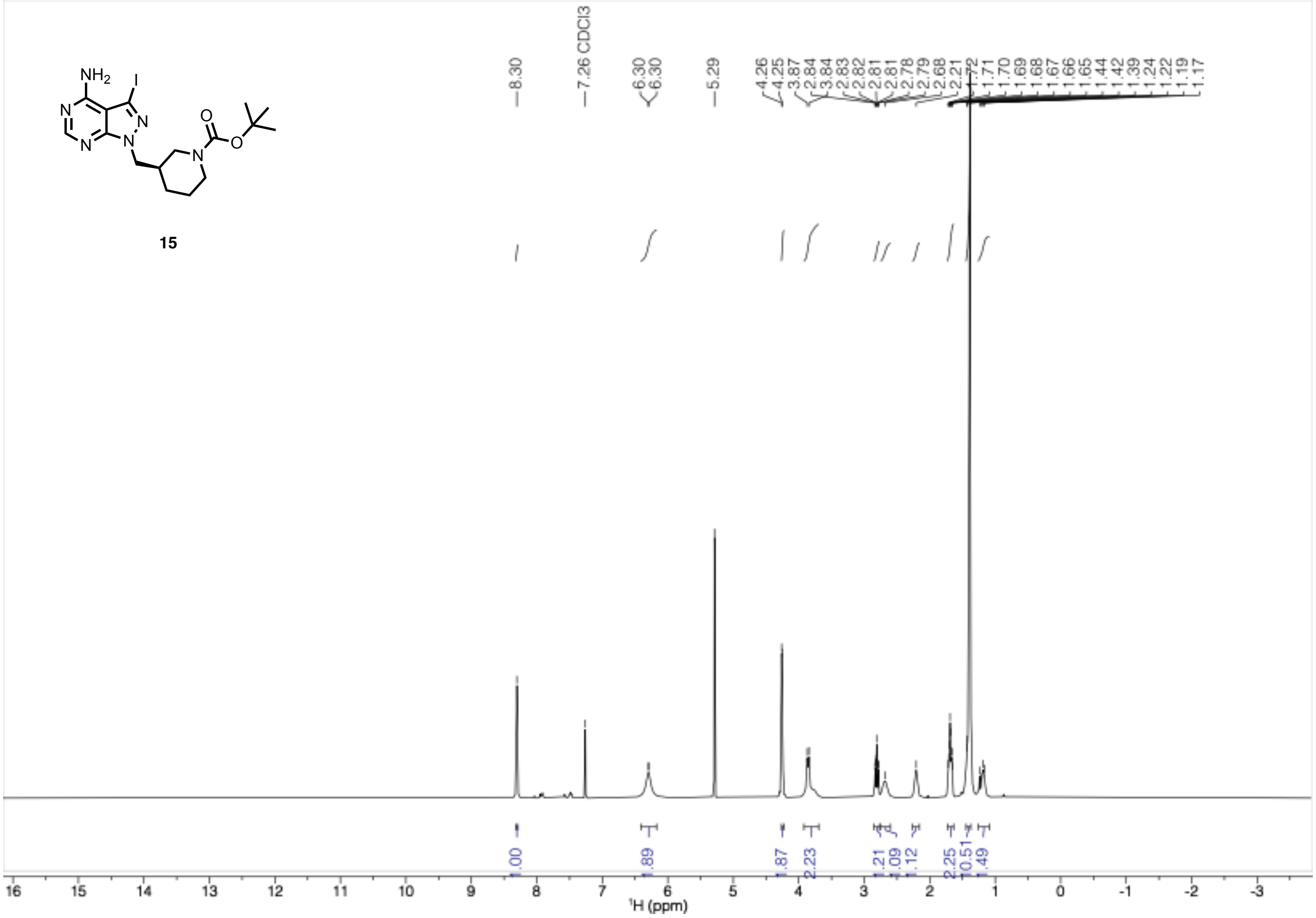

$^{1}$H-NMR (400 MHz, $CDCl_3$) of **16**

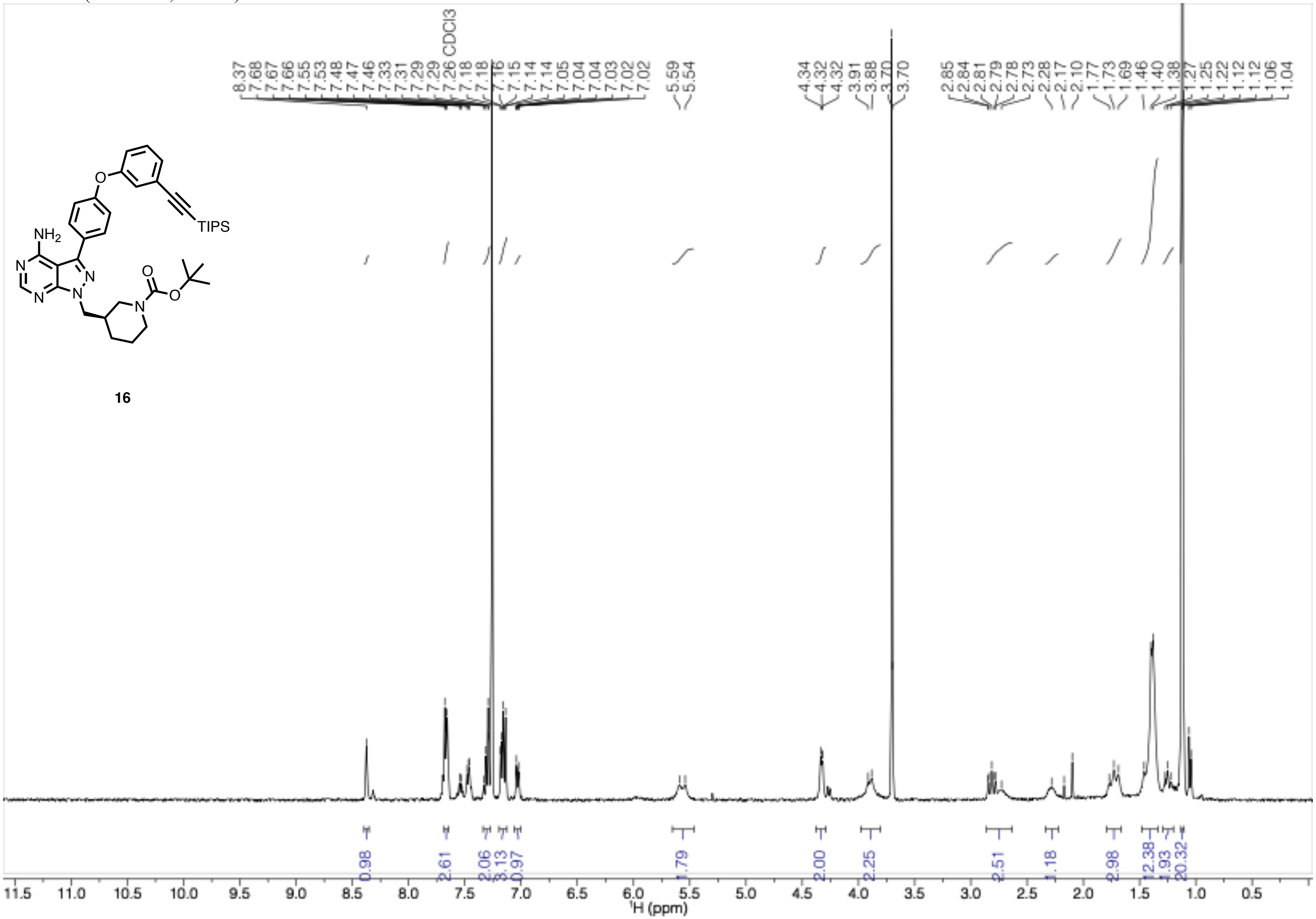

[1]H-NMR (500 MHz, $CDCl_3$) of **17**

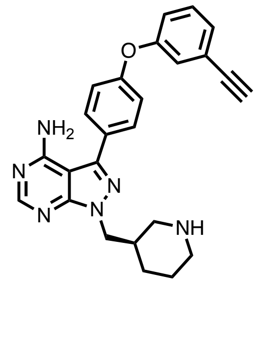

17

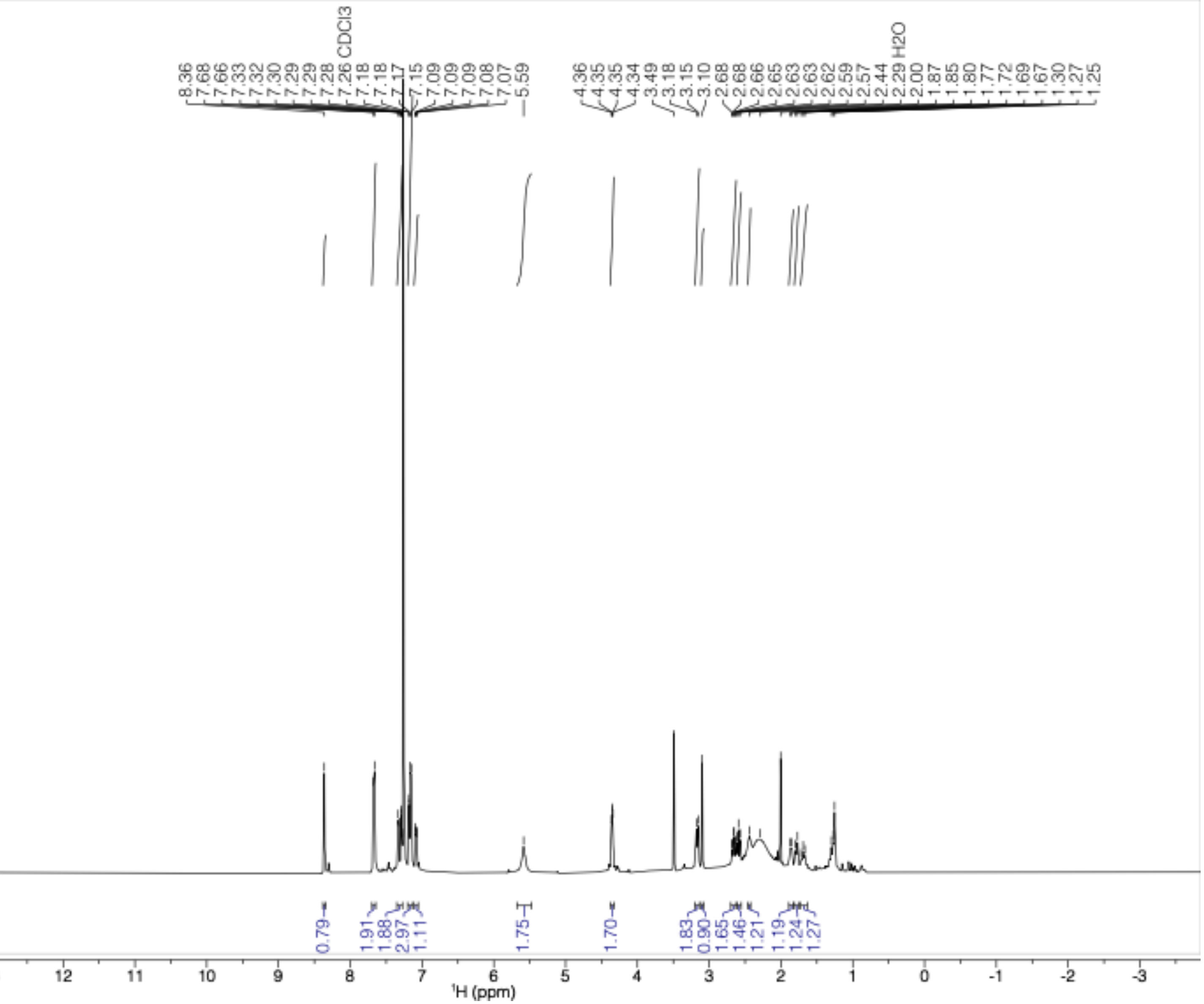

$^{1}$H-NMR (500 MHz, $CDCl_3$) of **18**

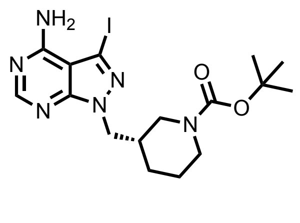

**18**

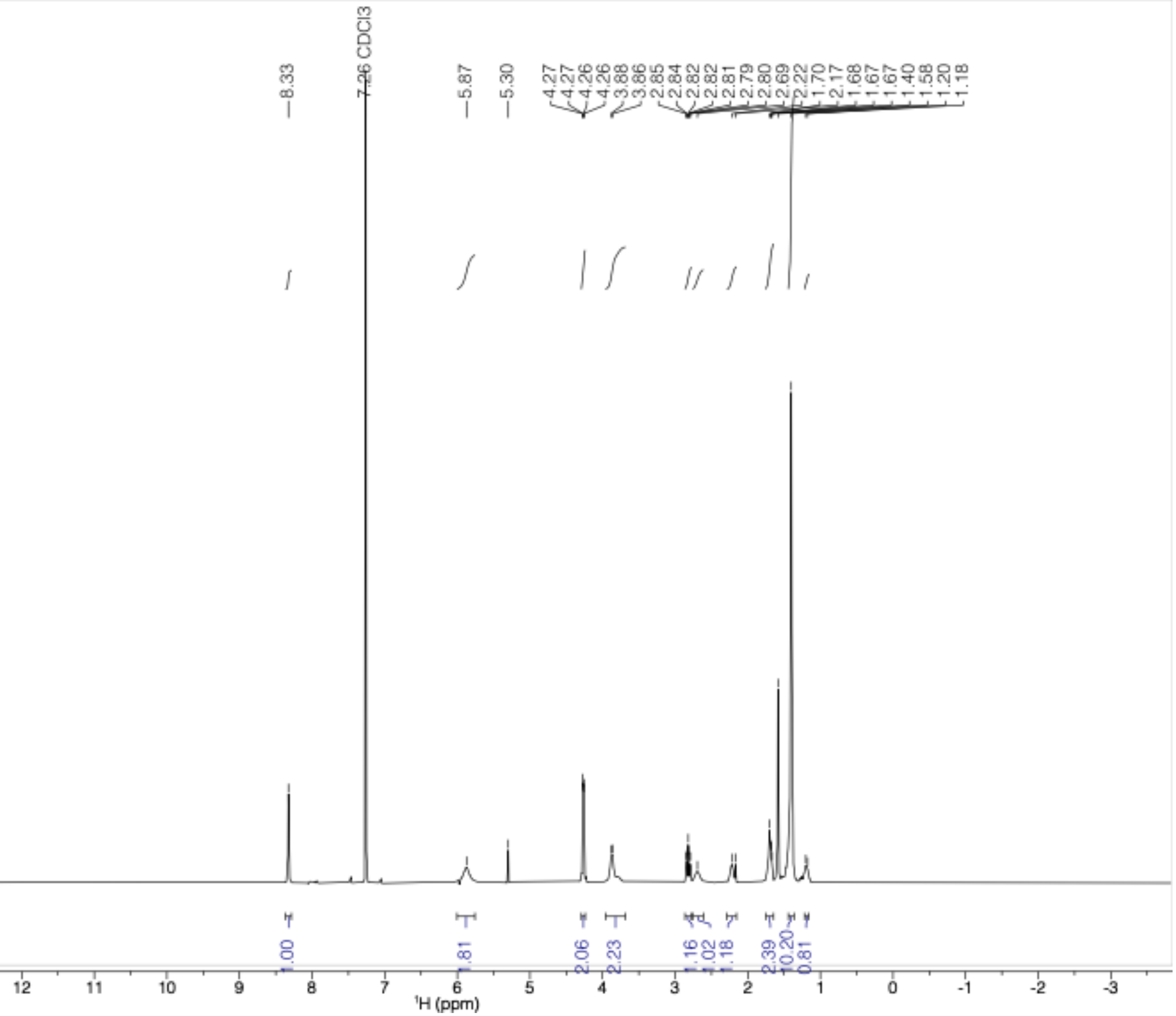

[1]H-NMR (400 MHz, $CDCl_3$) of **19**

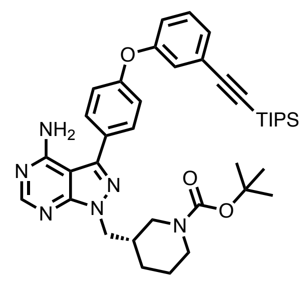

**19**

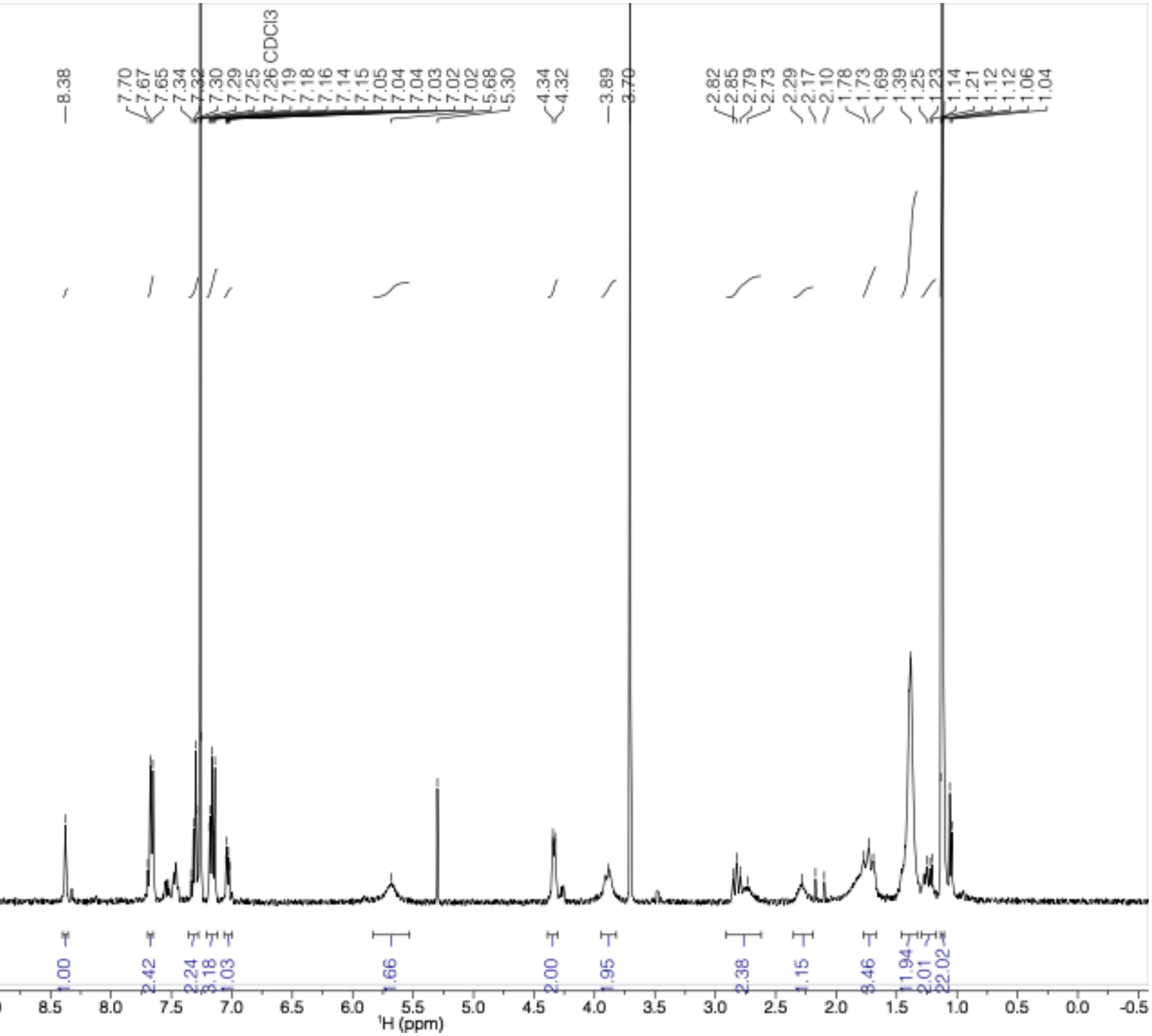

$^{1}$H-NMR (400 MHz, $CDCl_3$) of **20**

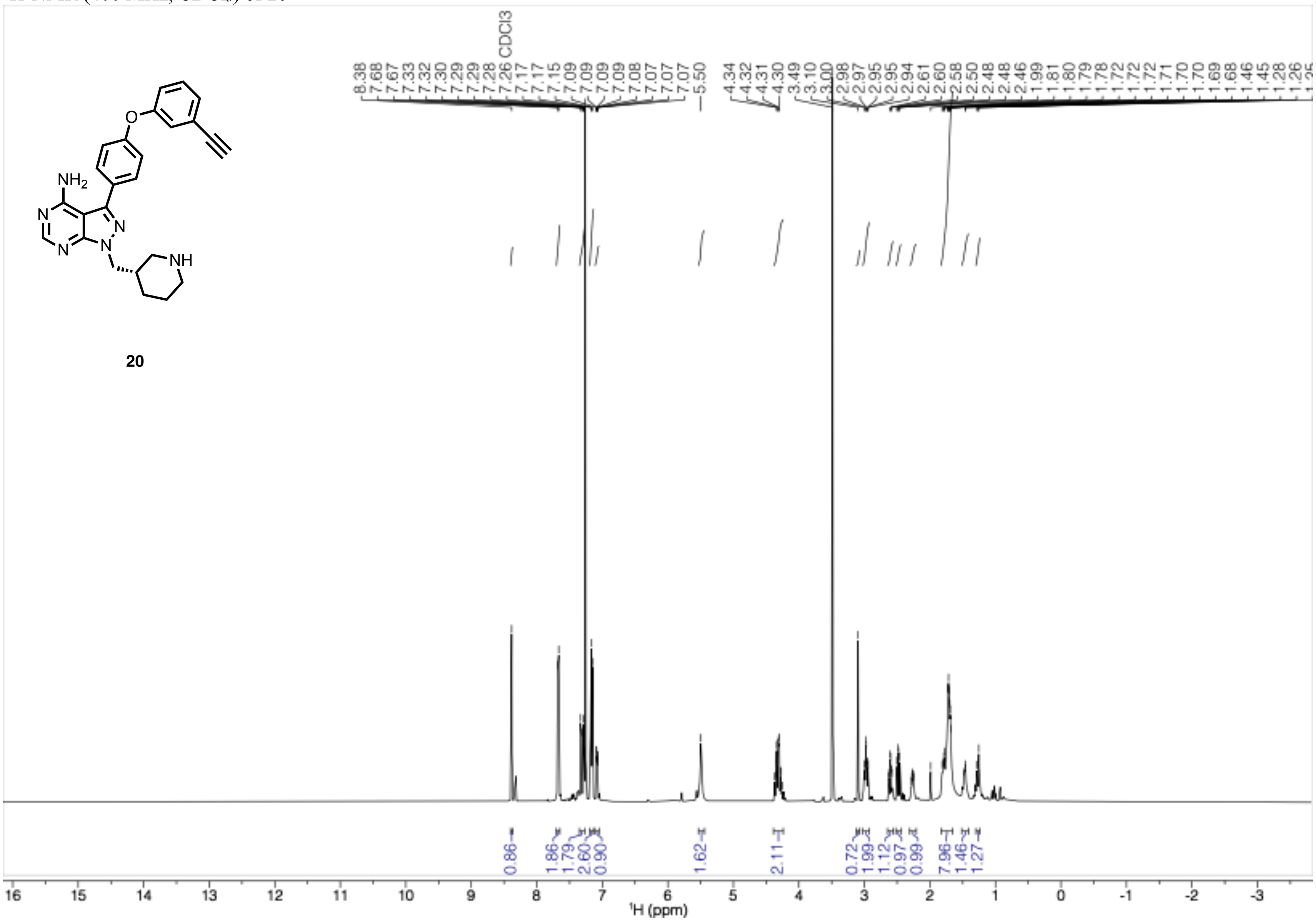
